# On the role of prefrontal and parietal cortices in mind wandering and dynamic thought

**DOI:** 10.1101/2023.11.07.566122

**Authors:** Tara Rasmussen, Hannah Filmer, Paul E. Dux

## Abstract

Mind wandering is a common phenomenon in our daily lives and can have both an adaptive and detrimental impact. Recently, a dynamic framework has been proposed to characterise the heterogeneity of internal thoughts, suggesting there are three distinct thought types which can change over time – *freely moving*, *deliberately constrained*, and *automatically constrained* (thoughts). There is very little evidence on how different types of dynamic thought map onto the brain. Previous research has applied non-invasive transcranial direct current stimulation (tDCS) to causally implicate the prefrontal cortex and inferior parietal lobule in general mind wandering. However, a more recently developed and nuanced technique, high-definition tDCS (HD-tDCS), delivers more focal stimulation able to target specific brain regions. Therefore, the current study investigated the effect of anodal HD-tDCS applied to the left prefrontal and right inferior parietal cortices (with the occipital cortex included as an active control) on mind wandering, and specifically, the causal neural substrates of the three internal dynamic thought types. This was a single session study using a novel task which allows investigation into how dynamic thoughts are associated with behavioural variability and the recruitment of executive control operations across the three brain regions. Anodal stimulation to the prefrontal cortex decreased freely moving thought and anodal stimulation to the parietal lobule decreased deliberately constrained thought, with preliminary evidence for an increase in freely moving thought in the occipital cortex as well. These findings support the heterogenous nature of mind wandering, revealing that different brain regions are implicated in distinct dynamic thought types.

---

We spend up to half our waking hours directing our thoughts towards self-generated, internally orientated representations: a phenomenon known as mind wandering (Kane et al., 2007; Seli, Beaty, et al., 2018; Smallwood & Schooler, 2006). Whilst mind wandering can be detrimental in contexts where a high level of sustained attention is required – such as air traffic control (Gouraud et al., 2017) and driving (Baldwin et al., 2017; Yanko & Spalek, 2014) – it has also been hypothesised that there are also associated benefits such as eliciting creativity (Baird et al., 2012) and facilitating future planning (Pachai et al., 2016). Given the broad range of impacts mind wandering has on our daily lives, and the extent to which we engage in it (Killingsworth & Gilbert, 2010), it is important to understand the neural, cognitive, and behavioural aspects which underly this phenomenon.

Mind wandering encompasses a broad range of experiences (Callard et al., 2013; Seli, Kane, et al., 2018), yet there has been limited exploration into the underlying complexities and heterogeneity of the associated internal thought processes (Kam et al., 2021; Martel et al., 2019). The “dynamic framework” has been proposed to account for this heterogeneity, suggesting that internal mental states arise and shift over time and can be distinguished as different thought types (Poerio et al., 2017; Seli, Kane, et al., 2018). There have been three key types of dynamic thoughts identified in the literature: 1) deliberately constrained thoughts, 2) automatically constrained thoughts and 3) freely moving thoughts (Christoff et al., 2016; Martel et al., 2019). Deliberately constrained thoughts require cognitive control and are directed towards goal-orientated information; automatically constrained thoughts focus largely on affective or sensory salient information which is difficult to disengage from and typically does not require cognitive control (Christoff et al., 2016; Irving, 2015; Seli et al., 2016); whereas, in comparison, freely moving thoughts arise without strong constraints or overarching direction and are not focused on particular topics for any period of time (Martel et al., 2019; Mills et al., 2018).

In terms of the neural substrates underlying mind wandering generally, an extensive literature has employed functional magnetic resonance imagining (fMRI) to implicate a large-scale collection of brain regions, known as the default mode network, in self-reported task unrelated thought (Christoff et al., 2009; Fox et al., 2015; Groot et al., 2021). Interestingly, this network shows increased metabolic activity when the mind is at rest and during internal thought processes, and decreased activity during cognitively demanding tasks (Mason et al., 2007; Poerio et al., 2017). The default mode network consists of a broad range of areas, with prominent regions including the medial prefrontal cortex (PFC), posterior cingulate cortex and posterior temporoparietal cortex (Fox et al., 2015; Groot et al., 2021). Researchers have also found mind wandering recruits the dorsolateral prefrontal cortex (DLPFC) and the anterior PFC, which are part of the frontoparietal control network (Christoff et al., 2009). These regions are associated with executive control processes, which hints at a negative correlation between executive functioning and mind wandering (Christoff et al., 2009; Fox et al., 2015). Executive functions are associated with actions including response selection, inhibition, attentional control, and multitasking (Diamond, 2013). These functions may be used to direct attention away from the current task, to allow the continuity of internal task unrelated thoughts and to guide and select between different types of dynamic thoughts (Christoff et al., 2016; Kam & Handy, 2014; Smallwood & Schooler, 2006).

Recently, cognitive neuroscientific investigations have begun to examine the neural processes associated with dynamic thought, with hints that different types may be linked with distinct neurocognitive operations (Martel et al., 2019; Wang et al., 2018). Initial research by Martel et al. (2019) used electroencephalogram (EEG) to investigate the neural correlates of the intentionality of task unrelated thought, reporting differences in the alpha oscillations and evoked sensory responses for deliberate and freely moving task unrelated thoughts. (Kam et al., 2021) conducted further research on this topic and observed the parietal event related potential P3 was greater for task-related compared to unrelated thoughts whereas the frontal P3 was larger for deliberate compared to automatic thoughts. This research provides correlational evidence that different types of dynamic thought may have distinct neural substrates, however, to date, there have been no attempts to causally implicate specific brain regions in dynamic thought.

One approach commonly used to investigate the casual neural substrates of mind wandering is transcranial direct current stimulation (tDCS). This is a form of non-invasive brain stimulation which passes low-intensity currents (typically 0.5mA to 3mA) between electrodes attached to the scalp (see Figure 1; Filmer et al., 2014, 2020; Mosayebi Samani et al., 2019). The effects of tDCS on modulating resting membrane potentials were initially thought to be polarity dependent, with anodal stimulation increasing the likelihood of cortical excitation and cathodal stimulation decreasing the likelihood of cortical excitation (Nitsche & Paulus, 2000). However, recent research suggests that the polarity of tDCS effects is far more complex and is affected by factors such as stimulation intensity, duration, and task type (Filmer et al., 2019, 2020; Mosayebi Samani et al., 2019). While the field of tDCS has faced criticism over variability in the outcomes and the use of poor methodological practices (Horvath et al., 2016), many of these criticisms have more recently been addressed. Specifically concerns about the reproducibility of tDCS have been discussed in a more recent review by Filmer et al. (2020), which identified various fields with consistent findings across studies, alongside highlighting the importance of methodological and scientific rigour for developing reproducible findings using tDCS. Thus, this form of non-invasive brain stimulation offers novel insights into our understanding of the brain and has many potential benefits within the field of cognitive neuroscience and for society (Filmer et al., 2020).

**Figure 1.**
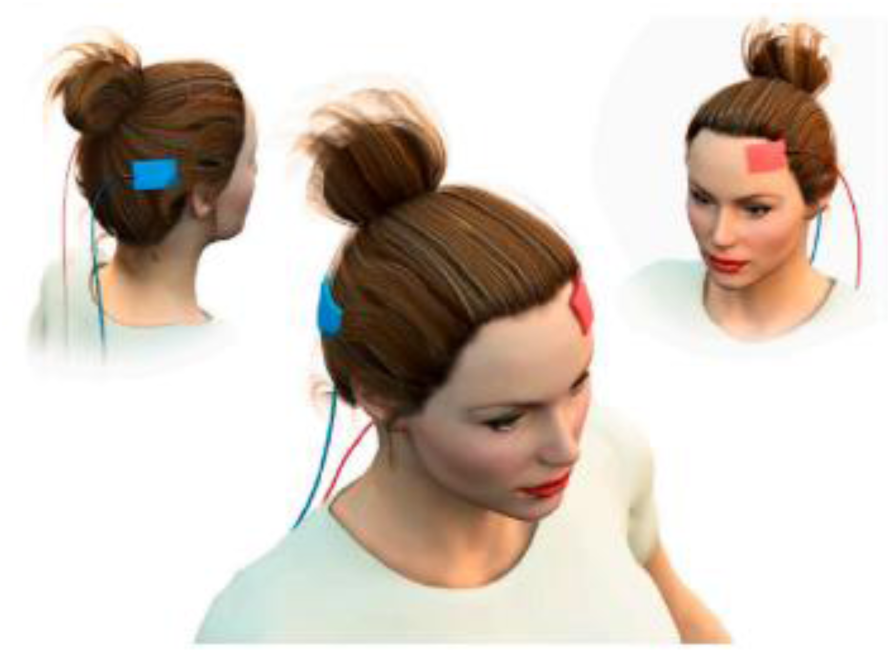
An illustration of a bipolar tDCS montage showing the target electrode (red) over the prefrontal cortex (F3) and reference electrode (blue) over the parietal cortex (P4).

There have been several studies investigating the effect of bipolar tDCS on mind wandering in the prefrontal cortex (Axelrod et al., 2015, 2018; Filmer et al., 2019). Initial studies observed that low intensity stimulation, specifically 1mA anodal tDCS applied to the left DLPFC (but was likely more broadly the left PFC given the bipolar tDCS montage, see Figure 1), increased participants mind wandering propensity (Axelrod et al., 2015, 2018). However, Boayue et al. (2020) completed a high-powered replication of Axelrod et al. (2015) and found strong evidence for a failure to replicate the effect of anodal stimulation over the left PFC. Bertossi et al. (2017) also investigated mind wandering and the medial PFC and found a decrease in the propensity to mind wander following cathodal stimulation, however the effect was only found in male subjects. Given evidence hinting at the importance of tDCS intensity, a recent large scale pre-registered study explored the effect of this variable on mind wandering propensity (Filmer et al., 2019). This study targeted the left PFC, applying 1 or 2mA of anodal or cathodal stimulation. The experiment found that only 2mA cathodal stimulation to the left PFC influenced mind wandering, such that the frequency of task unrelated thoughts was increased (Filmer et al., 2019). The inconsistencies in findings and failure to replicate early results has led to questions over the causal role of the prefrontal cortex in mind wandering and the mechanisms which may explain these underlying differences.

An additional region which has been implicated in modulating mind wandering propensity is the right inferior parietal lobule (IPL; Filmer et al., 2021; Kajimura et al., 2019; Kajimura & Nomura, 2015). Research applying 1.5mA anodal tDCS to the right IPL found stimulation reduced the propensity to mind wander (Kajimura et al., 2019; Kajimura & Nomura, 2015). Interestingly, Kajimura et al. (2019) uniquely stimulated the IPL region, finding the effect of anodal tDCS on mind wandering was specific to the right IPL alone, as opposed to finding effects in the IPL while concurrently stimulating the PFC. Filmer et al. (2021) investigated the role of both the PFC and IPL in mind wandering using 1mA and 2mA stimulation intensities and inversed polarity montages. These researchers found that anodal stimulation to the left PFC and cathodal stimulation to right IPL increased mind wandering propensity (Filmer et al., 2021). This research aligns with previous literature which, collectively, provides evidence for a causal role of both the PFC and IPL in mind wandering. However, these results display inconsistencies in the directionality of the effects and the interplay between the different types of dynamic thought and activity in these regions is still unknown (Chaieb et al., 2019).

There is debate that the effects of traditional bipolar tDCS are diffuse, with the electrodes stimulating relatively broad brain regions and thus not targeting a focal cortical area (Datta et al., 2009; Pisoni et al., 2018). To address this issue, a newly developed approach - high definition tDCS (HD-tDCS) - utilises multiple smaller electrodes (typically a 4 x 1 ring montage) to deliver more targeted stimulation (Nikolin et al., 2019; Villamar et al., 2013). As seen in Figure 2, the electrodes used in the HD-tDCS configuration cover a smaller cortical area than the larger electrodes used for bipolar tDCS, and are also more focal, with the 4×1 ring configuration designed to restrict the area of cortical excitability modulation to the region within the ring perimeter (Villamar et al., 2013).

**Figure 2.**
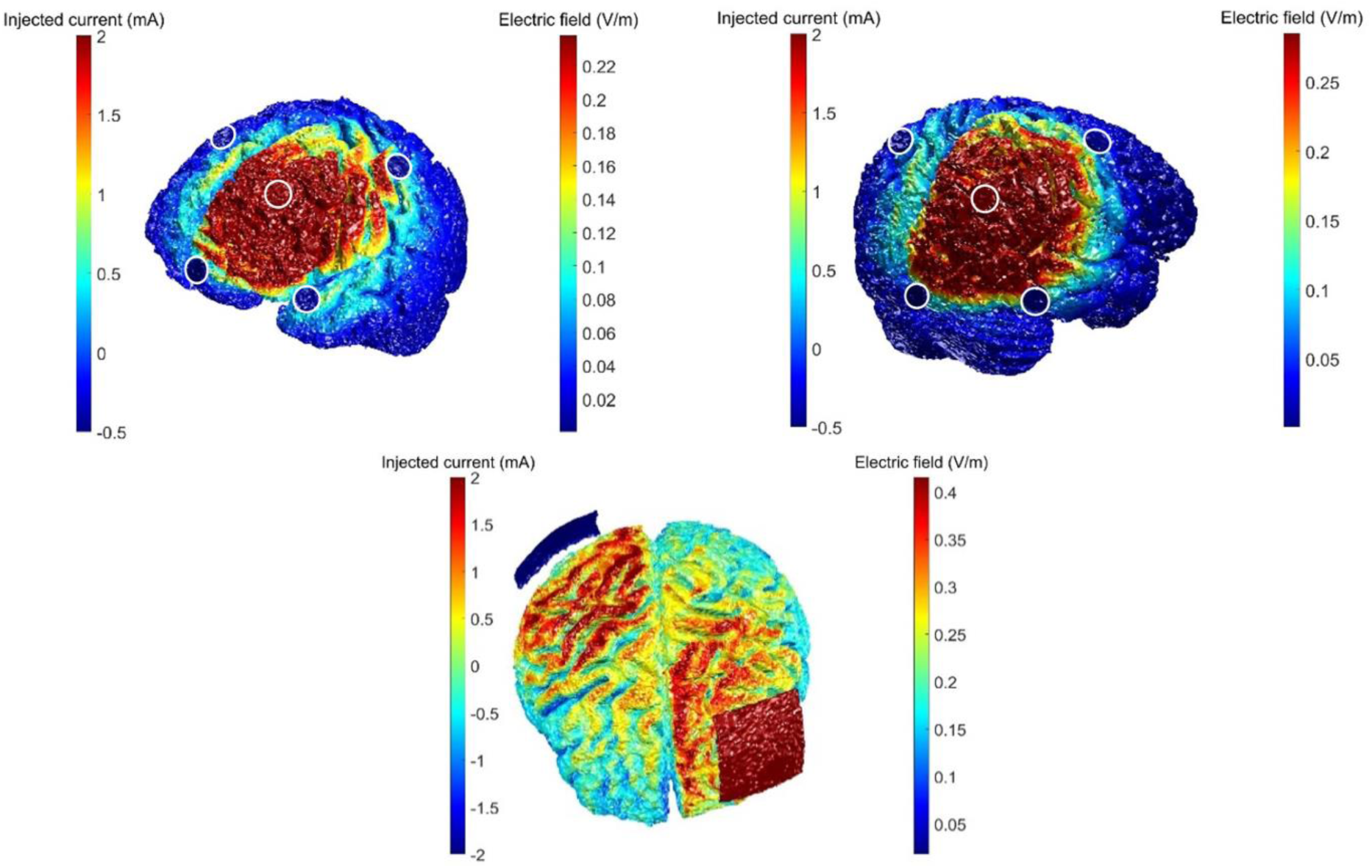
Current modelling comparing the 4 x 1 ring HD-tDCS montage over the left prefrontal cortex (on the upper left), with the anode at F3 and cathodes placed at F7, C3, Fz and Fp1 and the right parietal cortex (on the upper right), with the anode at P4 and cathodes placed at T6, O2, Pz and C4. This is compared to a bipolar tDCS montage (lower image), with the active electrode over the left prefrontal cortex and reference electrode positioned over the right parietal cortex.

A recent study conducted by Chou et al. (2020) investigated the effects of bilateral HD-tDCS to the posterior IPL on maladaptive forms of mind wandering. This study manipulated participants mood and probed them on the contents of their thoughts in between completing a multi-source inference task. The study utilised a double 3 x 1 electrode montage, applying 1mA to both the right and left IPL and included an anodal, cathodal, and sham group. Interestingly, this study found no effect of either anodal or cathodal stimulation on overall mind wandering frequency. However, cathodal stimulation was found to significantly decreased the frequency of negative mind-wandering thoughts about the past relative to sham. These findings offer initial support for distinctions in the causal brain regions of internal thought types; however, the research focused on negatively valanced thoughts for a single brain region.

An additional large-scale study conducted by Boayue et al. (2021) is the only study to employ HD-tDCS to the DLPFC to investigate the causal neural substrates of mind wandering. This study applied 2mA anodal HD-tDCS to the left DLPFC and tested performance on a novel task which consisted of generating random sequences in time with a metronome (Boayue et al., 2021). Throughout the task a thought-probe was presented asking participants to rate how focused they were on the task using a 4-point Likert scale. These researchers employed hierarchical order probit modelling and found anodal stimulation decreased the propensity to mind wander relative to sham. They argued this analysis technique improved sensitivity as the thought probe data is treated as an ordinal variable and assessed against a number of predictors including measures of task performance and the trial number (Boayue et al., 2020, 2021). This research provides preliminary evidence for the feasibility of anodal HD-tDCS modulating self-reported mind wandering in the PFC, using a simple thought probe measuring participants focus on the task (Boayue et al., 2021). However, HD-tDCS has not yet been applied to understand the causal neural substrates of different types of dynamic thought across brain regions.

## The Present Study

Considering the importance of mind wandering for both theoretical and applied settings it is crucial to understand the complex heterogeneity of dynamic thoughts. The current research employed a similar protocol to Boayue et al. (2021), however anodal HD-tDCS was used to explore the causal neural substrates of the dynamic thought types across three brain regions. Thus, our study aimed to use HD-tDCS to investigate whether task unrelated, freely moving, deliberately constrained, and automatically constrained thoughts were associated with distinct causal neural substrates, specifically in the PFC and IPL, with the occipital cortex being included as an active control region. We employed a double-blind protocol with six equal groups of participants who received 2mA anodal or sham HD-tDCS. The study also aimed to understand the effect of HD-tDCS on the frequency of different dynamic thoughts and task performance across these brain regions.

We predicted that there would be differences in the frequency of task unrelated thoughts between the active and sham HD-tDCS groups, consistent with the effect found by Boayue et al. (2021). Specifically, active anodal stimulation to the left PFC and right IPL would decrease the frequency of task unrelated thoughts, compared to sham and the occipital cortex (H_1_). Furthermore, we hypothesised that there would be a decrease in the frequency of the novel types of dynamic thought (i.e., freely moving, deliberately constrained, and automatically constrained thoughts) for groups receiving anodal HD-tDCS to the left PFC and the right IPL, relative to sham and the occipital cortex (H_2_). Finally, we hypothesised that HD-tDCS would affect task performance, such that active stimulation to the left PFC and right IPL would increase participants randomness (approximate entropy) and reduce behavioural variability, relative to sham and the occipital cortex (H_3_). Consistent with Boayue et al. (2021), we also predicted there would be a relationship between behavioural variability and approximate entropy.

## Method

### Overview

All participants completed one session, which consisted of HD-tDCS to one of three brain regions in conjunction with the Finger-Tapping Random-Sequence Generation Task (FT-RSGT). This task was designed for participants to generate random number sequences while listening to a repetitive metronome. Participants were pseudo randomly assigned to one of six equal groups, with half the participants for each brain region receiving active anodal HD-tDCS and the other half receiving sham HD-tDCS. The specific groups were: (1) anodal HD-tDCS over the left PFC; (2) sham HD-tDCS over the left PFC; (3) anodal HD-tDCS over the right IPL; (4) sham HD-tDCS over the right IPL; (5) anodal HD-tDCS over the occipital cortex (VC); and (6) sham HD-tDCS over the VC. At the beginning of the session, participants completed a questionnaire assessing their experience with musical instruments and video games. They were then taught about how to generate random sequences with a short practice block; given explanations on the four types of thought probes which will be included in the experiment and, following this, tested on their ability to identify each dynamic thought type, alongside completing a short training block on the task, and thought probes combined. All participants then completed a baseline block of the FT-RSGT, followed by 30mins of online active or sham stimulation in conjunction with the FT-RSGT. After concluding the experiment, participants completed an end of session questionnaire and were debriefed on their involvement in the study. See Figure 3 for the full outline of the experimental procedure. The full stage one registered report manuscript can also be accessed on the OSF (https://doi.org/10.17605/OSF.IO/XZ4RN).

**Figure 3.**
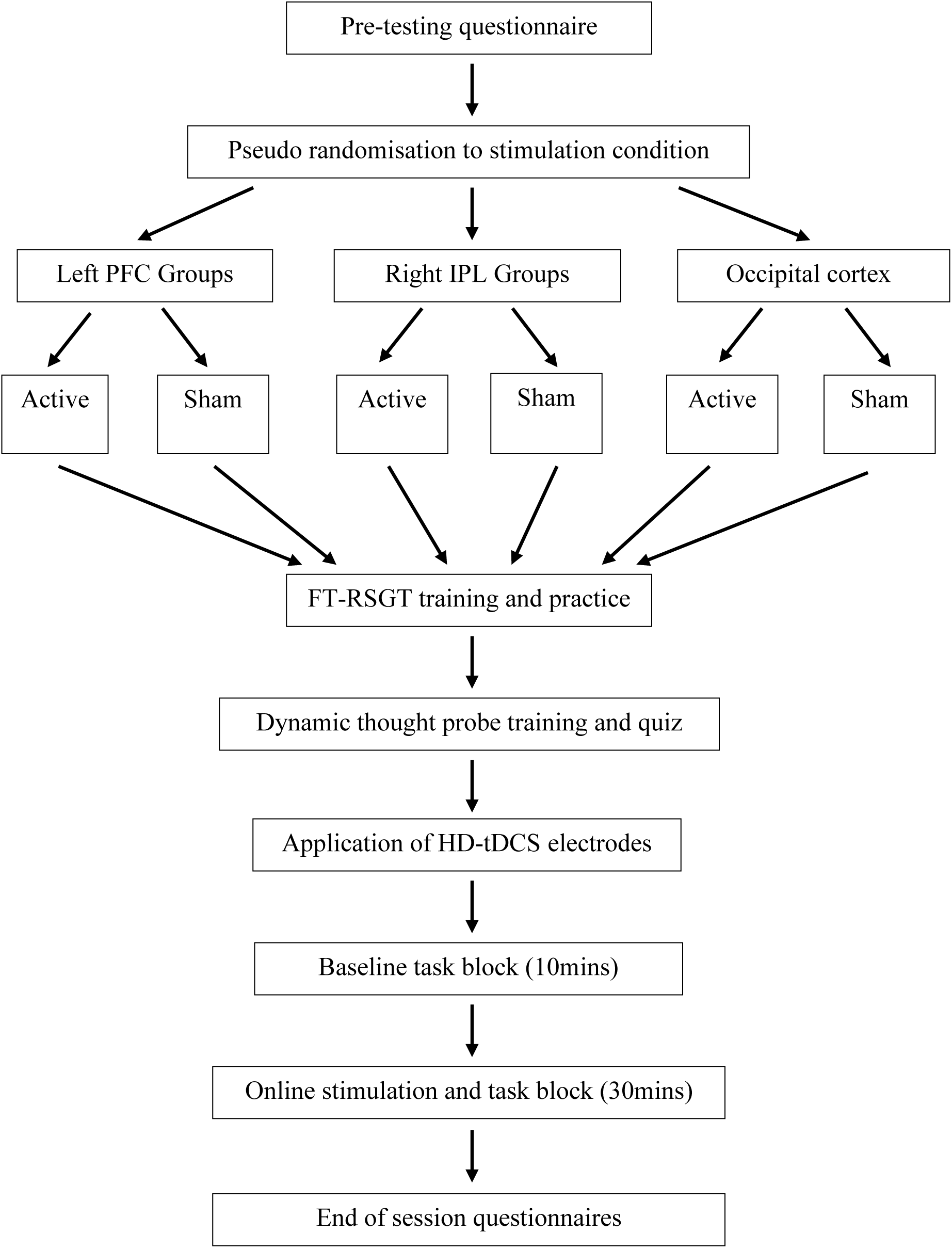
Conceptual diagram of the experimental procedure beginning with the pre-screening and pseudo random allocation to one of the 6 stimulation groups and followed by training on both the dynamic thought probes and the task, before participants complete a 10-minute baseline block and the 30-minute online stimulation and task block. The session then concludes with an end of session questionnaire and debriefing on the experiment.

### Participants

#### Recruitment and inclusion criteria

Participants aged 18-40 were recruited through the first-year student volunteer participant pool and also through the paid participant pool, The University of Queensland staff newsletter, and flyers around the St Lucia Campus. All participants were compensated with either course credit or $20 per hour for participation. To be included participants had to have normal or corrected to normal vision and hearing and they had to meet the following criteria: (1) English as their first language; (2) No current use of psychiatric medication(s); (3) No current or previous psychiatric/neurological condition(s); (4) No current use of psychotropic drugs; (5) Not currently taking part in other tDCS studies and they must meet the tDCS safety screening questionnaire criteria (see Appendix A; e.g., no metal in the head, or implanted neurostimulator). All participants provided written, informed consent and were right-handed. Participants were also excluded if they failed to understand the thought probes after training and additional clarification from the experimenter. Following recruitment and screening, participants were pseudo-randomly allocated to demographically balanced groups, using a script which factored in age, sex, time spent playing video games and musical instruments and time of day for testing. We also included a question assessing participants hours of sleep from the night before and this variable was included in the script to ensure sleep was equivalent across the groups. To be included in the analysis, participants were also required to comply with all the instructions throughout the experiment and the stimulation had to remain functional across the entire session. The University of Queensland Human Research Ethics Committee approved this study.

#### Sample size – Bayesian sampling plan

A Bayesian sampling approach was used, and thus, the sample size was predetermined for this study. A total of 251 participants were recruited, which resulted in a final sample of 228 participants after post-data exclusions. It was stated that participants would continue to be recruited until a Bayes Factor (BF)_10_ > 6 or BF_01_ > 6 was established for the critical hypothesis test (see below). However, recruitment was stopped when the maximum sample size of 240 (40 participants per group) was reached, as we believed inconclusive results at this sample size would still contribute meaningfully to the field. The stopping rule was first checked once 10 participants in each group were tested and for every 5 participants thereafter. The critical hypothesis test which was selected was the effect of stimulation on task unrelated thought in the PFC, which was the thought probe that broadly asks participants whether they were focused on or off task. Specifically, we predicted that there would be a difference in the frequency of task unrelated thoughts between participants who received active anodal HD-tDCS to the left PFC and those who received sham stimulation. We selected this test as Boayue et al. (2021) previously found that HD-tDCS to the DLFPC significantly decreased the frequency of task unrelated thought. However, as we did not find evidence for this effect in the t-test analysis, the maximum sample size was reached.

### Behavioural assessments

#### Finger-Tapping Random-Sequence Generation Task (FT-RSGT)

In the FT-RSGT task, adopted from Boayue et al. (2021), participants were required to respond in a random sequence to an ongoing metronome tone using two response-buttons (see Figure 4). The button presses were mapped to two separate keys which participants used their index fingers to press (‘z’ for left-hand and ‘m’ for right-hand). Participants were instructed to time their responses as accurately as possible to a metronome tone which was presented at 440Hz for 75ms duration, with an inter-stimulus interval of 750ms. They were also instructed to maintain their focus on a white (RGB 255 255 255) fixation cross in the centre of the screen with a grey (RGB 128 128 128) background for the duration of the experiment.

**Figure 4.**
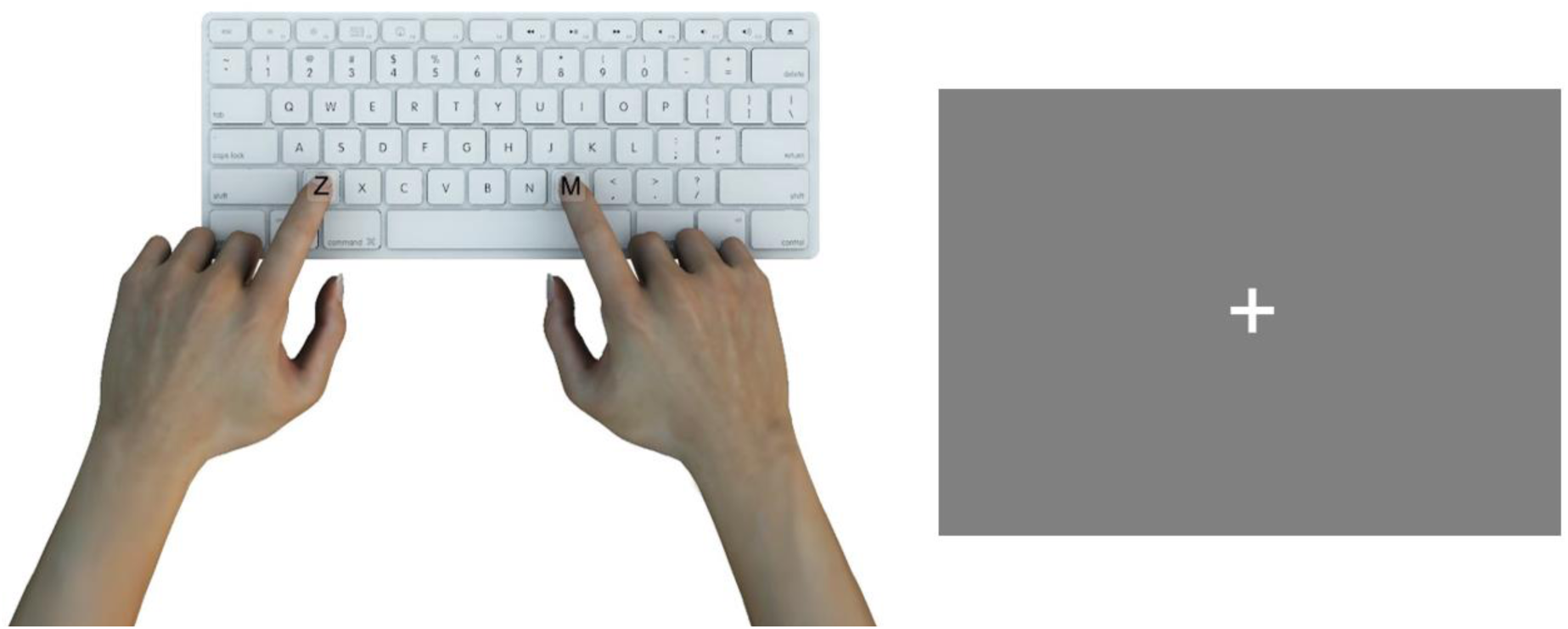
An illustration of the Finger-Tapping Random-Sequence Generation Task, showing the two keys’ participants used to generate the random sequences (on the left) and the fixation cross displayed during the task (on the right).

Performance on this task was determined by the randomness of the sequence, which was measured through approximate entropy (see Analysis Overview for detailed explanation). This measure was used as there is an expected relationship between executive functioning and randomness, such that diverting more executive resources towards the task will result in greater randomness which can be used to imply participants are more focused on the task (Boayue et al., 2021). Behavioural variability was also used to measure task performance and was calculated by the deviation in response times from the metronome tone. For the 20 trials before each set of mind wandering probes, the difference between participants key response and the tone was calculated and the standard deviation of the difference scores for each set of 20 trials was included in the analysis. This measure is designed to directly assess participants focus on the task and is also associated with executive functioning (Boayue et al., 2021).

The task was presented on a 24-inch LED monitor, with a refresh rate of 100 Hz. Participants sat approximately 70cm away from the monitor and used a standard Macintosh keyboard and mouse to respond. The auditory tone was presented through the computer speakers. Initially, participants completed 20 practice trials to familiarise themselves with the task alone. Following training on the thought probes, an additional 20 trials were completed which included the task and the four thought probes. Participants then completed a baseline block of 720 trials which ran for 10 minutes, followed by the stimulation block which consisted of 2160 trials over approximately 30 minutes, with a 30 second break after 15 minutes. The stimulation block was completed while participants received online active or sham stimulation to one of the three targeted brain regions.

#### Mind wandering probes

Throughout the experiment, participants were also presented with four thought probes which were designed to assess the contents of their thoughts. The probes were presented pseudo randomly, every 45 to 75 seconds during the baseline and stimulation blocks. Thus, there were 10 probes in the baseline block and 30 probes in the stimulation block. The questions assessed participants thoughts in the 10 to 15 seconds before the probes were presented. The four questions were: (1) Before the probe, were you thinking about something other than the random sequence generation task; (2) Before the probe, was your mind wandering around freely; (3) Were you actively directing your thoughts; and (4) Was your mind stuck on something. These questions were presented sequentially, with participants rating their responses on a 7-point Likert scale ranging from “Not at all” (1) to “Very much” (7), and the middle point (4) was labelled “Moderately”. Participants responded using the number keys labelled 1 through to 7 and they had an unlimited amount of time to categorise their thoughts before moving onto the next probe question.

Before participants completed the task, they were given detailed explanations on the four types of thought probes, alongside example scenarios where these thoughts may occur. The four dynamic thought probes, alongside explanations and questions given to participants replicated those used in research by Kam et al. (2021). Participants were then tested on their knowledge by categorising example scenarios, and the experimenter explained and corrected any incorrect answers. See Appendix B for the full description of the probes and the examples which were shown to participants and Appendix C for the four Likert scales which were shown during the experiment. All instructions and questions were presented in the centre of a grey background (RGB 128 128 128) in white Arial font (visual angle = 1.1°).

#### Self-report questionnaires

Prior to the task participants indicated how much time per week they spent playing musical instruments or video games (see Appendix D). This was included as the FT-RSGT requires fast responses to the metronome tone and thus it was important to account for any influence of musical or video game training on performance (Boayue et al., 2021). After the FT-RSGT task was completed, participants completed the Mindful Attention and Awareness Scale (MAAS), which was designed to assess participants disposition to focus on the present (see Appendix E). The MAAS has 15 items such as: “I find myself doing things without paying attention” and “I find myself listening to someone with one ear, doing something else at the same time”. Participants rated how frequently they currently had each experience on a 6-point scale ranging from “Almost Always” (1) to “Almost Never” (6). The mean of the 15 Items was used to calculate participants scores, with higher scores indicating higher levels of dispositional mindfulness.

There were two additional measures which were designed to assess participants distractibility and rumination, to account for group differences across these domains. The Adult ADHD Self-Report Scale was designed to identify the frequency participants exhibit symptoms associated with attention-deficit/hyperactivity disorder (ADHD) and consisted of eighteen questions (see Appendix F). The first six questions have been found to be the most predictive symptoms of ADHD, such as “When you have a task that requires a lot of thought, how often do you avoid or delay getting started?”. The remaining twelve questions probe additional areas which the participant may exhibit symptoms, such as “How often are you distracted by activity or noise around you?”. The questions are rated on a 5-point scale ranging from “Never” (1) to “Very Often” (5) and the total score for each section was calculated and added together to develop a total score, with higher scores indicating a greater number of symptoms consistent with ADHD. The Rumination Responses Scale was also included and consisted of 22 items that assessed how often participants engage in these thoughts when they are feeling down, sad, or depressed (see Appendix G). Participants rated responses on a 4-point scale ranging from “Almost never” (1) to “Almost always” (4) and the items include thoughts such as: “What am I doing to deserve this?” and “Why can’t I handle things better?”. The total score for participants was calculated, with higher scores indicating greater ruminative symptoms.

There was also a detailed questionnaire assessing participants involvement with musical instruments and video games, including the types of games or instruments they played, alongside their engagement in other forms of entertainment. Finally, participants completed an end of session questionnaire including a motivation assessment, asking participants “How motivated were you to perform well in this task?” (Seli et al., 2015). This question was rated on a 7-point scale ranging from “Not at all motivated” (1) to “Extremely motivated” (7). This questionnaire also assessed participants perspective on their task performance and stimulation experience. The full end of session questionnaire can be found in Appendix H.

### Stimulation protocol

Each participant received one of six stimulation protocols, which were delivered online during the FT-RSGT task: (1) anodal HD-tDCS over the left PFC; (2) sham HD-tDCS over the left PFC; (3) anodal HD-tDCS over the right IPL; (4) sham HD-tDCS over the right IPL; (5) anodal HD-tDCS over the VC; and (6) sham HD-tDCS over the VC (see Figure 5). This was a between-groups design, such that there was only one session for each group of participants. The stimulation conditions for each brain region were double-blinded, such that the experimenter and participant were both unaware whether participants are receiving active or sham stimulation. It was not feasible to blind participants to the location of the targeted brain region, as the electrodes were visible to the participants, however the participants were not informed of the study’s hypotheses. At the end of each participant’s session, they were asked which experimental condition they thought they were in – active or sham – to assess the effectiveness of the blinding. To ensure that the double-blinding was effective, the same question was also included for the experimenter to respond at the end of each testing session. In addition, a confidence rating was included for the experimenter and participants asking, “How confident are you in your judgement of your experimental condition?” (see Appendix H). This question consisted of a 7-point likert scale, ranging from “Not at all confident” (1) to “Extremely confident” (7), with the middle value as “Moderately confident” (4).

**Figure 5.**
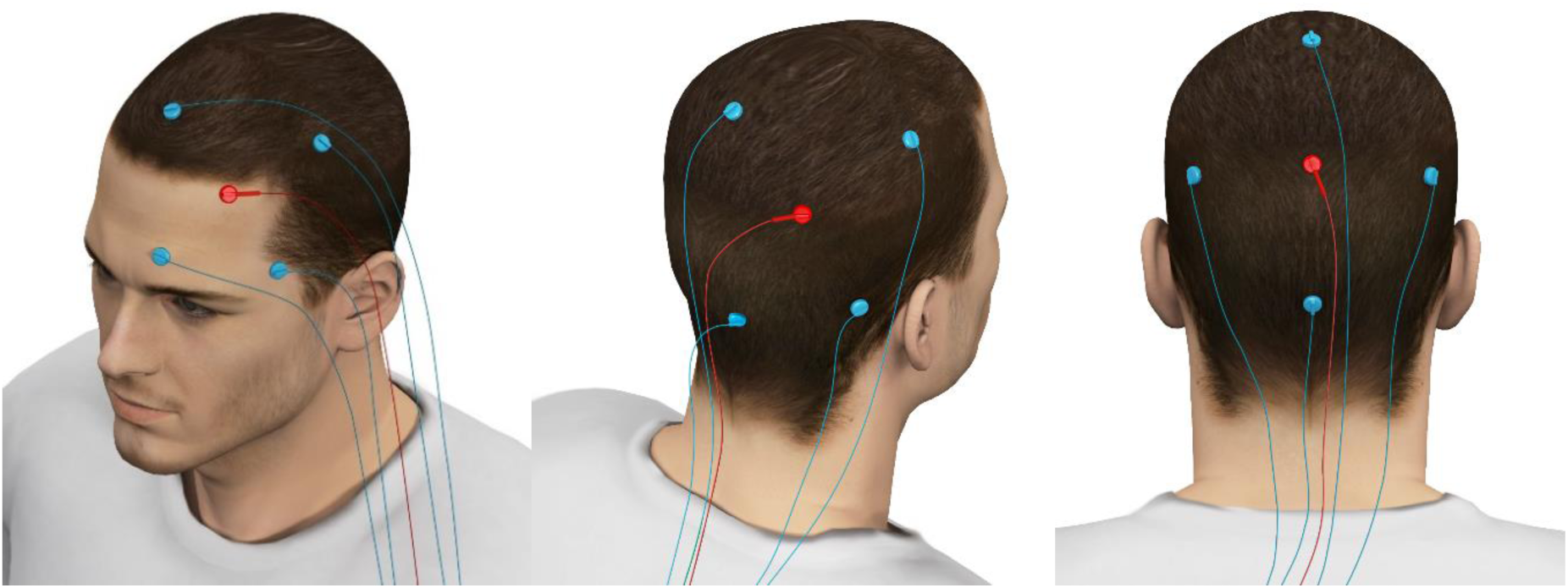
An illustration of the three HD-tDCS montages, showing the set up for the left PFC (far left), the right IPL (middle) and the occipital cortex (far right) with the anode (in red; positioned at F3, P4 and Oz, respectively) and four reference cathodes for each montage.

#### HD-tDCS montage

The HD-tDCS was administered using a Nurostym stimulator, with a 4 x 1 electrode ring arrangement. The stimulator was manufactured in the United Kingdom, by the Neuro Device Group. The electrodes were 12mm Ag/AgCl electrodes which were attached to the scalp using a conductive gel. All electrode placements were determined using the International 10-20 EEG system. For the left DLPCF stimulation, the anode was placed over F3, and the four reference cathodes were placed on F7, Fp1, C3, and Fz. This configuration was chosen to replicate the HD-tDCS montage used by Boayue et al. (2021). The right IPL stimulation groups had the anode placed on P4 and the surrounding cathodes placed at T6, O2, Pz and C4. There was evidence anodal bipolar tDCS to this region also had an effect on mind wandering propensity (Kajimura et al., 2019; Kajimura & Nomura, 2015), thus it was included to explore the effects using the anodal HD-tDCS. Finally, the VC groups had the anode placed at Oz with the upper cathode at Pz and three additional cathodes were measured to each point, one below Oz and one straight out to the left and one to the right of Oz, at an equal radius as the distance from Oz to Pz. The VC was included as an active control region (Filmer et al., 2020). It is important to note that there has been research which utilises more focal HD-tDCS designs, with the electrode placements in more proximal locations such as the 3×1 montage used by Chou et al. (2020). However, the current study was designed to replicate the findings of Boayue et al. (2021) in the PFC using the same task and to expand on this research into the IPL and VC, with the inclusion of the additional dynamic thought probes, thus the more distal 4×1 montage used by Boayue et al. (2021) was the most applicable for the three regions being investigated in this research.

All stimulation groups used 2.0mA current density with 0.5mA to each of the four reference electrodes (Boayue et al., 2021). The anodal stimulation lasted 30 minutes, including a 30 second ramp up and ramp down period. Participants in the sham conditions experienced 75 seconds of stimulation, with the same ramp up and ramp down times as the active stimulation groups. All participants initially completed the baseline block of the FT-RSGT, before then completing the 30-minute task block with stimulation (active or sham). If participants experienced any discomfort, the stimulation was discontinued, and alternatively if there are any technical issues that affected the intensity or duration of the stimulation, or if the electrode impedance increased above the programmed impedance cut-off, then the session did not proceed.

### Analysis overview

All analyses were completed in Rstudio using the *brms* (Bayesian Regression Models using Stan; Bürkner, 2017) and *BayesFactor* packages (Mulder et al., 2021), or using JASP. All analyses employed a default Jeffreys-Zellner-Siow prior of r = 0.707, centred around 0 (Rouder et al., 2009). We chose to use a default prior as there were no previous studies which investigated the causal neural substrates of dynamic thoughts. The results assessed which model the observed results were more likely under, as indicated by BF_10_ (alternative) and BF_01_ (null). A BF_10_ of 1-3 or BF_01_ of 1-3 was considered anecdotal evidence for the alterative or null hypothesis, respectively. BF_10_ of 3-10 or BF_01_ of 3-10 was interpreted as moderate evidence in for the respective hypothesis, and BF_10_ > 10 or BF_01_ > 10 was considered strong evidence for the alterative or null hypothesis, respectively (Lee & Wagenmakers, 2013). We also reran the analyses for all four dynamic thought types using frequentist methods for completeness and to allow researchers who are more familiar with p-values to fully comprehend the results. All the raw data, study materials and analysis scripts from our study can be located at https://doi.org/10.48610/74fcc20.

#### Approximate entropy

An approximate entropy (ApEn) value was employed to calculate the randomness of a participant’s generated sequence, as it is mathematically impossible to calculate the entropy of a finite sequence. ApEn is designed to evaluate the randomness in a sequence of *m* numbers, by calculating the predictability of the next item in a sequence, given the previous sequence of *m* numbers (Pincus, 1991; Pincus & Singer, 1996; Yentes et al., 2013). A value of *m* = 2 was used, which replicates that employed by Boayue et al. (2021), as this value was calculated to be the most sensitive for detecting differences in the randomness of sequences for the FT-RSGT. A smaller ApEn value indicated a data set contains many repetitive patterns, whereas a larger ApEn score indicated less predictable response patterns in the data.

### Data exclusions

After participants completed the study, the data was pre-processed and individuals who scored 3 standard deviations above or below the mean for approximate entropy and behavioural variability across all participants in the baseline block were excluded from the study. This resulted in 8 participants being removed. Furthermore, we assessed participants responses to the end of session questionnaire and if the answers suggested that the participant did not generate random number sequences for the task correctly, we excluded their data before the analyses were conducted, thus they were replaced in the final sample. An example which would suggest the task has not been completed correctly would be if the participant cites a specific pattern that they used to approach the task (e.g. they repetitively used z,z,m,m,m,z,z,m,m,m to generate the sequences). An additional exclusion criterion which was not included in the stage one manuscript was that if any participants scored more than 3 standard deviations above or below the mean for approximate entropy and behavioural variability or the four thought probe responses, across each region in the stimulation block, they were also removed. This resulted in an additional 3 people being removed, creating a final sample of 228 participants. Finally, to avoid extreme behavioural variability and randomness scores skewing the time-on-trial analyses, trials which were above or below 3 standard deviations for each brain region were removed post-hoc. This resulted in 38 trials (1.25% of trials) being removed for the PFC groups, 53 trials (1.74%) being removed for the IPL groups and 42 trials (1.38%) for the VC groups being removed, with a final trial sample of 3002, 2987 and 2998 for each region, respectively.

## Results

### Using modelling to investigate the effect of HD-tDCS on the dynamic thought types

To investigate the effect of HD-tDCS on each dynamic thought type, we employed hierarchical order probit modelling (Alexandersen et al., 2022; Boayue et al., 2021; Filmer et al., 2019). Each probe and region analysis consisted of running 23 models including the predictor variables of behavioural variability, approximate entropy, trial, block (baseline vs. stimulation) and stimulation group (active vs. sham), alongside their interactions (see Figure S1 for full model comparison list). This approach allowed the thought probe responses to be treated as an ordinal variable, which has been argued to be a more accurate approach to investigate the response variability within an individual’s performance and to account for time-on-task effects (Boayue et al., 2021; Filmer et al., 2019). The 23 models increased in complexity and the model weights were interpreted using two different methods. They were first calculated using pareto smoothed importance sampling leave-one-out cross-validation scores (PSIS-LOO; Vehtari et al., 2017, 2022), whereby the LOO information criterion values (LOOIC; Vehtari et al., 2017; Wagenmakers & Farrell, 2004) were compared using a stacking procedure (Vehtari & Gabry, 2018; Yao et al., 2018). Second, the model weights were assessed using pseudo-Bayesian model-averaging (pseudo-BMA;Vehtari & Gabry, 2018; Yao et al., 2018).

Where there were discrepancies between the LOOIC and pseudo-BMA winning models, the preferred LOOIC model was selected, as it has more reliable predictive accuracy by selecting the optimal predictive distribution, rather than being biased towards the model closest to the Kullback-Leibler divergence (as the pseudo-BMA method is) when the true model is not included in the model list (Vehtari & Gabry, 2018; Yao et al., 2018). When these discrepancies occurred, we ran additional post-hoc exploratory analyses to confirm our findings. Given the main analyses included the baseline data which had relatively low numbers of trials (720), the model comparisons were re-run on the stimulation block alone, to ensure any potential confounds due to the differences in probe and trial numbers between the baseline and stimulation blocks were removed. The additional analyses on the stimulation block data can be seen in the exploratory analyses section.

#### Stimulation did not modulate propensity for task unrelated thoughts

Our critical hypothesis test investigated the effect of PFC HD-tDCS on task unrelated thought, however no such stimulation effect was observed. Specifically, the key predictor indicating an effect of stimulation on the thought probe responses was the stimulation x block interaction. The model favoured by the LOOIC method included only main effects for randomness, behavioural variability, block, trial, and a behavioural variability x randomness interaction (*p_PSIS-LOO_* = 0.24; see Figure S2 for winning model). Similarly, for IPL stimulation the preferred LOOIC model included only a main effect of behavioural variability, randomness, and block, alongside a behavioural variability x randomness interaction (*p_PSIS-LOO_* = 0.24; see Table S1 for the top model selection weights). In both regions the LOOIC and Pseudo-BMA weights did not agree on the preferred model. Thus, these findings suggest HD-tDCS did not influence task unrelated thoughts for these two regions.

No influence of stimulation was also observed for the VC. The preferred model did not agree between the selection procedures, but the winning LOOIC model (*p_PSIS-LOO_* = 0.25) only included a meaningful effect of behavioural variability, block, and a positive behavioural variability x randomness interaction. There was no evidence for a main effect of stimulation or a block x stimulation interaction. There was also a meaningful randomness x stimulation interaction (*b* = 0.13, 95% credible interval (CI) [0.03, 0.22]), however as the stimulation variable was averaged across the task blocks, this was not a direct effect of stimulation on randomness. Overall, the missing block x stimulation interaction shows that HD-tDCS also did not influence task unrelated thought in the VC.

#### Freely moving thoughts were influenced by PFC stimulation

The modelling revealed PFC stimulation reduced freely moving thought. Specifically, the critical block x stimulation interaction showed that in the stimulation block, relative to the baseline block, freely moving thought was reduced in the active group compared to the sham condition (*b* = -0.31, 95% CI [-0.69, 0.07]). The winning Pseudo-BMA and LOOIC models did not align, however the preferred LOOIC model (*p_PSIS-LOO_* = 0.49) also included main effects of behavioural variability, randomness, block, trial, and stimulation, alongside a behavioural variability x randomness interaction (see Figure 6). There was no main effect of stimulation alone (*b* = -0.03, 95% CI [-0.50, 0.43]), as would be expected as this included pre-stimulation (baseline) data.

**Figure 6.**
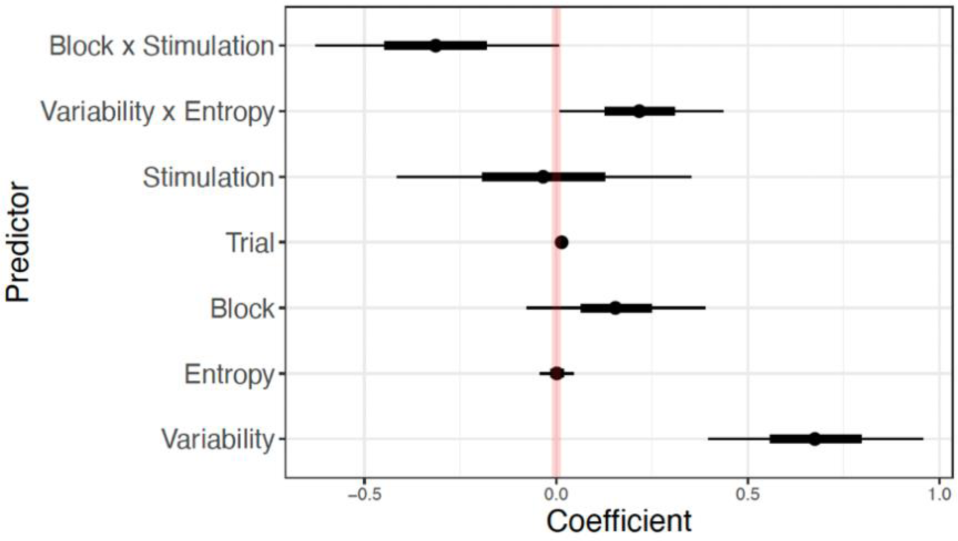
The PFC winning model for freely moving thought.

For participants receiving stimulation to the VC, the winning LOOIC model (*p_PSIS-LOO_* = 0.32) was consistent with the PFC findings and included main effects of behavioural variability, randomness, block, trial, and stimulation, alongside a behavioural variability x randomness and block x stimulation interaction (see Figure 7). However, there was no evidence for a block x stimulation interaction for this model (*b* = -0.08, 95% CI [-0.39, 0.23]). In contrast, there was evidence for a main effect of stimulation (*b* = 0.51, 95% CI [0.07, 0.96]), averaged across the two task blocks. This suggests there is a difference between the stimulation groups, however this is not a function of task block, and thus there is no effect of stimulation in the VC.

**Figure 7.**
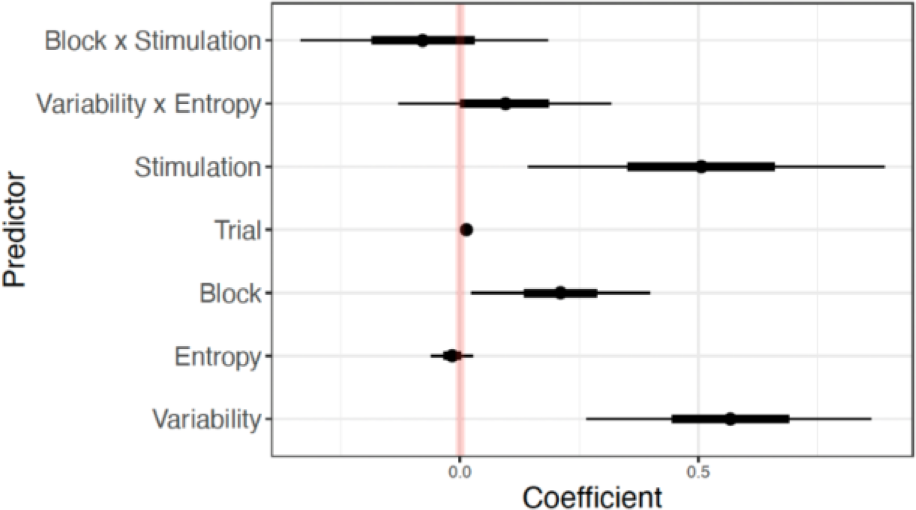
The VC winning model for freely moving thought.

For the IPL, the preferred model did not align between the model selection procedures, and the winning LOOIC model (*p_PSIS-LOO_* = 0.34) found no evidence for effects of stimulation on freely moving thought, only a main effect of behavioural variability, randomness, block, trial, and stimulation, alongside a behavioural variability x randomness interaction.

#### Deliberately constrained thoughts were reduced by IPL stimulation

Anodal stimulation of IPL reduced deliberately constrained thought as was indicated by a negative block x stimulation interaction (*b* = -0.52, 95% CI [-1.02, - 0.01]). Here, the Pseudo-BMA and LOOIC weights agreed upon winning model (*p_PSIS-LOO_* = 0.38), which also included weak evidence for an effect of trial and a block x trial x stimulation interaction (for both active and sham as the reference condition), and no evidence for an effect of trial or a trial x stimulation interaction (see Figure 8). There was also no evidence for a main effect of stimulation, averaged across the two blocks (*b* = 0.43, 95% CI [-0.13, 1.02]) which supports the changes in deliberately constrained thought reported in the active condition, being specific to the differences found between the baseline and stimulation blocks.

**Figure 8.**
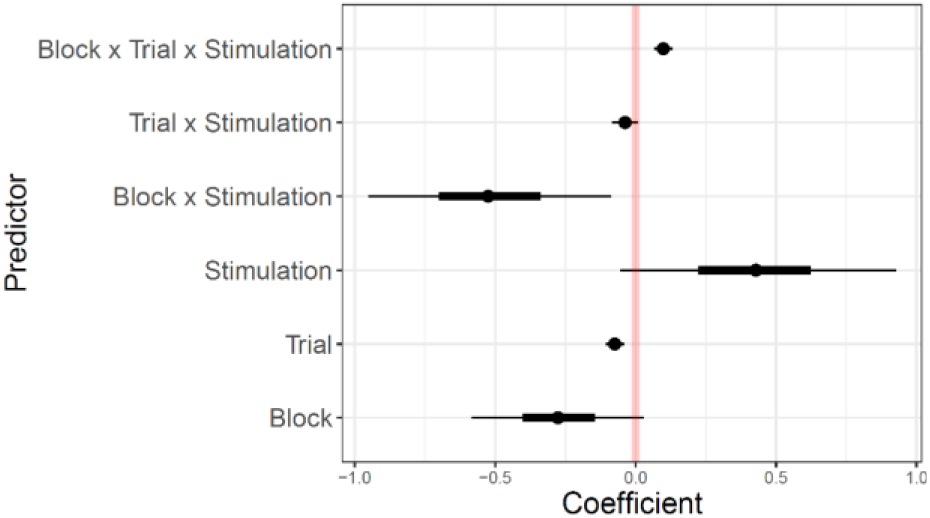
The IPL winning model for deliberately constrained thought.

Stimulating the PFC or the VC did not modulate deliberately constrained thoughts. For the PFC, the winning LOOIC model only included main effects of behavioural variability, randomness, and block, alongside a behavioural variability x randomness interaction (*p_PSIS-LOO_* = 0.20). However, this model was not selected by the Pseudo-BMA procedure. While the model selections agreed for the VC (*p_PSIS-LOO_* = 0.25), the winning model did not include any evidence for a main effect or interaction including stimulation.

#### Stimulation did not modulate automatically constrained thoughts

For the automatically constrained thought probe, there was no evidence of stimulation effects in any region, with all three winning models not including the stimulation predictor in any main effects or interactions. In addition, the winning models did not agree between the Pseudo-BMA and LOOIC procedures for any region. In the PFC, the winning LOOIC model (*p_PSIS-LOO_* = 0.18) only included a main effect of behavioural variability. The IPL winning LOOIC model (*p_PSIS-LOO_* = 0.26) had no evidence of meaningful effects, with all the credible intervals crossing 0. Finally, in the VC, the winning LOOIC model (*p_PSIS-LOO_* = 0.40) only found a meaningful main effect of behavioural variability and a behavioural variability x randomness interaction.

### The relationship between task performance and dynamic thought probe responses

In addition to the probit modelling, we also conducted a simplified analysis investigating the relationship between task performance and the dynamic thought responses. We calculated the approximate entropy and behavioural variability mean values for trials that occurred 15 seconds before the thought probes (*n* = 20 trials) throughout the stimulation block. The mean values for approximate entropy and behavioural variability were then compared for on-task and off-task responses, averaged across conditions. Scores of 1-3 were considered on-task and scores of 4-7 were considered off-task, with 4 categorised as off-task for this experiment because the labelling of the Likert scale and phrasing of the questions indicated 4 was “moderately” high.

As anticipated, and across all dynamic thought types, randomness was lower when off-task and higher when on-task (see Figure 9). Also, in line with our predictions, ratings of being off-task for freely moving thoughts led to higher, and on-task lower, behavioural variability. However, for the other three thought types, behavioural variability showed the opposite pattern: off-task leading to reduced variability compared to on-task. This may reflect that due to the nature of our paradigm and tasks requirements participants prioritised generating random sequences over accurately timing their responses to the tone.

**Figure 9.**
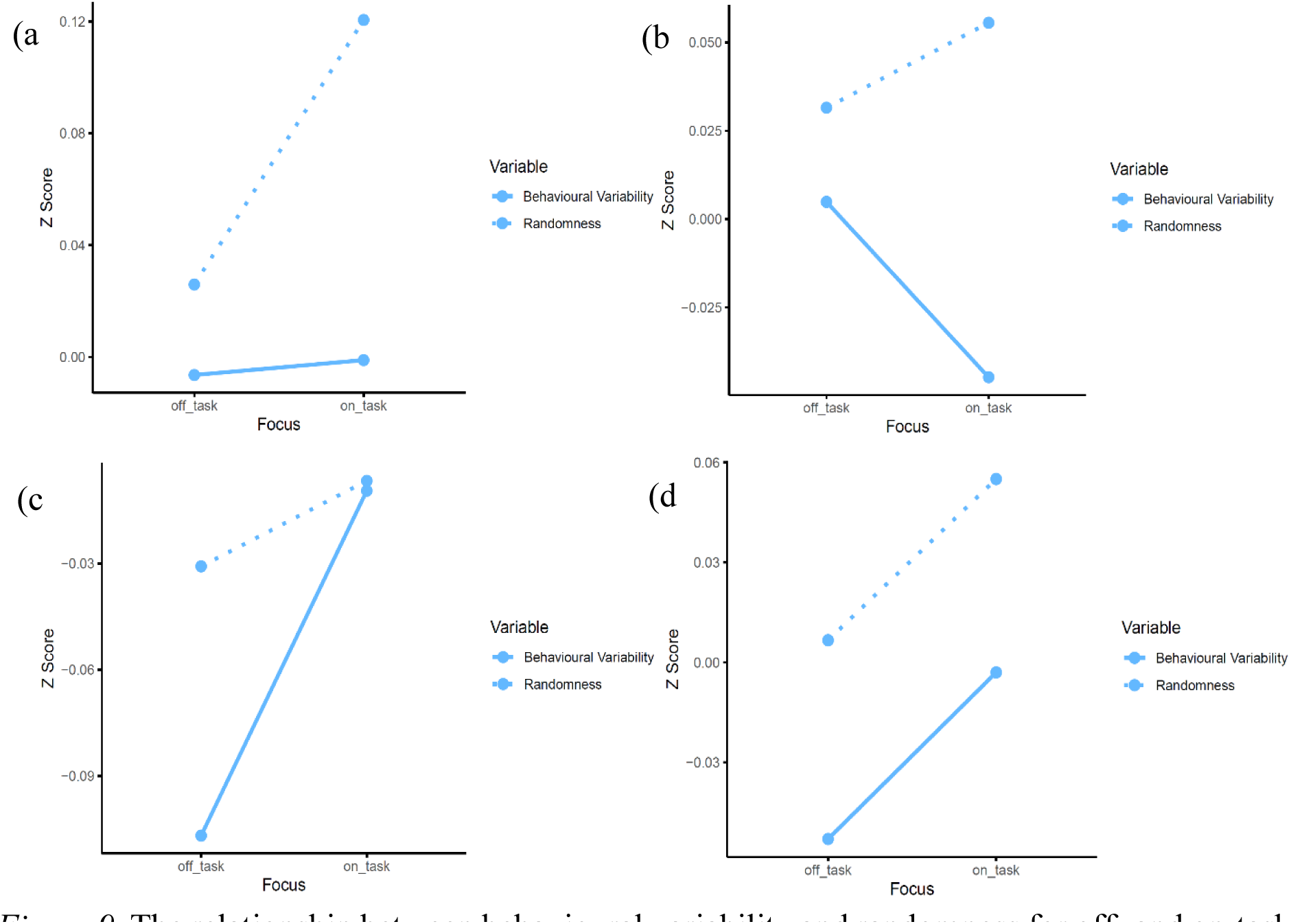
The relationship between behavioural variability and randomness for off- and on-task thoughts across (a) task unrelated thought, (b) freely moving thought, (c) deliberately constrained thought, (d) automatically constrained thought.

### Stimulation modulated dynamic thought

We also investigated the direct relationship between stimulation and participants mean thought probe responses using a series of Bayesian independent samples t-tests. For each thought probe and brain region, we compared the anodal HD-tDCS group to the sham group. We also ran three 2 (Stimulation: active, sham) x 2 (Region: PFC, IPL, or VC) Bayesian between-subjects ANOVAs for each probe to assess the overall effect of stimulation across the three brain regions on the frequency of the dynamic thought responses.

#### Task unrelated thought

Consistent with our modelling analyses, there was no evidence for meaningful differences between active and sham stimulation in any brain region for task unrelated thought. These findings indicate that HD-tDCS did not affect participants task unrelated thoughts in any brain region. There was no evidence for any effects found in the three Bayesian ANOVAs (BF_excl_ > 1.84 for all). The Bayesian independent sample t-tests also found no evidence across the three regions for a difference between the active and sham groups (BF_01_ > 1.35 for all).

#### Freely moving thought

There was a meaningful difference between the active and sham conditions in the PFC and VC for freely moving thought. These effects expand on the modelling results and suggest that anodal HD-tDCS may have differential effects on the frequency of freely moving thought across the brain. Specifically, anodal stimulation reduced freely moving thought in the PFC and increased freely moving thought in the VC (see Figure 10). This was revealed in the 2 (Stimulation: active, sham) x 2 (Region: PFC, VC) ANOVA which showed strong evidence for the inclusion of the interaction term [BF_incl_ = 11.12, *F(*1, 148) = 8.145, p = .005, *η_p2_* = 0.052]. The t-tests also found anecdotal evidence for a difference between the active and sham conditions in the PFC [BF_10_ = 1.93 (M_Active_ = 3.51, M_Sham_ = 4.08), *t*(74) = -1.84, *p* = .035, *d* = -0.422)] and in the VC [BF_10_ = 1.89 (M_Active_ = 4.11, M_Sham_ = 3.46, *t*(74) = 2.21, *p* = .030, *d* = 0.507)]. There was no evidence for a difference between the active and sham groups in the IPL [BF_10_ = 0.32 (M_Active_ = 3.99, M_Sham_ = 4.10, *t*(74) = -0.38, *p* = .705, *d* = -0.087)], and no evidence for an effect in this region in the Bayesian between subject ANOVAs (BF_excl_ > 0.90).

**Figure 10.**
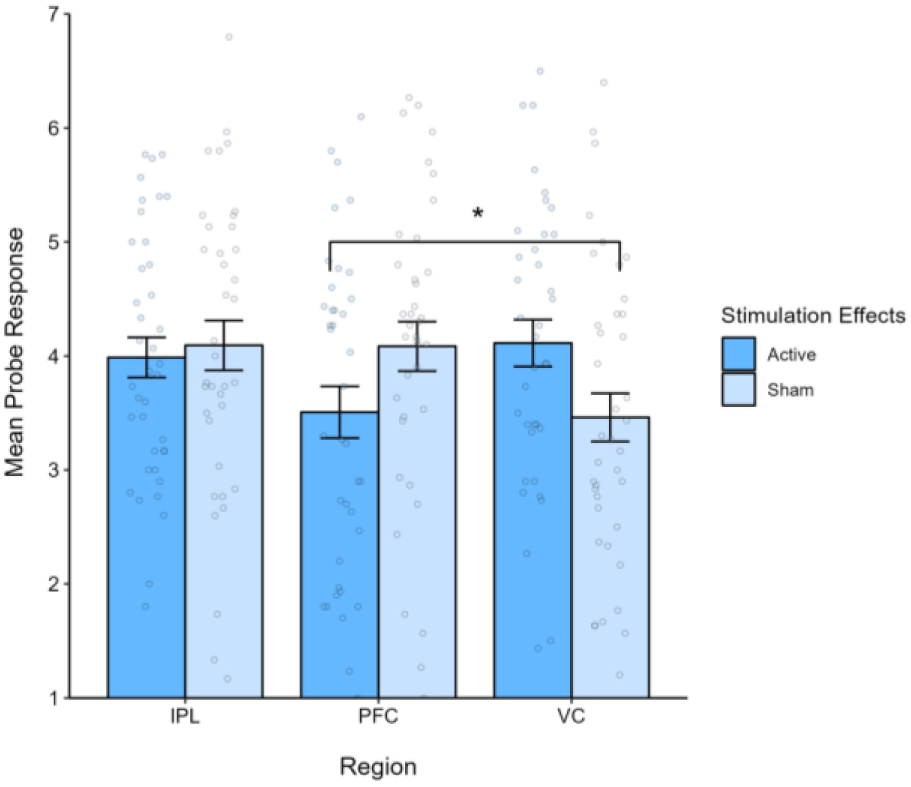
The effect of anodal HD-tDCS on freely moving thought in the three target brain regions.

#### Deliberately constrained thought

There was no evidence for an effect of HD-tDCS on deliberately constrained thought, with no differences found between the active and sham conditions for the three brain regions in the t-tests (BF_01_ > 3.08 for all). This aligns with the modelling analyses for the PFC and VC; however, it does not reflect the meaningful stimulation x block interaction effect found in the IPL. This may be due to the improved sensitivity of the hierarchical order probit modelling to detect time-on-task effects across the task, rather than taking an average for each participant’s probe responses.

#### Automatically constrained thought

There was no evidence for differences between the active and sham conditions found in any of the three brain regions for automatically constrained thought (BF_01_ > 1.56 for all). These findings align with the modelling analyses and suggest that anodal HD-tDCS did not affect the reporting of automatically constrained thoughts.

### Testing for baseline differences

Collectively there was little evidence that any differences at baseline had an impact on the above findings. To ensure there were no meaningful differences between the groups in their baseline performance, we reran the primary 12 Bayesian independent samples t-tests on the baseline data alone (see Table 1). A BF_01_ ≥ 3 indicated there was moderate evidence for no meaningful differences between the groups, and thus the null hypothesis was accepted. All baseline comparisons were in favour of the null with one exception. For VC stimulation, there was anecdotal evidence for an effect (BF_10_ = 1.89).

**Table 1.** Descriptive statistics, sample size, and independent samples t-tests for all baseline tests

| Probe | Region | Active | | Sham | | $BF_{01}$ | $t$ | $p$ | $d$ |
| --- | --- | --- | --- | --- | --- | --- | --- | --- | --- |
| | | $M$ (SD) | $n$ | $M$ (SD) | $n$ | | | | |
| Task unrelated thought | PFC | 3.57 (1.46) | 38 | 3.49 (1.26) | 38 | 4.09 | 0.26 | .794 | 0.060 |
|  | IPL | 3.72 (1.21) | 39 | 3.71 (1.44) | 37 | 4.21 | 0.04 | .968 | 0.009 |
|  | VC | 3.67 (1.47) | 37 | 3.75 (1.28) | 39 | 4.08 | -0.27 | .792 | -0.061 |
| Freely moving thought | PFC | 3.46 (1.42) | 38 | 3.61 (1.33) | 38 | 3.79 | -0.49 | .624 | -0.113 |
|  | IPL | 3.69 (1.16) | 39 | 3.86 (1.62) | 37 | 3.76 | -0.51 | .611 | -0.117 |
|  | VC | 3.73 (1.25) | 37 | 3.07 (1.37) | 39 | 0.53 | 2.21 | .030 | 0.507 |
| Deliberately constrained thought | PFC | 3.96 (1.52) | 38 | 4.08 (1.27) | 38 | 3.96 | -0.38 | .707 | -0.086 |
|  | IPL | 4.20 (1.52) | 39 | 3.89 (1.54) | 37 | 2.82 | 0.96 | .339 | 0.221 |
|  | VC | 3.94 (1.40) | 37 | 4.08 (1.62) | 39 | 3.95 | -0.38 | .702 | -0.088 |
| Automatically constrained thought | PFC | 1.89 (1.89) | 38 | 2.00 (1.04) | 38 | 3.76 | -0.51 | .612 | -0.117 |
|  | IPL | 2.17 (1.09) | 39 | 2.00 (0.87) | 37 | 3.34 | 0.73 | .466 | 0.168 |
|  | VC | 2.30 (1.09) | 37 | 2.31 (1.50) | 39 | 4.21 | -0.02 | .987 | -0.004 |

Further to this, to investigate any group differences in responses to the three self-report measures we used three one-way independent groups ANOVAs. These included the MAAS, the Adult ADHD Self-Report Scale, the Rumination Responses Scale as the dependent variable, respectively, and the six stimulation groups as the independent variable (anodal and sham over the PFC, IPL, and VC, respectively). These analyses revealed strong evidence for excluding the group variable in all three questionnaires (BF_excl_ > 34.79 for all), suggesting there were no differences between the stimulation groups.

### Assessing blinding

It was also important to ensure that the HD-tDCS blinding was effective across the groups and did not contribute to the pattern of results observed. Thus, we assessed the proportion of correct guesses for people in the active compared to sham conditions. Overall, the sham blinding was effective in blinding people to their stimulation condition. Specifically, this analysis revealed that 115 out of 228 participants correctly guessed their group (50%). Generally, there was a bias for participants to select active stimulation regardless of group. In the active group, 99 participants out of 114 correctly guessed their group (87%) and 15 guessed incorrectly (13%). However, in the sham group only 16 participants, out of 114, correctly guessed their group (14%) and 98 participants guessed incorrectly (86%).

To further understand the effect of subjective belief on the frequency of the dynamic thought types, we implemented two Bayesian ANOVAs for each thought probe, employing objective intervention and subjective intervention as between-subject factors and the outcome measure of the average ratings for each thought probe (Gordon et al., 2022). The first set of ANOVAs compared all stimulation groups for each region (see Table S2); however, the second set of ANOVAs excluded conditions and probes which showed limited evidence for differences between stimulation and sham according to the modelling analyses, thus only evaluating conditions that demonstrate meaningful results (see Table 2). For the freely moving thought results, in the PFC which found evidence for stimulation effects in the modelling analyses, the results showed the strongest predictor was the objective stimulation. The IPL result for the deliberately constrained thought found no evidence for both objective and subjective condition.

**Table 2.** Blinding results for the objective and subjective stimulation effects, specifically for the meaningful thought types found in the probit modelling.

| Probe | Region | | Active | | Sham | | BF<sub>10</sub> | BF<sub>incl</sub> | F | p | $\eta_p^2$ |
| --- | --- | --- | --- | --- | --- | --- | --- | --- | --- | --- | --- |
|  |  |  | M (SD) | n | M (SD) | n |  |  |  |  |  |
| Freely moving thought | PFC | Objective | 3.51 (1.40) | 38 | 4.08 (1.33) | 38 | 1.01 | 1.00 | 0.75 | .390 | 0.010 |
|  |  | Subjective | 3.73 (1.35) | 65 | 4.17 (1.63) | 11 | 0.46 | 0.45 | 0.90 | .347 | 0.012 |
|  |  | Objective * subjective |  |  |  |  |  | 0.48 | 0.31 | .582 | 0.004 |
| Deliberately constrained thought | IPL | Objective | 3.72 (1.38) | 39 | 3.81 (1.50) | 37 | 0.25 | 0.24 | 0.97 | .327 | 0.013 |
|  |  | Subjective | 3.82 (1.38) | 67 | 3.35 (1.78) | 9 | 0.47 | 0.47 | 0.64 | .425 | 0.009 |
|  |  | Objective * subjective |  |  |  |  |  | 0.59 | 1.20 | .277 | 0.016 |

We also reran the proportion analyses on the experimenter blinding responses. Overall, blinding was effective. The experimenter made 128 correct guesses out of 228 sessions (56%). In the active group, 48 out of 114 guesses were correct (42%) and in the sham group 80 out of 114 guesses were correct (70%).

### Exploratory Analyses

In addition to the hierarchical order probit modelling conducted on the full dataset, an exploratory analysis was conducted when there were discrepancies between the LOOIC and Pseudo-BMA winning models. In these cases, the same model comparisons were re-run on the stimulation block data alone, which resulted in 16 models being compared for relevant thought probes and regions. The same model weighting techniques were applied and to be consistent with the overall analyses conducted above, if there were conflicting findings, the winning LOOIC model was selected (Vehtari A & Gabry J, 2018; Yao et al., 2018). The full list of models and model comparisons can be seen in Figure S3.

#### Task unrelated thought stimulation block modelling

There was no effect of stimulation found for task unrelated thought in the stimulation block alone across the three brain regions. The preferred LOOIC model for the PFC (*p_PSIS-LOO_* = 0.48; see Table S3 for the top model selection weights) found no evidence for an effect of stimulation (*b* = -0.23, 95% CI [-0.76, 0.28]). Furthermore, in the VC and IPL, there was no stimulation effect included in the winning LOOIC models (*p_PSIS-LOO_* = 0.60 and 0.39, respectively). See Figure S4 for the stimulation block winning model predictors.

#### Freely moving thought stimulation block modelling

Stimulating the PFC was again found to reduce freely moving thoughts via a main effect of stimulation (*b* = -0.34, 95% CI [-0.81, 0.08]), supporting the main finding above. Interestingly, anodal stimulation to the VC had an opposing effect, whereby freely moving thoughts were increased in the active group relative to sham (*b* = 0.43, 95% CI [0.01, 0.83]). Despite not finding evidence for a block x stimulation interaction in the main findings, this main effect of stimulation suggests anodal HD-tDCS may increase freely moving thought in the VC, relative to sham. For both the PFC and VC, the preferred LOOIC model also found evidence for an effect of behavioural variability and trial (*p_PSIS-LOO_* = 0.35 and 0.46, respectively). Finally, the stimulation results for the IPL winning model (*p_PSIS-LOO_* = 0.34) were consistent with the main findings and showed no effect of stimulation (*b* = -0.29, 95% CI [-0.71, 0.11]).

#### Deliberately constrained thought stimulation block modelling

In the PFC, there was no evidence for an effect of stimulation on deliberately constrained thought in the stimulation block alone (*b* = 0.18, 95% CI [-0.24, 0.61]), which mirrors the overall analyses (*p_PSIS-LOO_* = 0.37). In addition, because the model selection procedures agreed on a preferred model for the IPL and VC analyses above, there was no exploratory modelling conducted in these regions.

#### Automatically constrained thought stimulation block modelling

Consistent with the overall modelling above, there were no effects of stimulation across the three brain regions on automatically constrained thoughts for the stimulation data. For the PFC and IPL, the winning LOOIC models did not include the stimulation predictor (*p_PSIS-LOO_* = 0.41 and 0.28, respectively). Finally, in the VC, the agreed on preferred model (*p_PSIS-LOO_* = 0.41) showed no evidence the main effect of stimulation (*b* = 0.33, 95% CI [-0.16, 0.76]).

## Discussion

Here, we investigated whether different internal thought types were associated with distinct causal neural substrates using HD-tDCS, applied to three brain regions in a sham-blinded, active control study. Our results found that anodal stimulation had differential effects across the brain regions examined, specifically reducing freely moving thought in the PFC and reducing deliberately constrained thought in the IPL. There was also preliminary evidence for an effect of stimulation increasing freely moving thought in the VC. However, there was no evidence for stimulation affecting task unrelated thought or automatically constrained thought in any brain region. These findings highlight the importance of understanding mind wandering as a heterogeneous construct which recruits a range of neural networks according to the current direction of the individuals’ thoughts (Kam et al., 2021; Martel et al., 2019).

The evidence against stimulation affecting task unrelated thought did not support our hypothesised effect of stimulation reducing these thoughts, however it aligns with the recent replication of Boayue et al. (2021) by Alexandersen et al. (2022), who found no evidence for the original effect of 2mA anodal HD-tDCS reducing mind wandering in the PFC. Given the inconsistencies in mind wandering literature to date, it is important to understand the methodological practices which may affect these results (Alexandersen et al., 2022; Boayue et al., 2020; Chaieb et al., 2019). In this study, the groups were demographically balanced with large sample sizes in each group, which reduces the possibility of individual differences causing differences between the groups. In addition to the large sample size, the sham blinding was effective for both the subjects and experimenter which suggests the current findings were not driven by a placebo effect. Thus, these results suggest a measure of task unrelated thought may not always be sensitive to detecting shifts to internal thought processes.

The reduction in freely moving thought found in the PFC, however, did support our hypothesised effect of anodal stimulation reducing the novel thought types. This finding converges with EEG research by Kam et al. (2021), who found frontal alpha power was associated with freely moving thoughts. Furthermore, this result highlights the potential for freely moving thought to act as a more sensitive measure for detecting mind wandering. To wit, freely moving thought has been found to be independent of task-relatedness (Kam et al., 2021; Mills et al., 2018), and it has also been linked to creative and spontaneous thought (Christoff et al., 2016), which may represent a more valid assessment of mind wandering. In the VC, there was also weak evidence for an increase in freely moving thought, given the meaningful stimulation effect in the stimulation block alone. While the weak evidence for a baseline difference in the VC group for freely moving thought could explain why there was only an effect in the stimulation block, this is unlikely because there was significantly more data in the stimulation block for detecting differences between the stimulation groups. Thus, these findings provide preliminary evidence for differential effects of HD-tDCS across regions for freely moving thought, with an increase in the PFC and a decrease in the VC.

As we predicted, anodal stimulation was also found to reduced deliberately constrained thought in the IPL, which aligns with Kam et al. (2021) finding distinct electrophysiological signatures for different type thoughts. It is important to note that deliberately constrained thoughts are not a direct measure of task focus as they capture all goal direct thoughts, which can be both directed at the task or something unrelated. Furthermore, the results differed from Kam et al. (2021), as they found differences for the frontal ERPs and in this study the effects were found in the parietal lobule. This contrasting result could be due to the use of distinct methodological practices, such as contrasting task and study designs. For example, EEG has been recognised as having weak spatial resolution, which makes the localisation of the electrical activation highlighted by the frontal effects difficult (Michel et al., 2004). Alternatively, the stimulation targeting the parietal network may have also indirectly affected PFC activity, as there is evidence for the frontoparietal control network being recruited during internal thought processes (Christoff et al., 2009; Groot et al., 2021). However, the IPL specific results are strengthened with the use of an active control region, as it ensured the distinct PFC and IPL effects between the dynamic thought types were specific to the region of interest.

The changes in randomness scores and behavioural variability between on- and off-task thoughts of freely moving thought also suggest a relationship between internal thoughts and the recruitment of executive functioning resources. Specifically, shifts towards periods of freely moving thought reduce randomness and increase individuals’ behavioural variability, thus limiting the executive resources required to stay on task. This is consistent with the relationship found by Boayue et al. (2021), and supports evidence for the overlap in the brain regions recruited during executive control processes and mind wandering (Christoff et al., 2009; Fox et al., 2015). However, there was no effect of stimulation on the measures of task performance across the thought probes, which did not align with the hypothesised reduction in behavioural variability and increase in randomness for the anodal stimulation groups in the PFC and IPL. There is previous literature which has also found limited evidence of stimulation on task performance measures in the context of mind wandering, with some task effect being dependent on the stimulation intensity (Filmer et al., 2021). This suggests the null effect may also be due to methodological factors which were not manipulated in the current research.

In summary, this study has highlighted the role of distinct neural substrates in internal thought processes. We have shown that anodal stimulation reduces freely moving thought in the left PFC and reduces deliberately constrained thought in the right IPL, with preliminary evidence for an increase in freely moving thoughts in the VC also. By understanding the importance of distinct types of dynamic thought, across different brain regions, this will enable researchers to target brain activity in a context dependent manner, to improve executive functioning and cognitive control processes. Specifically, this may be used to improve performance in contexts which require sustained attention and may also have clinical applications in being able to target pervasive negative thoughts in distinct brain regions. Collectively, this research provides evidence for HD-tDCS being an effective tool to modulate and reduce dynamic thought types across the brain.

## Appendix A tDCS Safety Screening Questionnaire

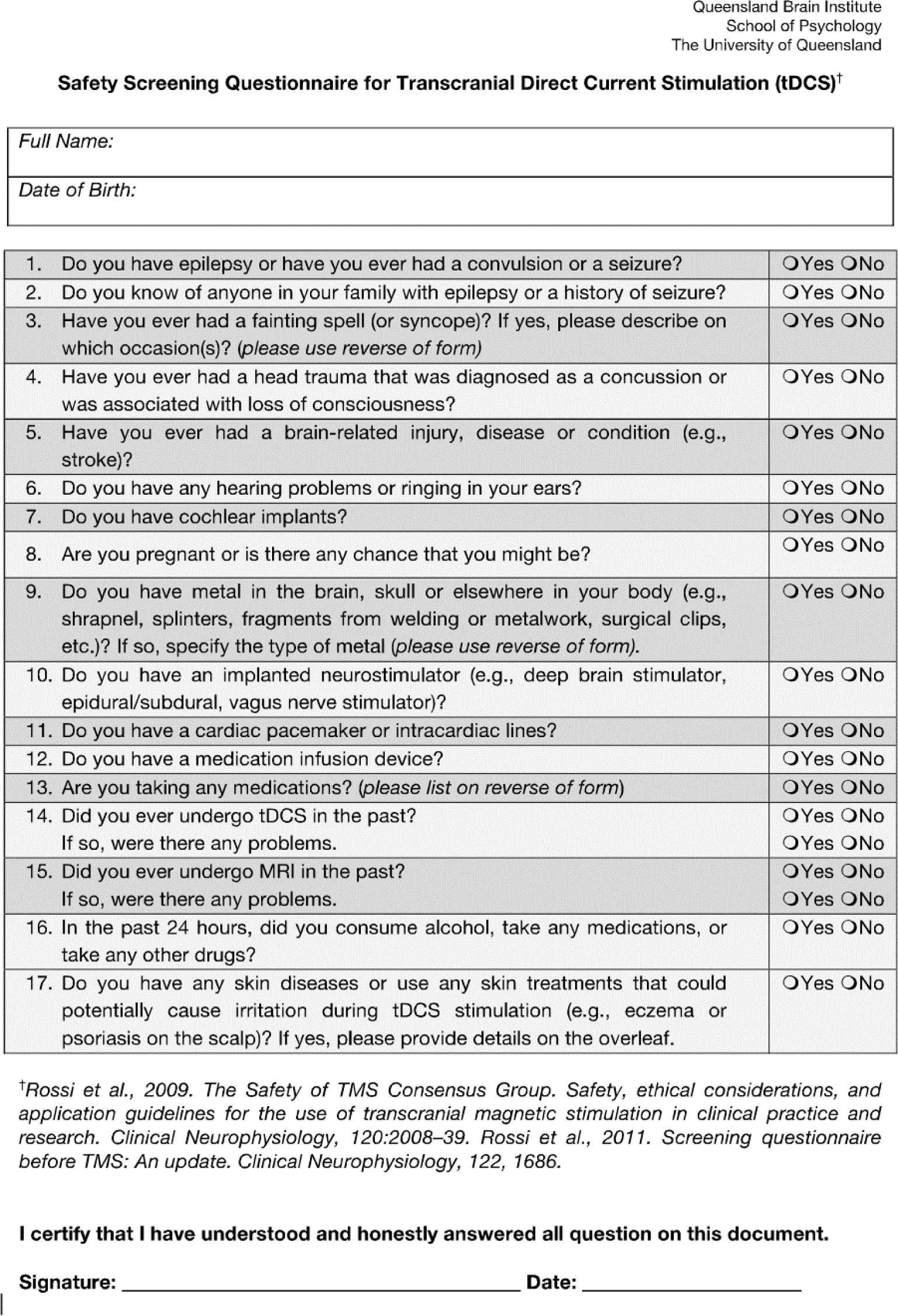
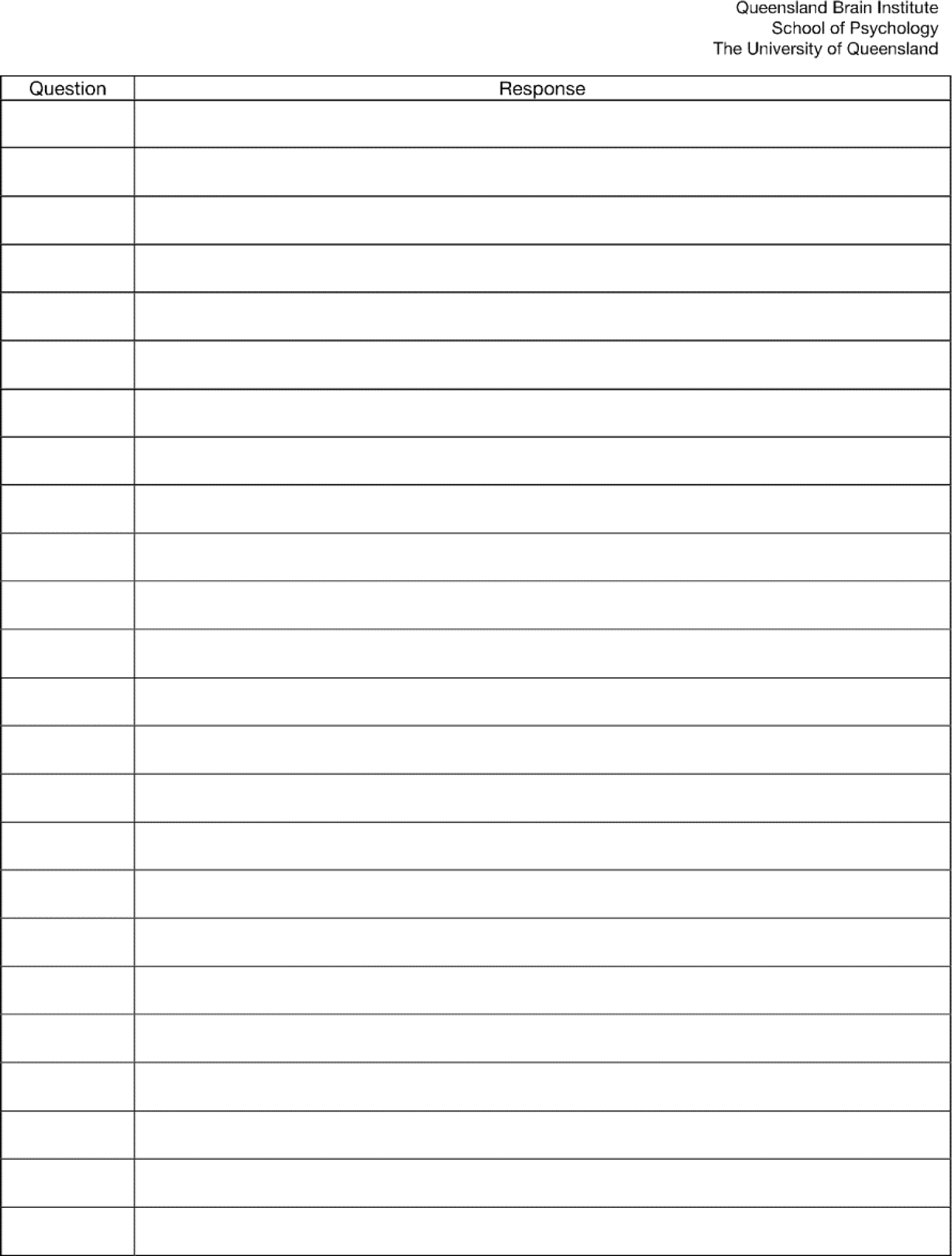

## Appendix B Dynamic Thought Probe Descriptions

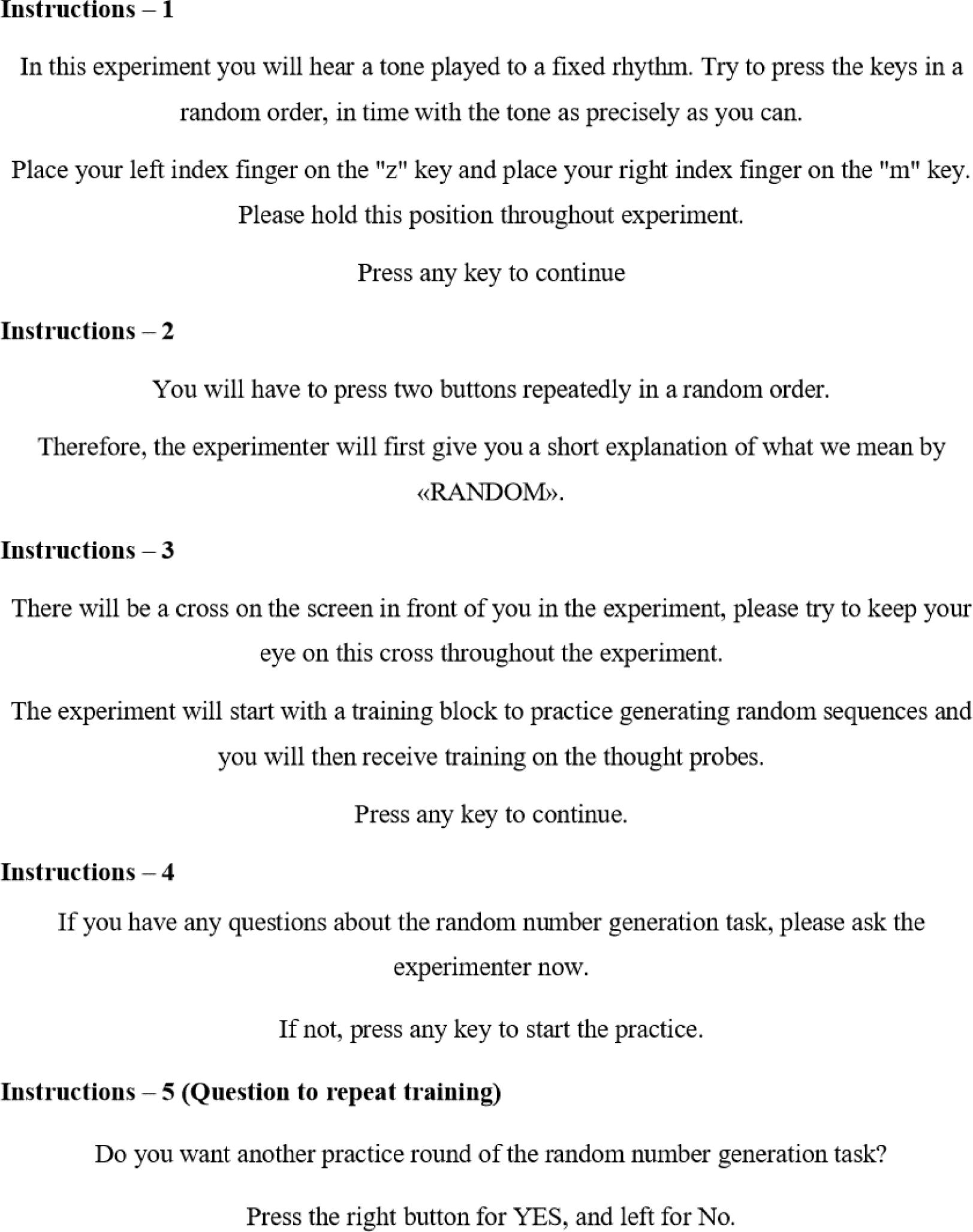
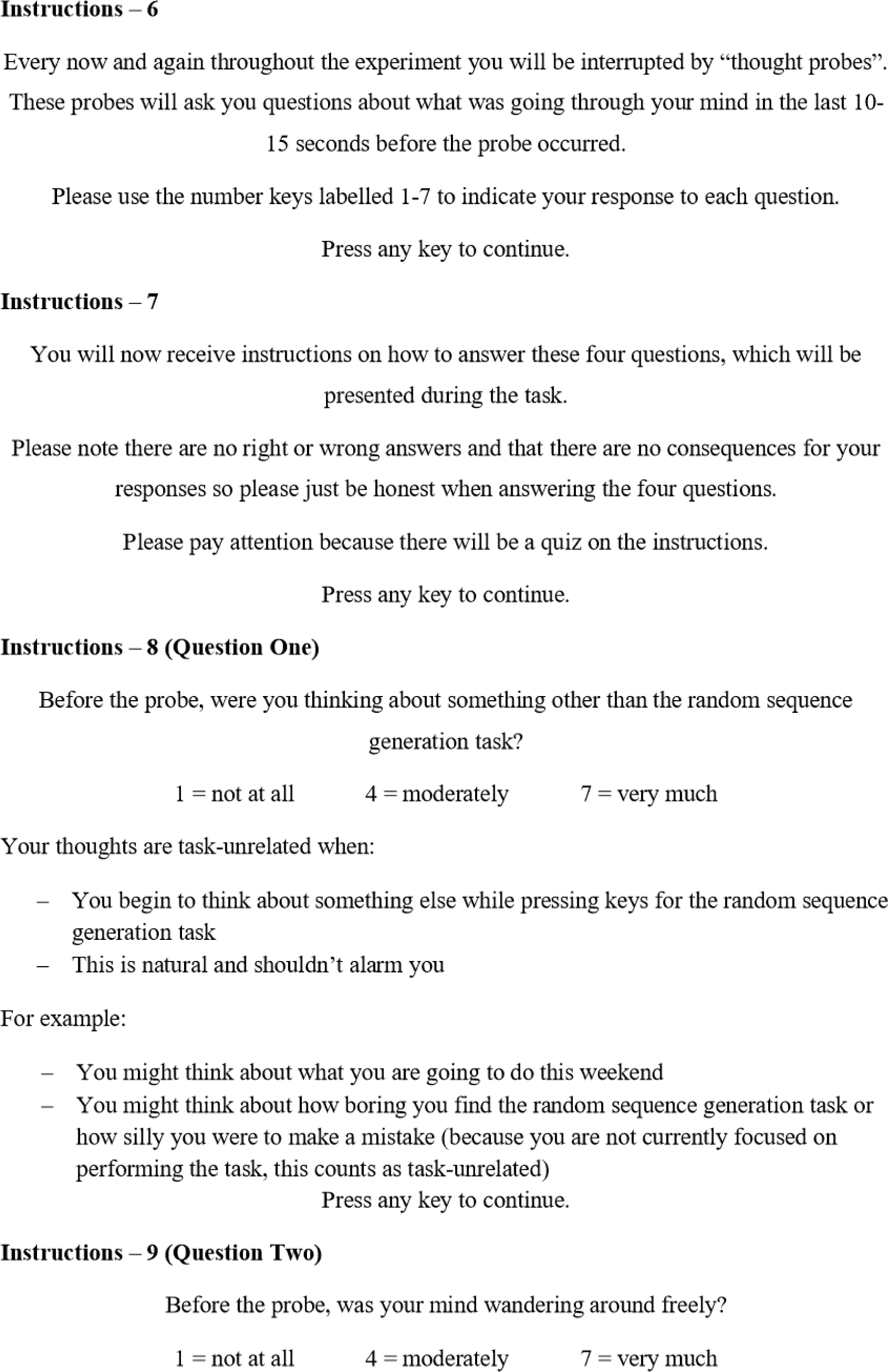
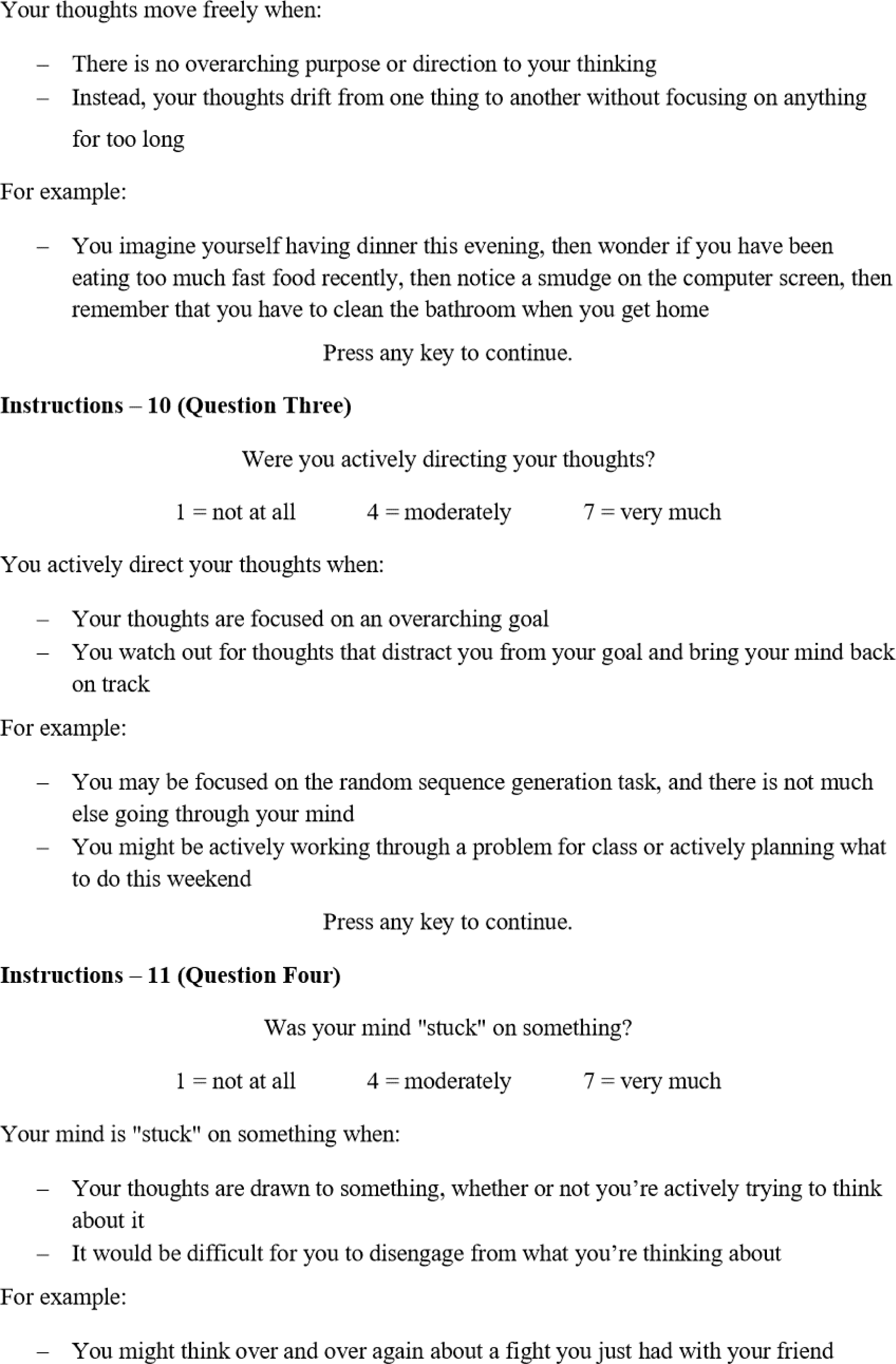
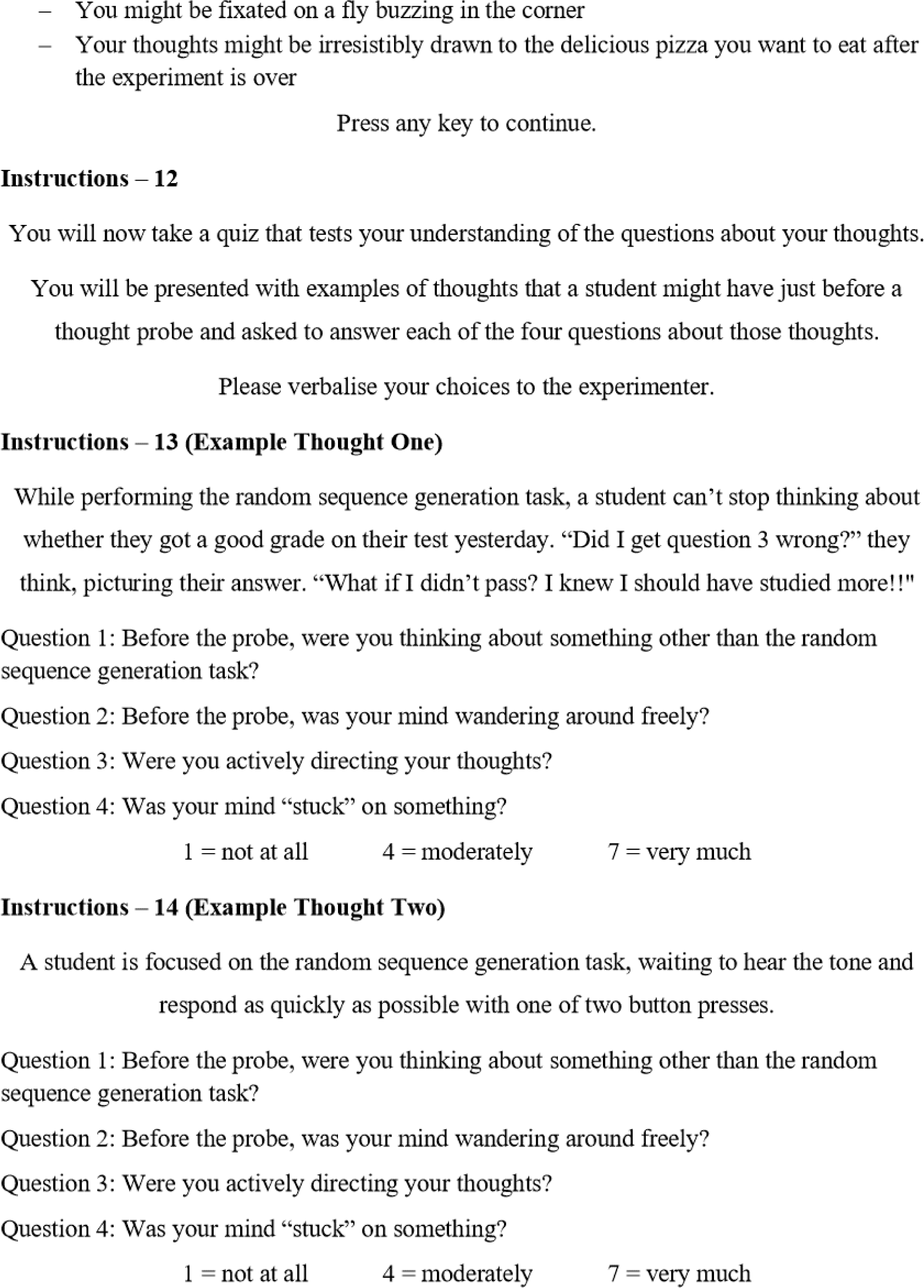
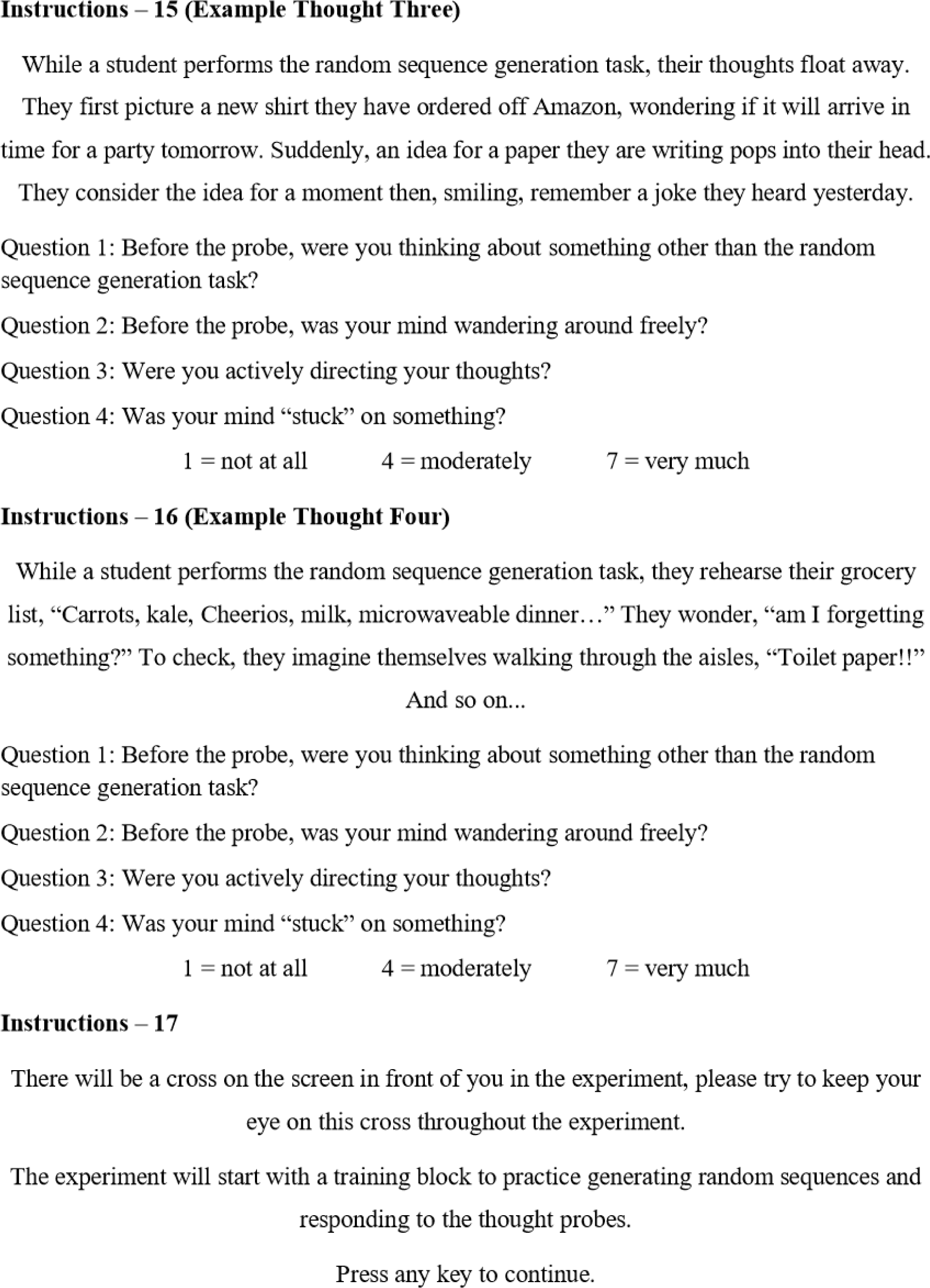
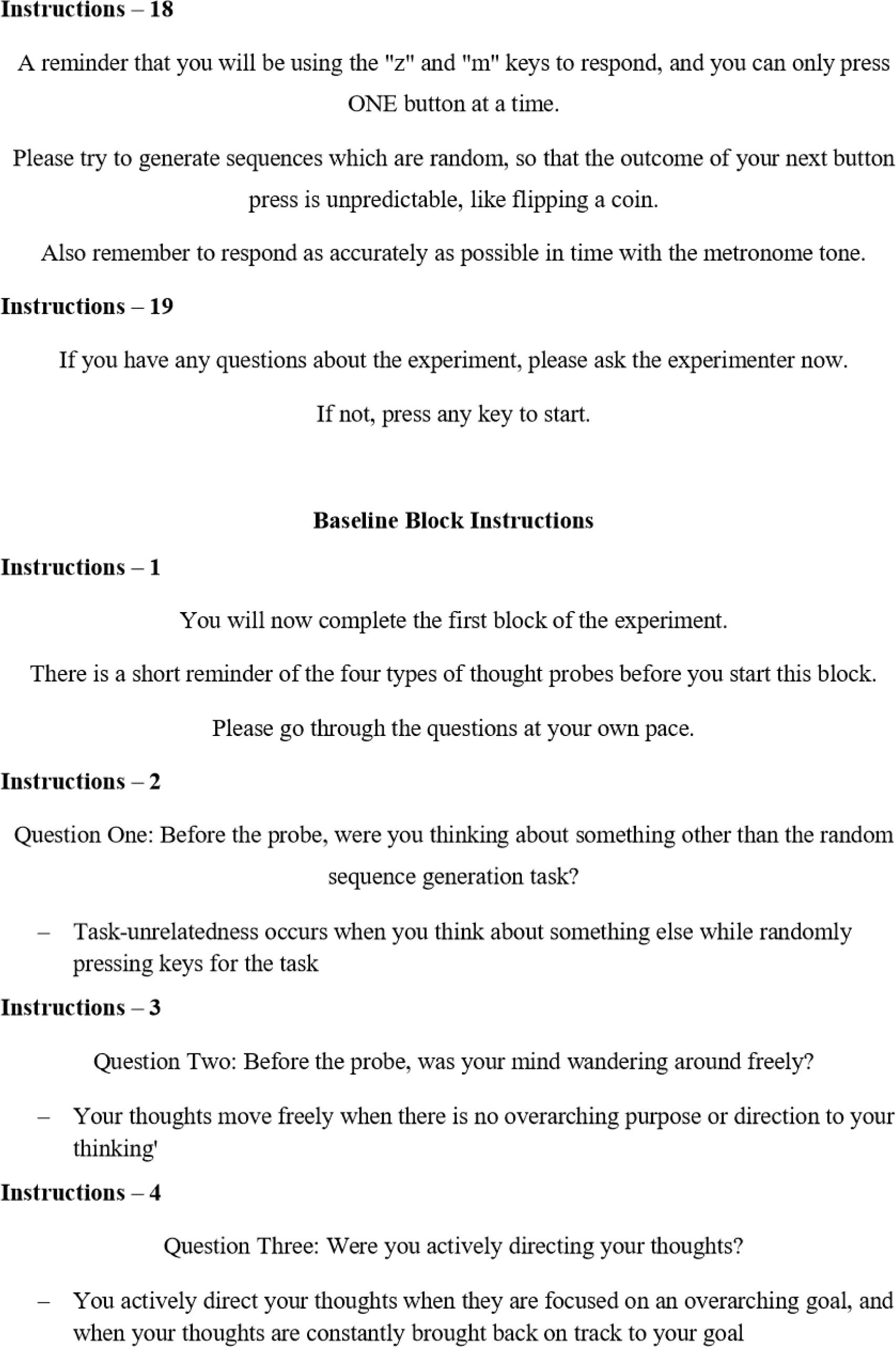
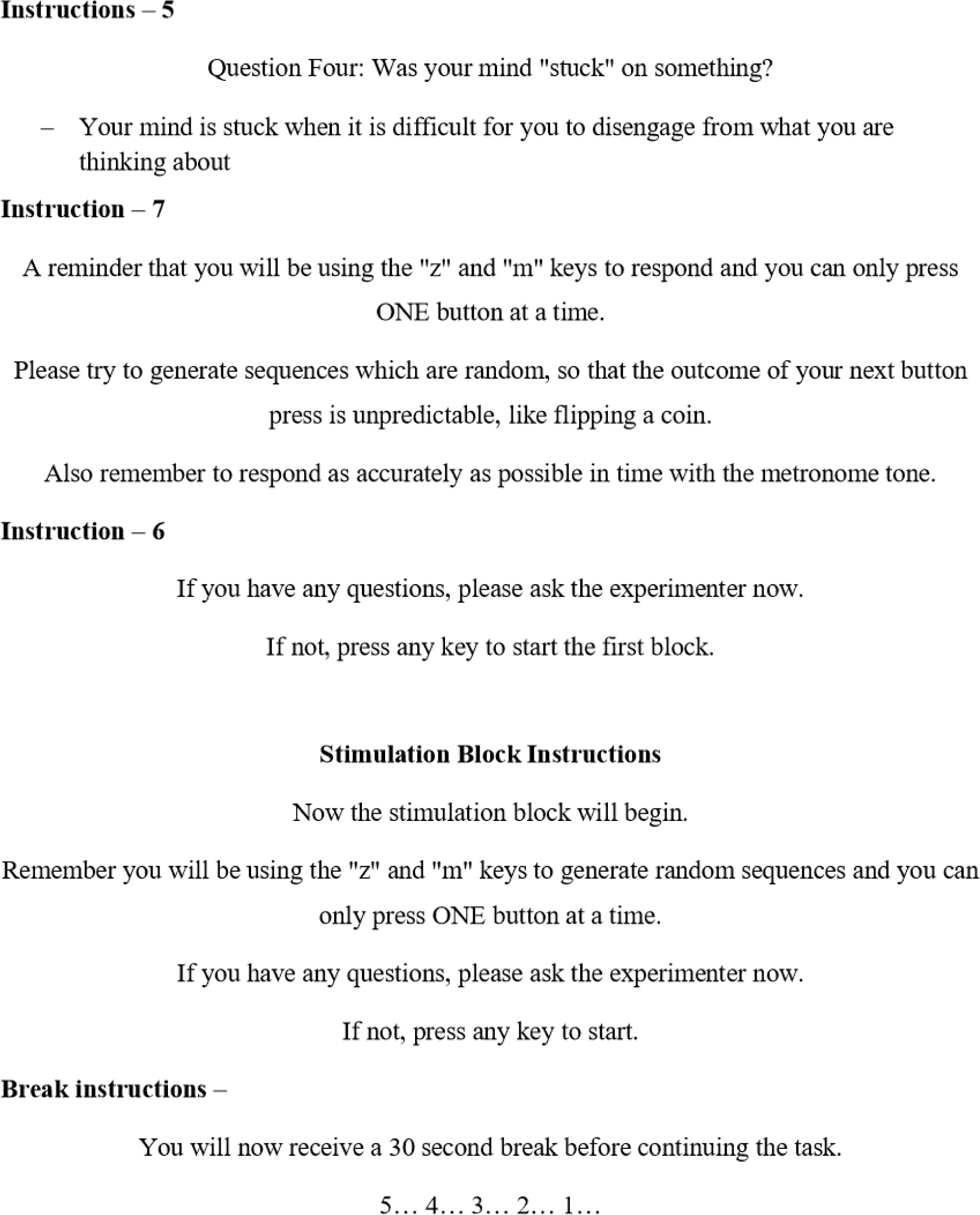

## Appendix C Thought Probe Likert Scales

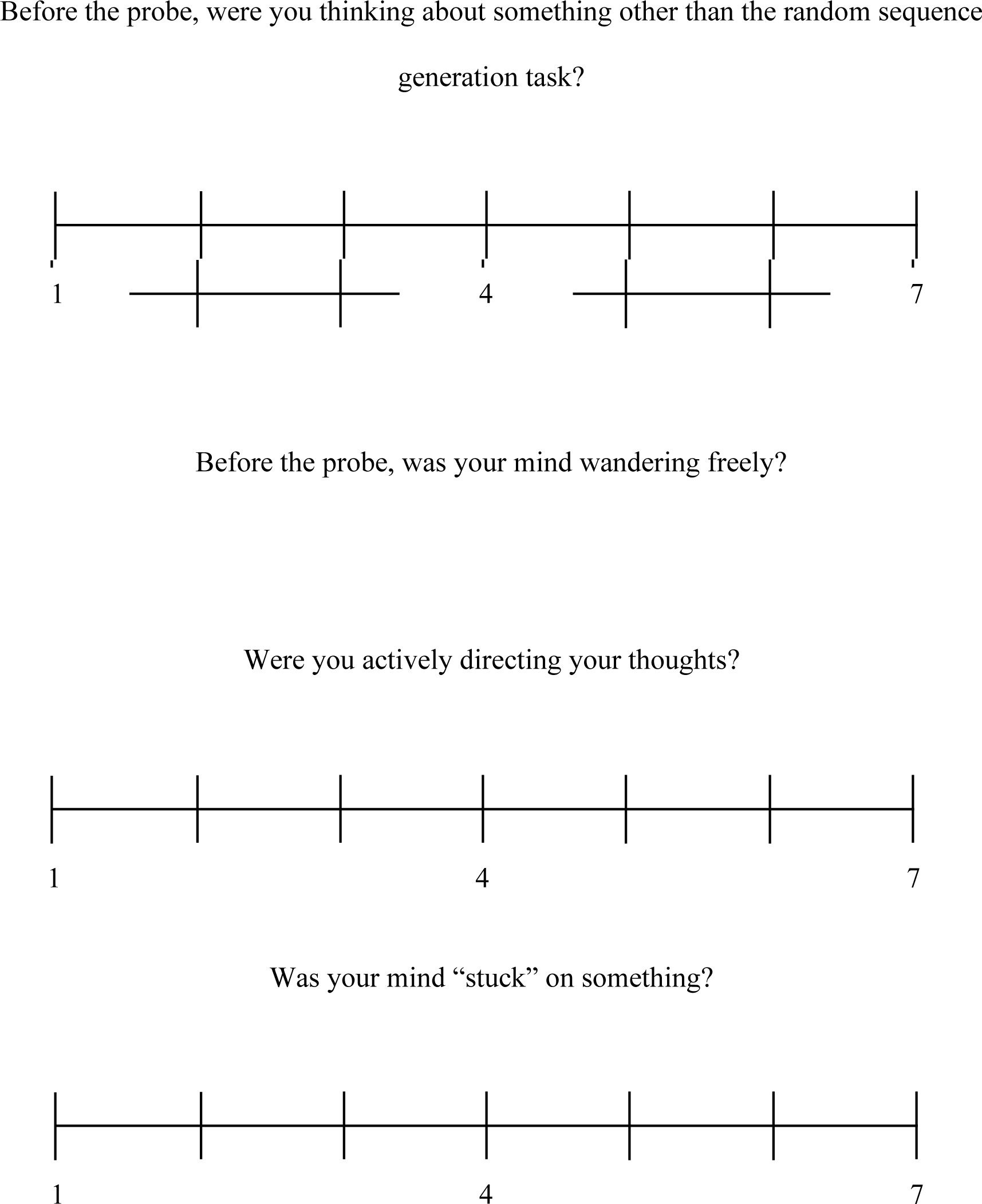

## Appendix D Pre-Task Questions

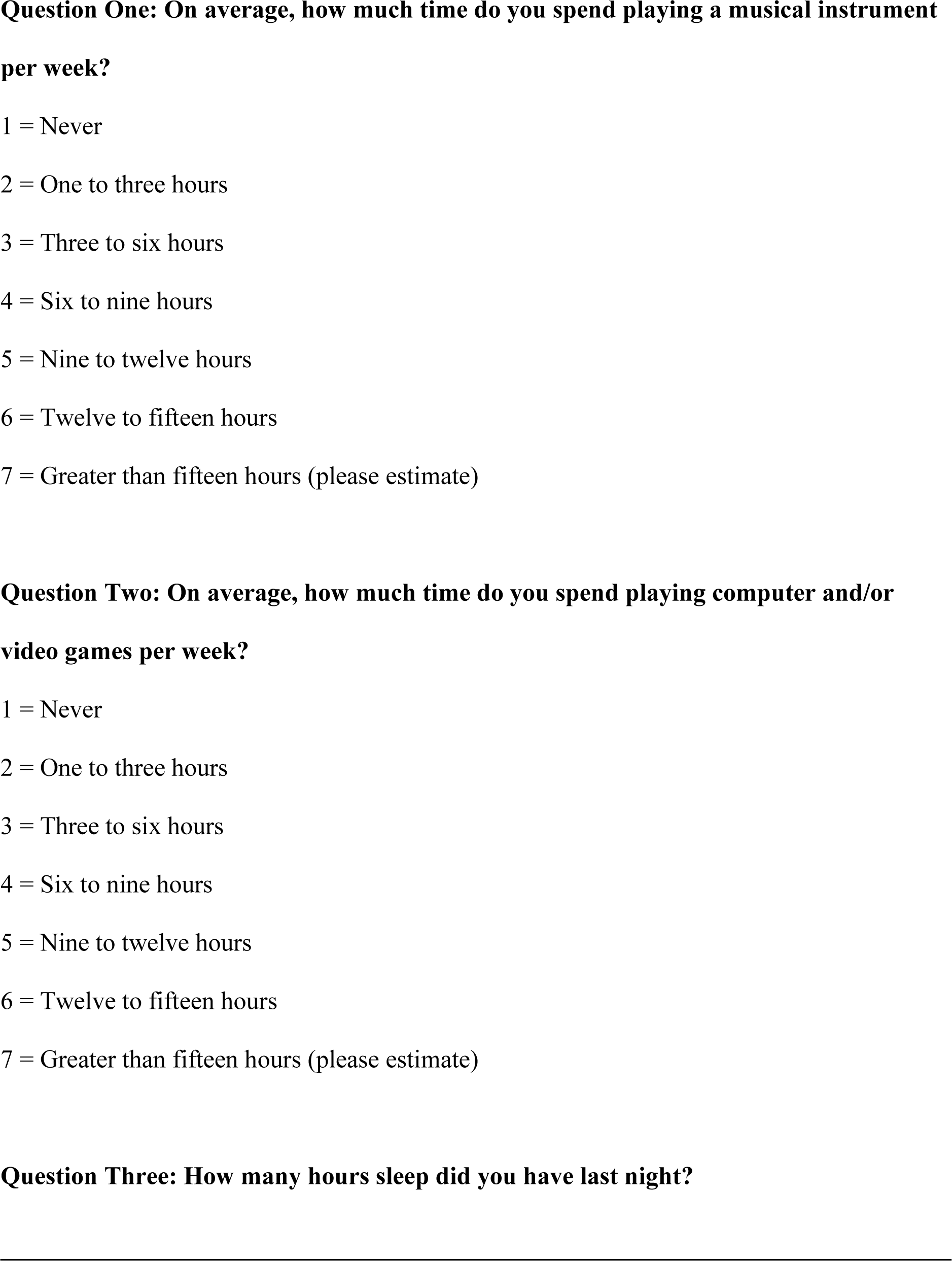

## Appendix E MAAS Questionnaire

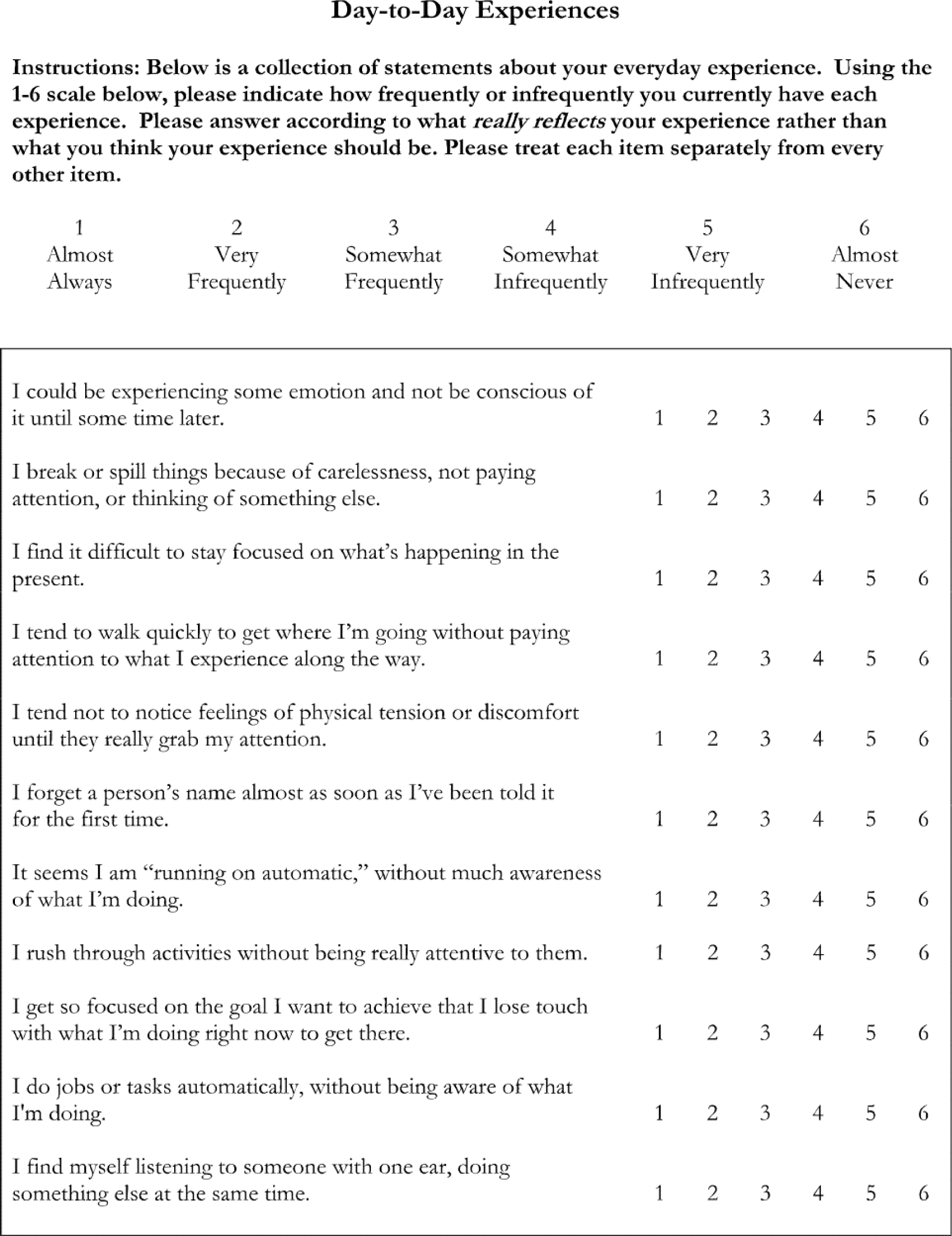
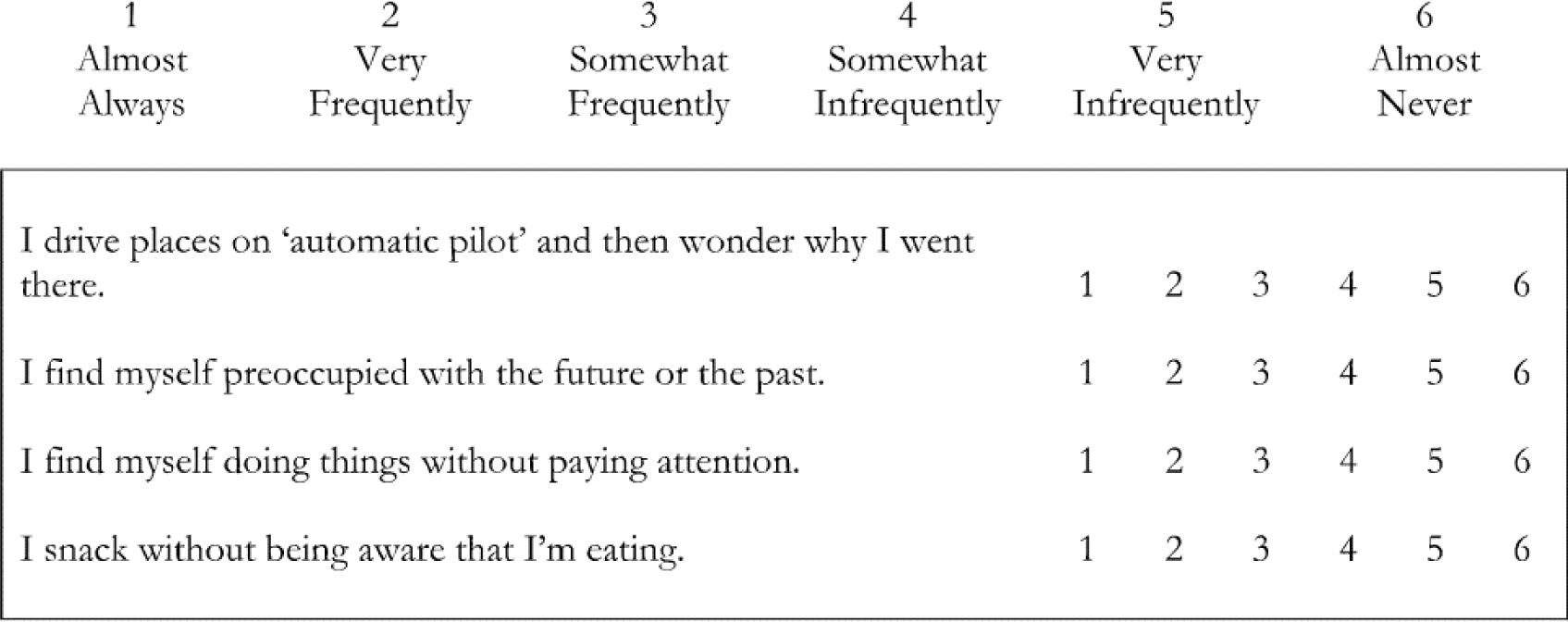
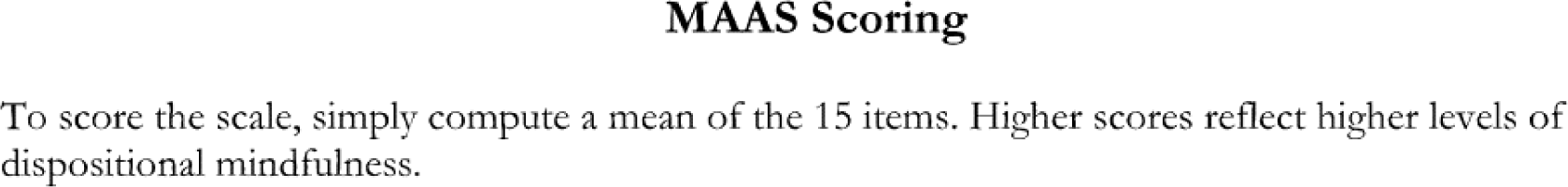

## Appendix F Adult ADHD Self-Report Scale

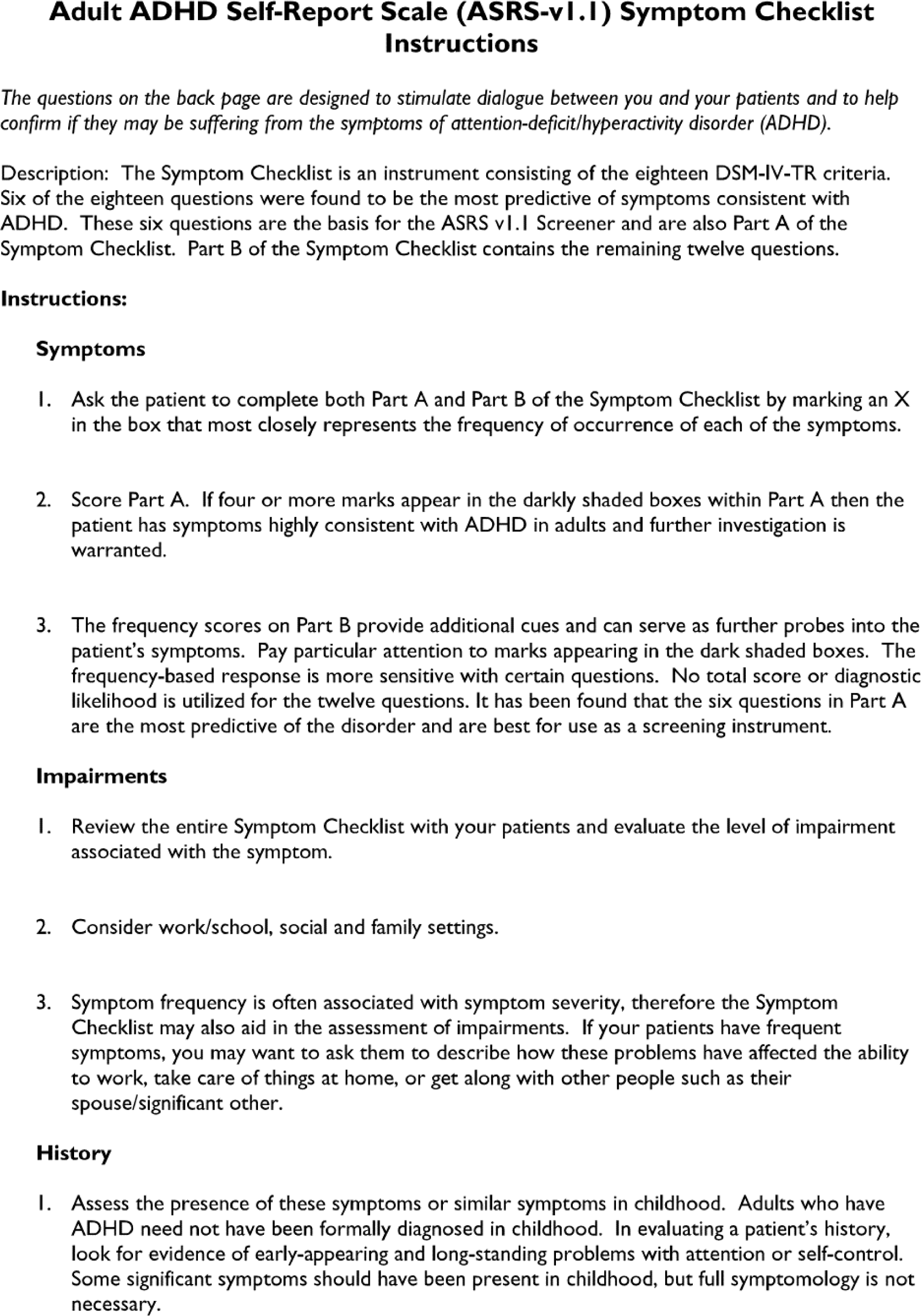
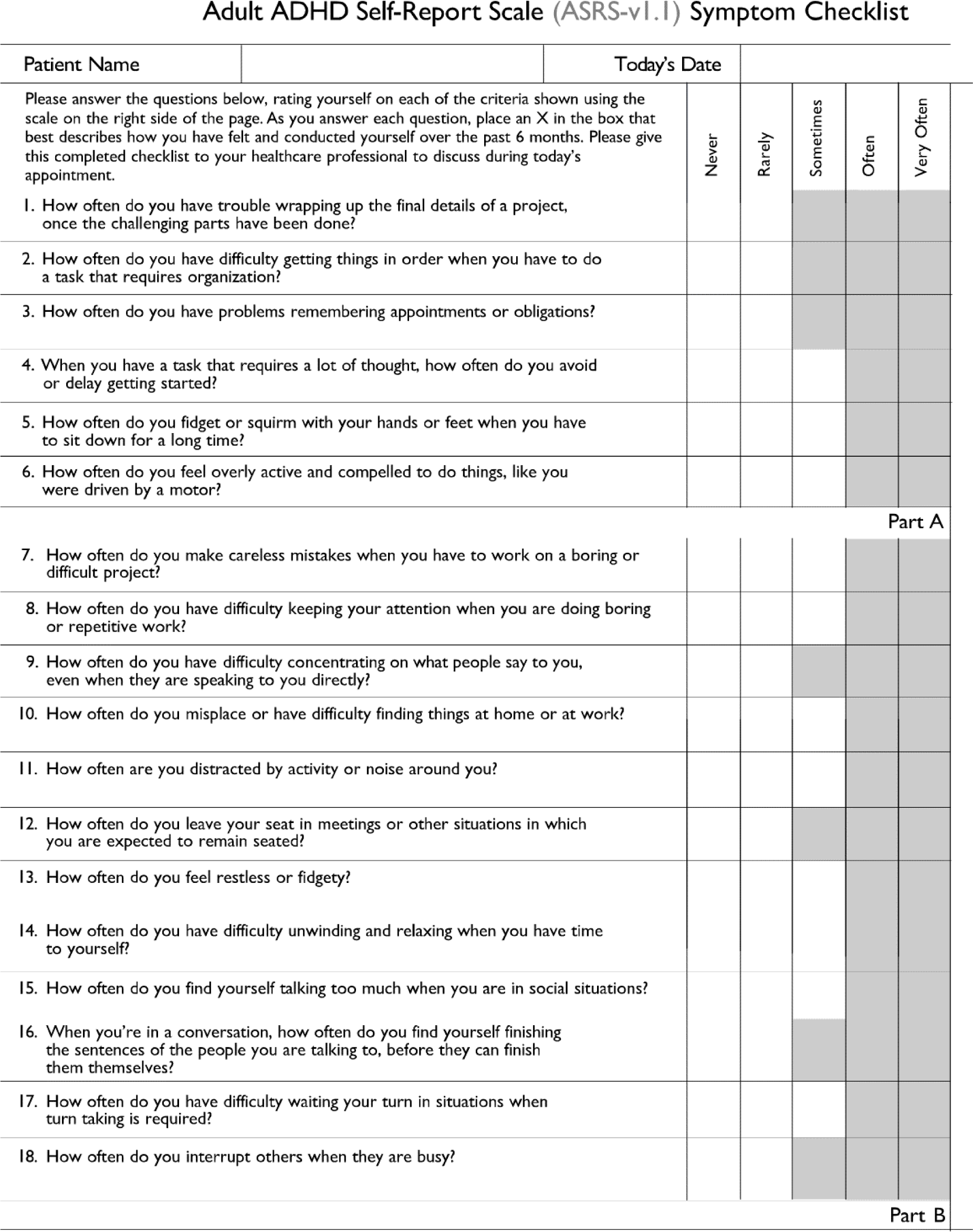

## Appendix G The Rumination Response Scale

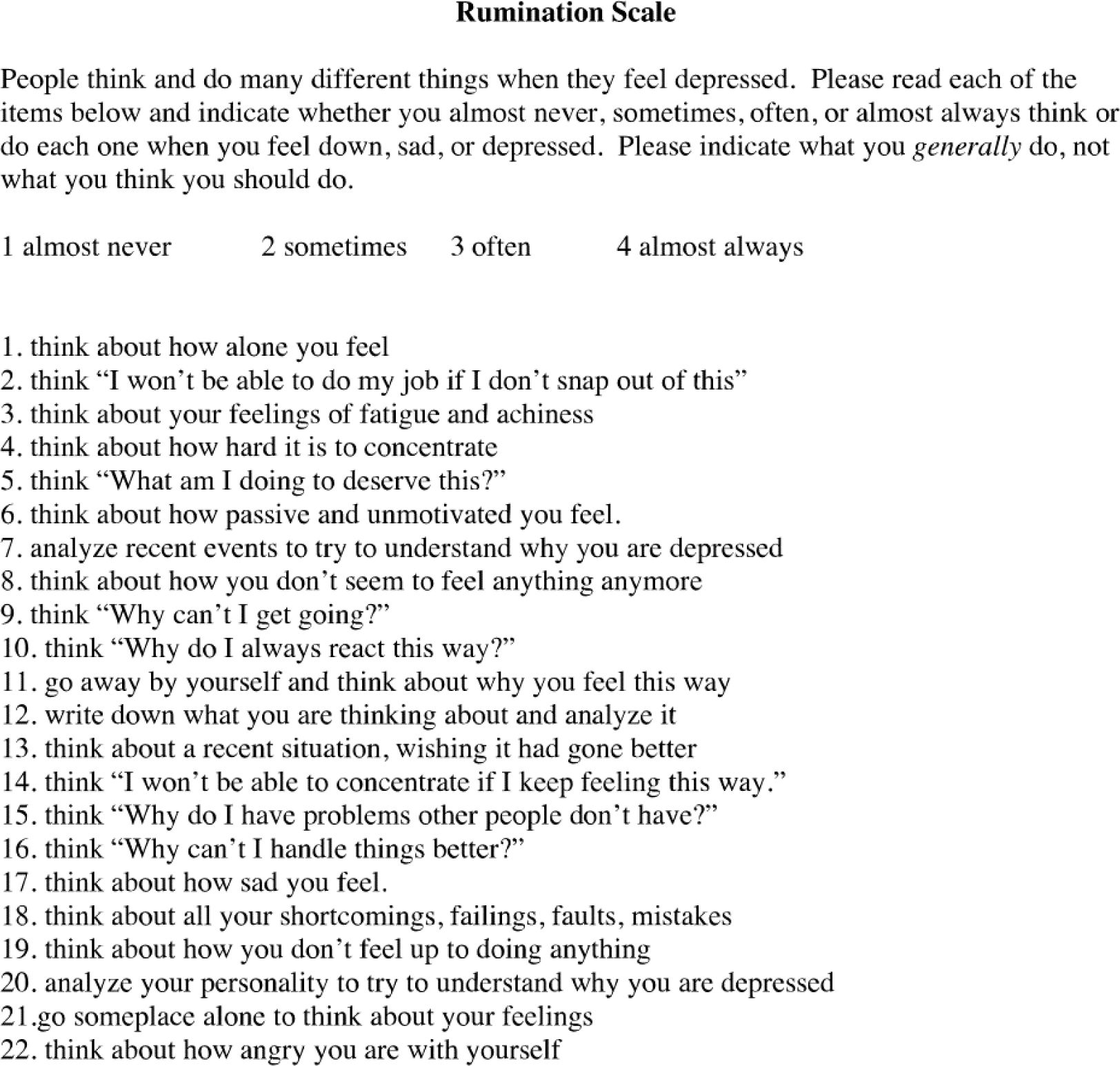

## Appendix H Post-Task Questionnaire

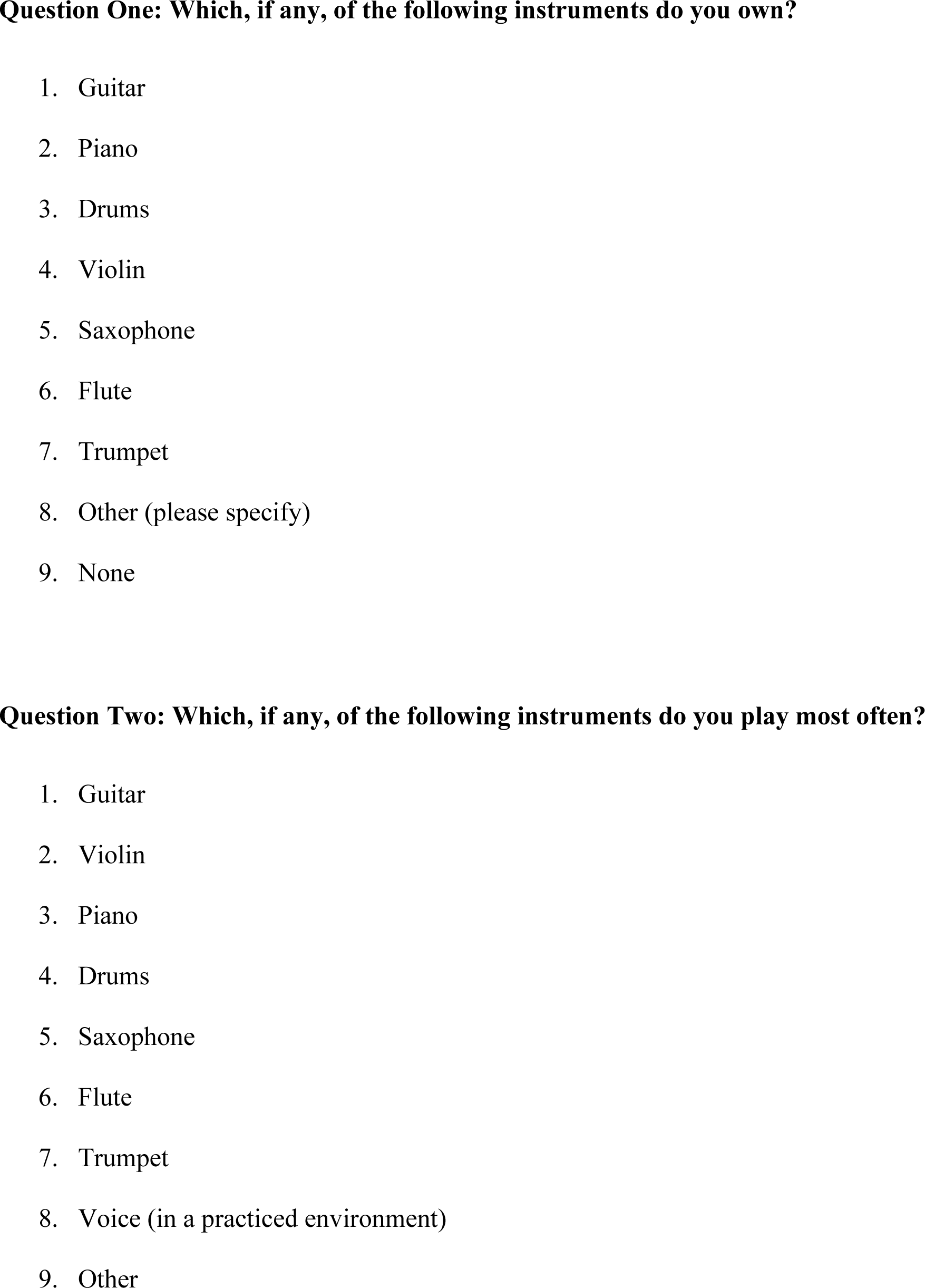
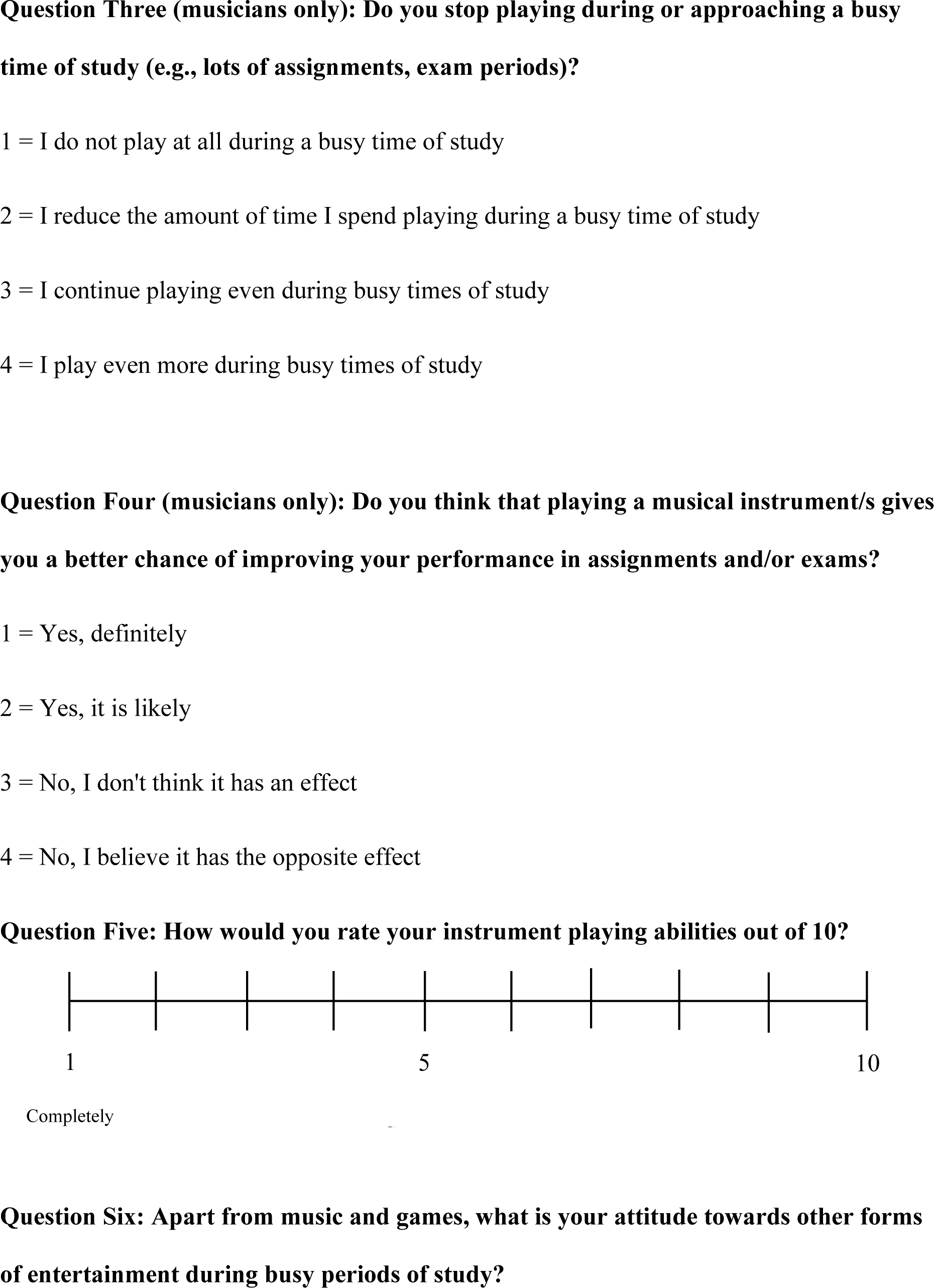
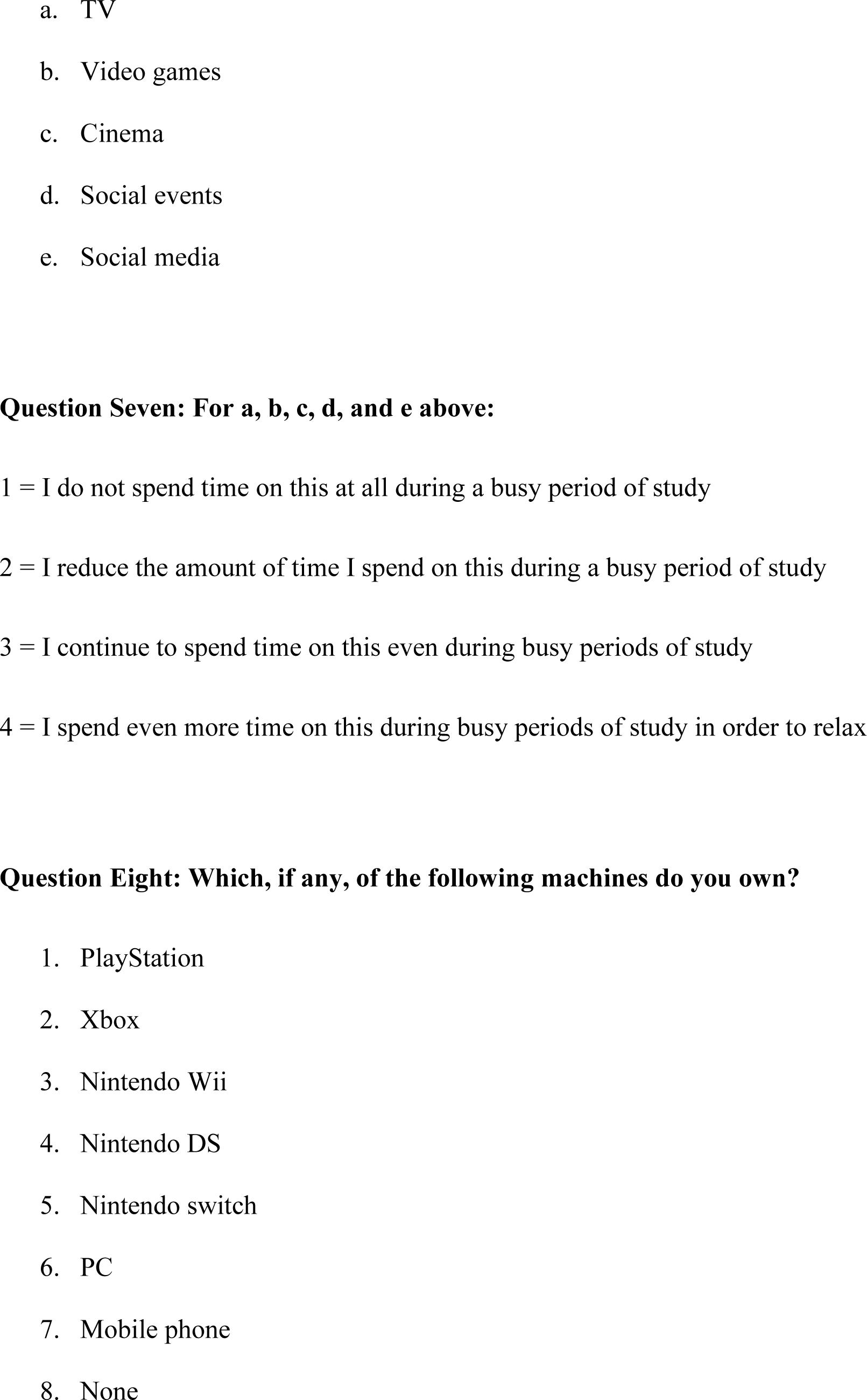
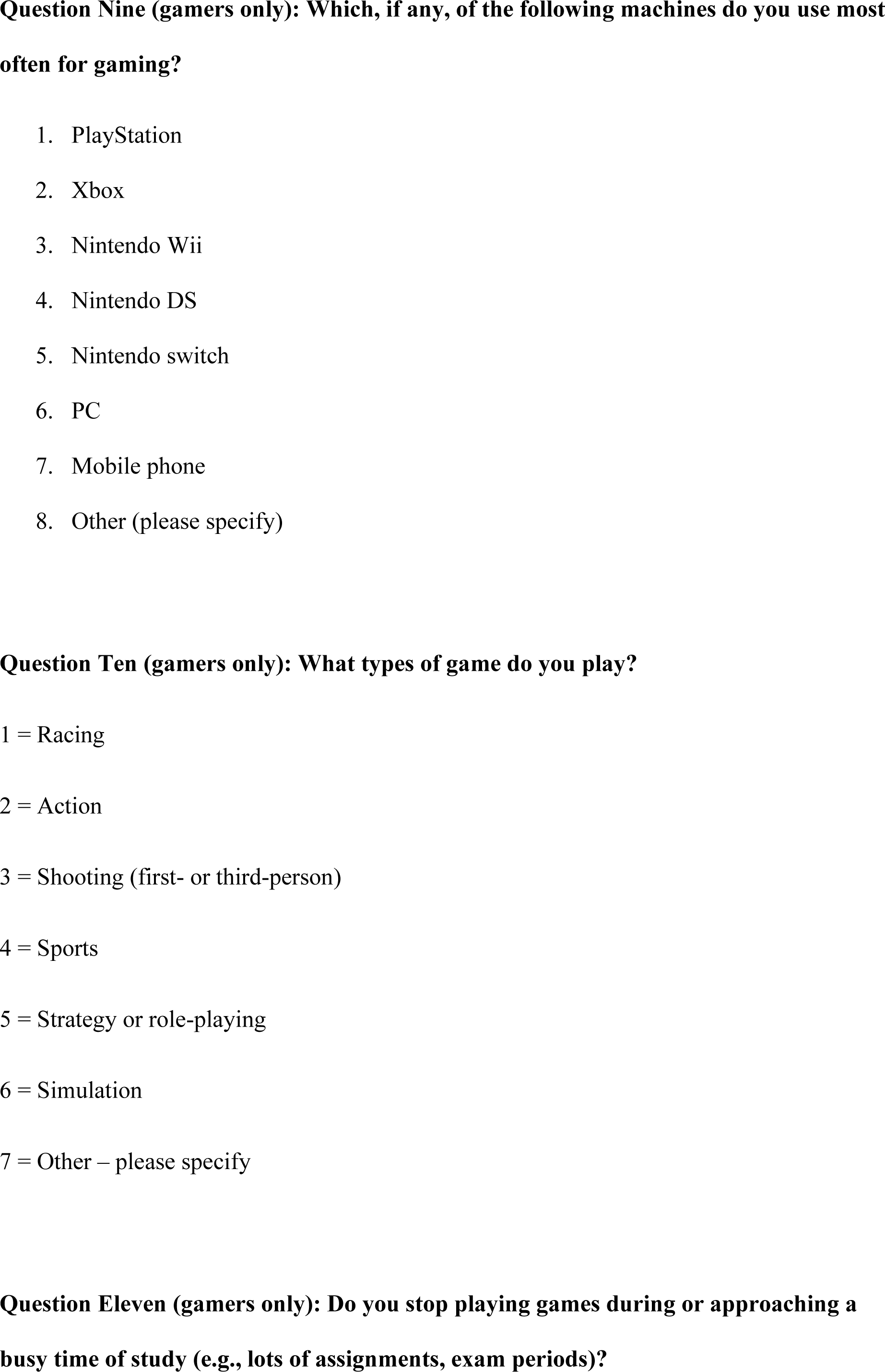
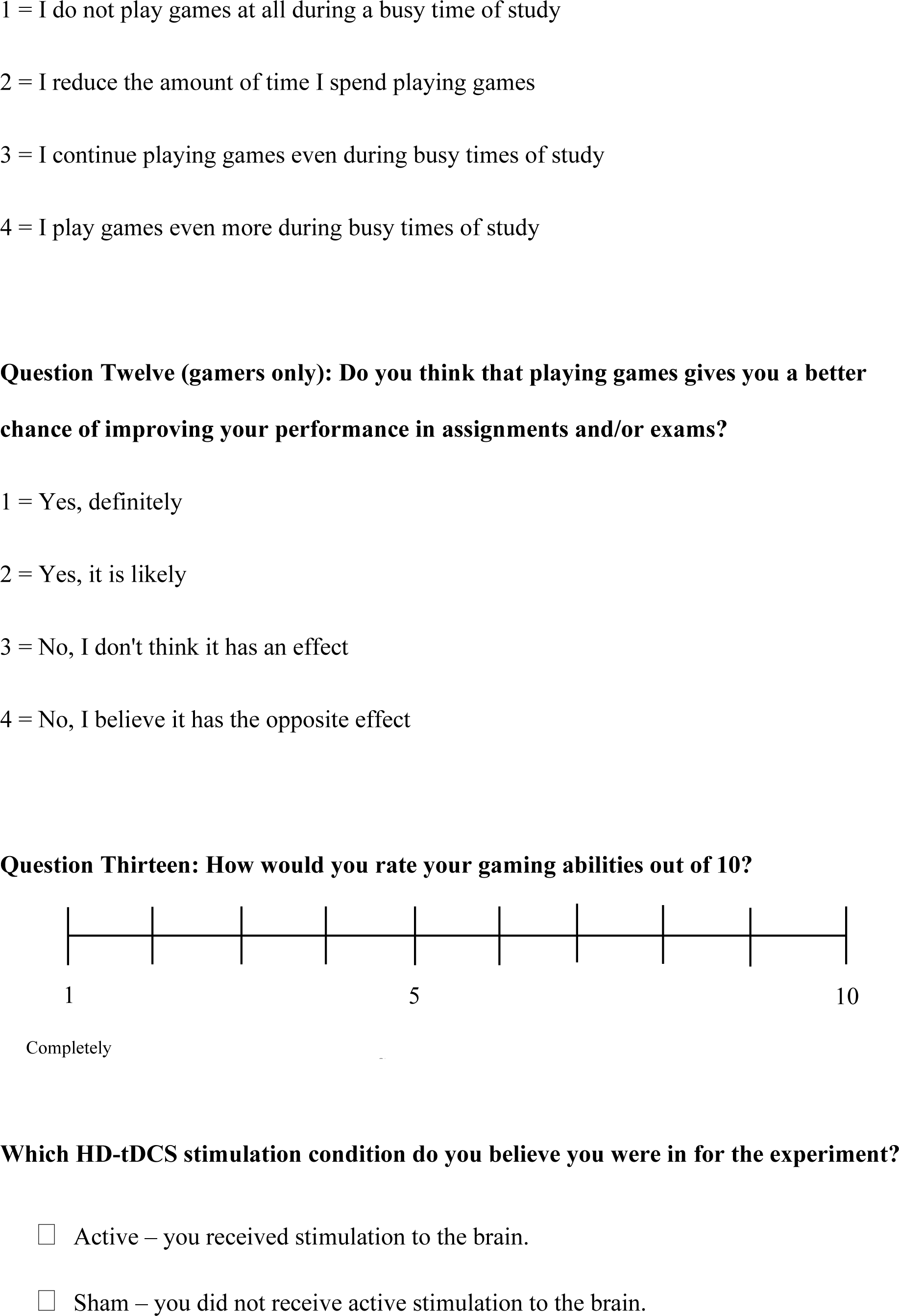
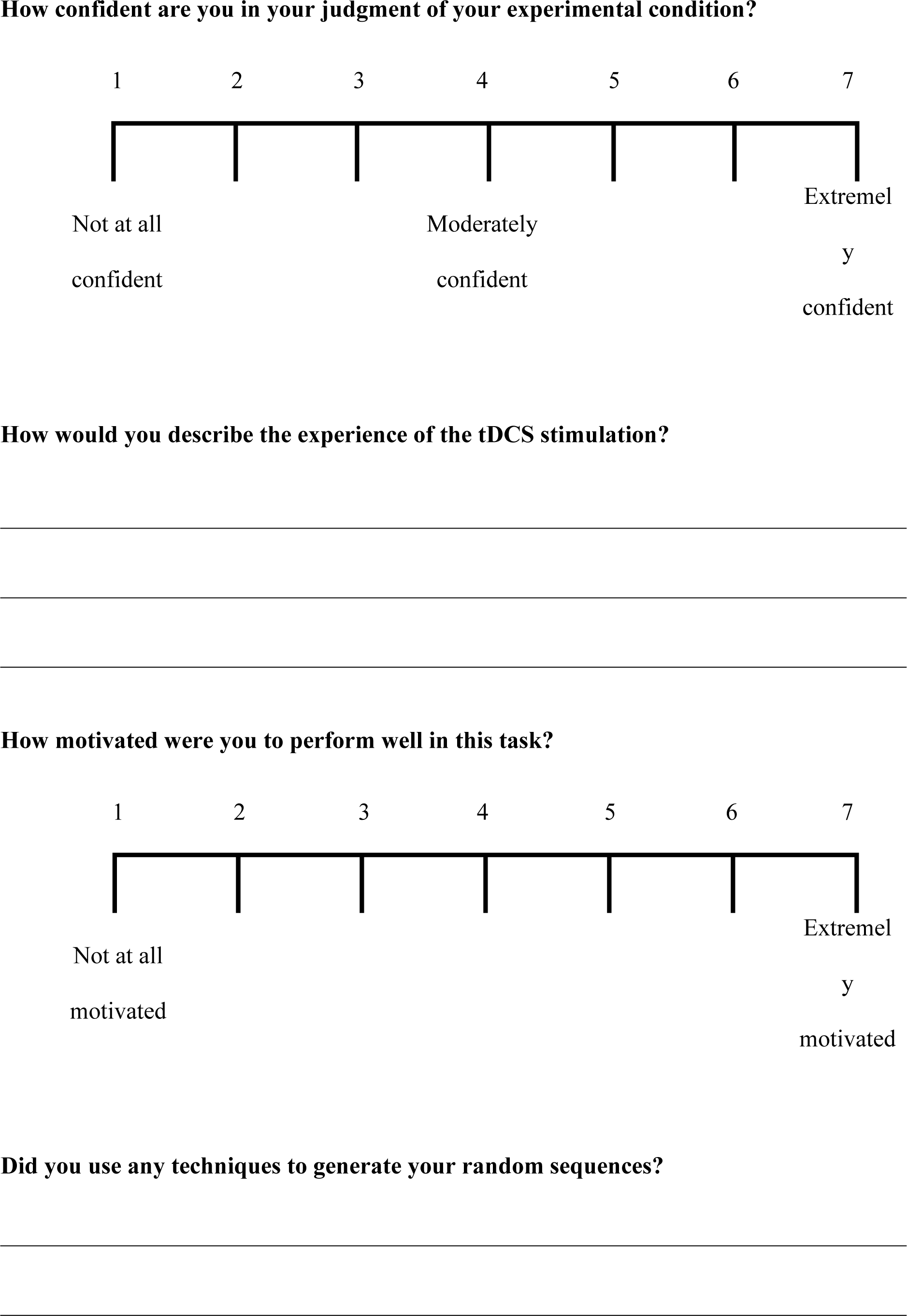
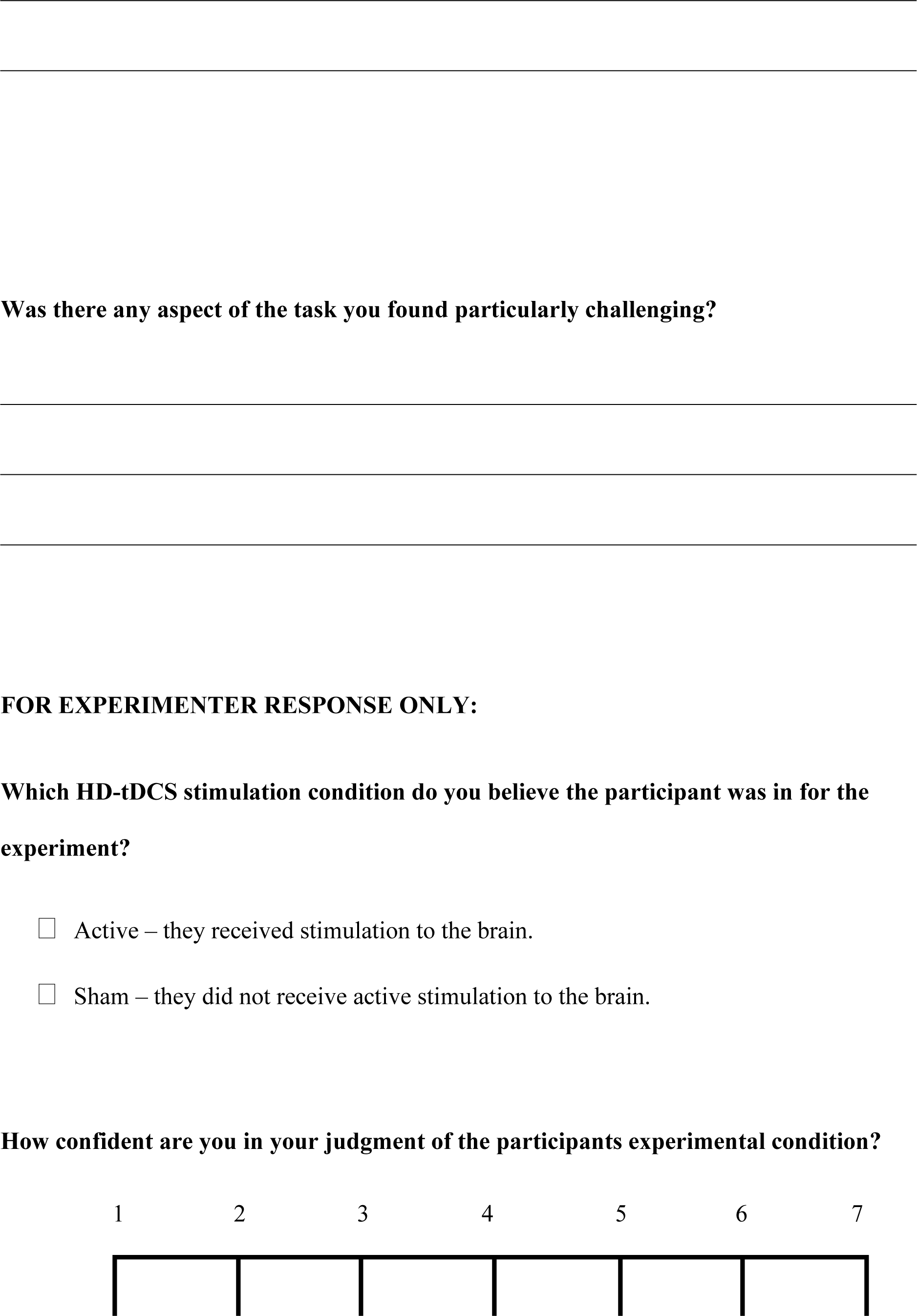
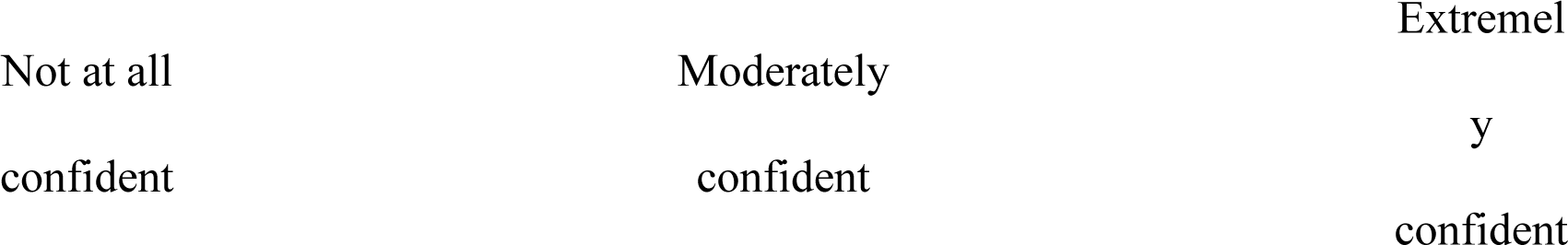

