## Supplementary Materials for "On the role of prefrontal and parietal cortices in mind wandering and dynamic thought"

#### Figure S1. Full Model Comparisons

A. PFC Task Unrelated Thought

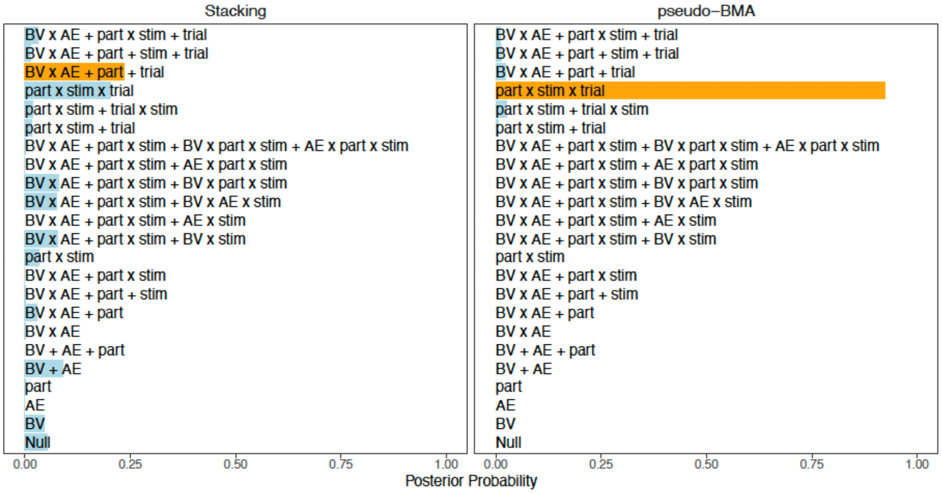


B. IPL Task Unrelated Thought

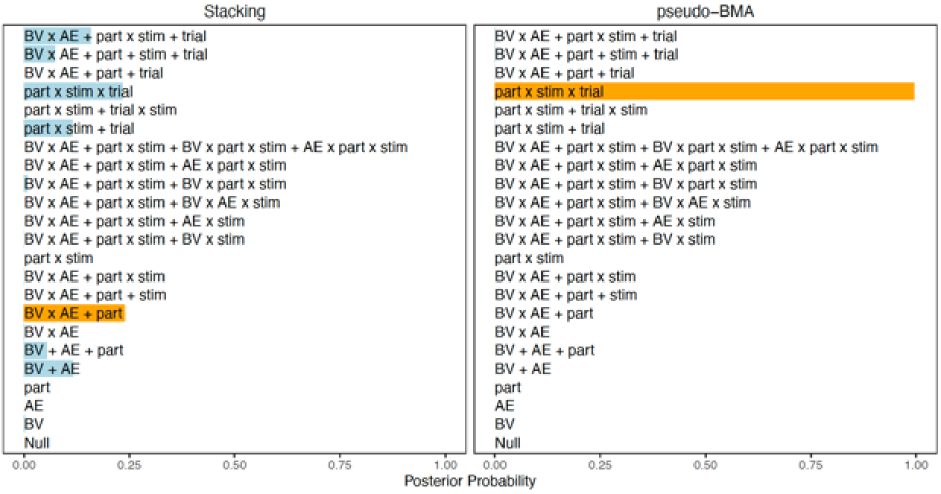


C. VC Task Unrelated Thought

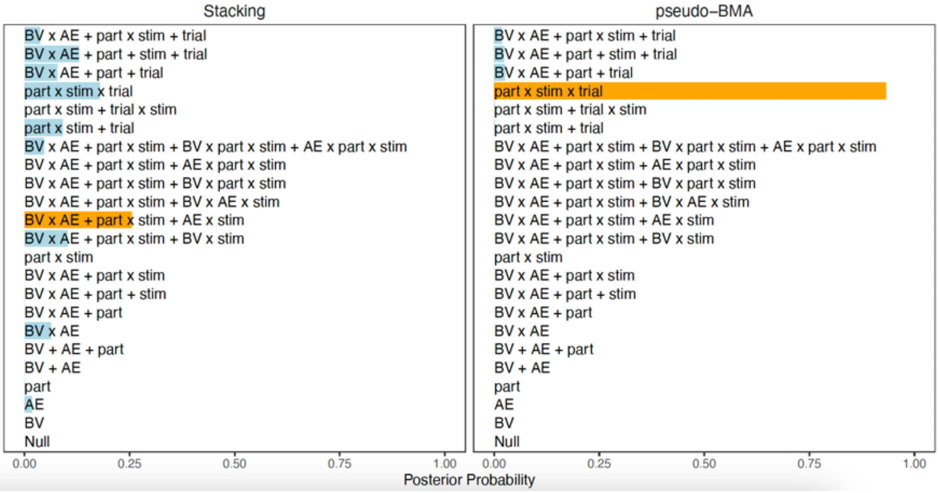

D. PFC Freely Moving Thought

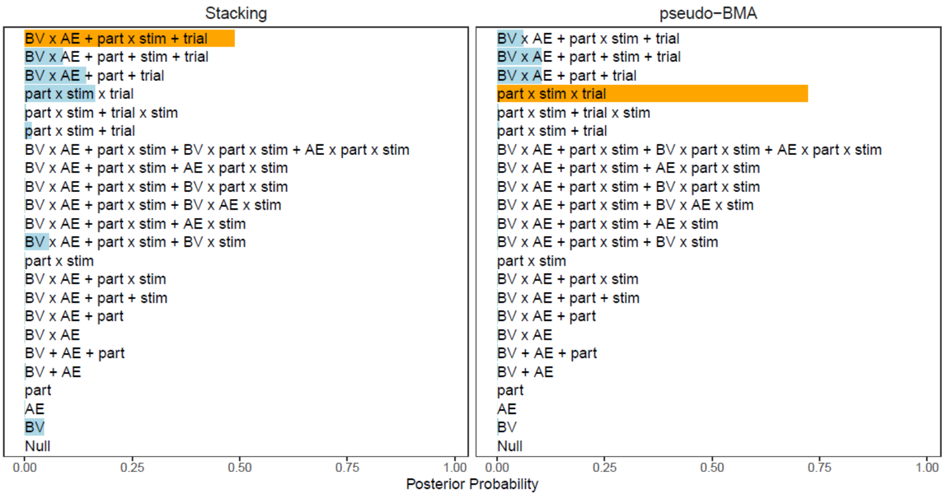


E. IPL Freely Moving Thought

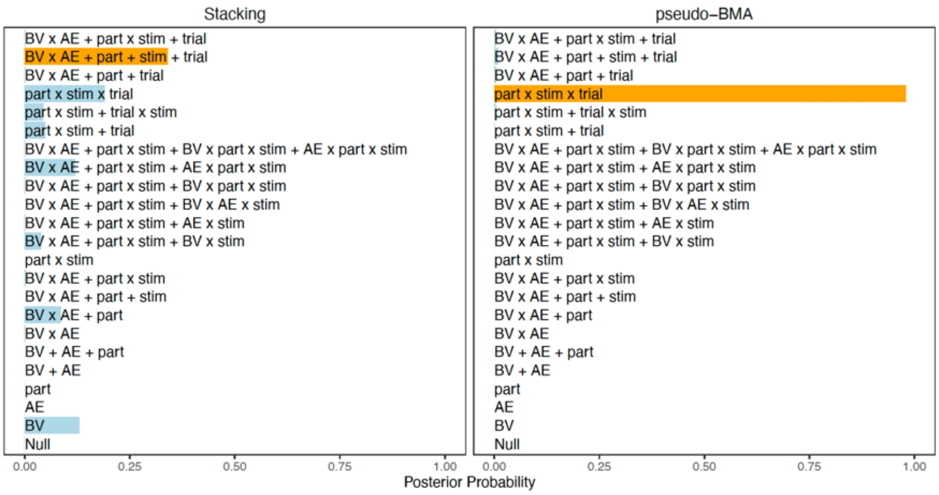


F. VC Freely Moving Thought

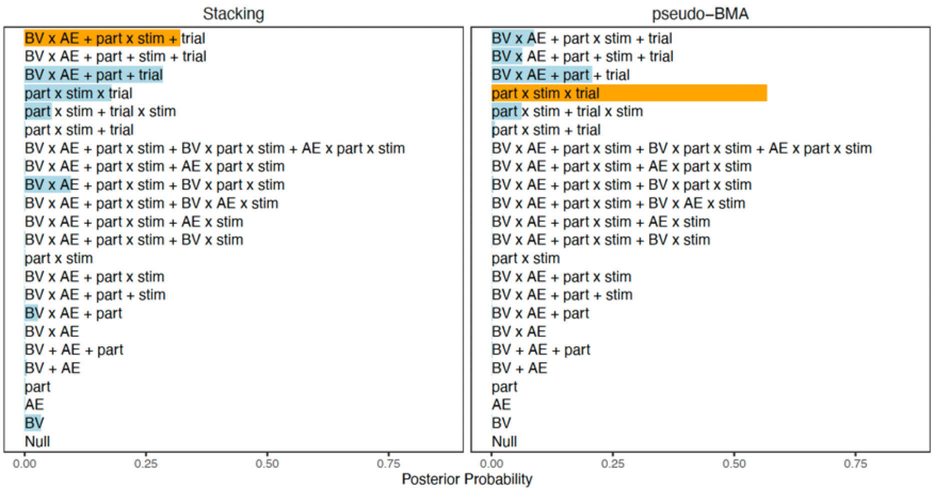

### G. PFC Deliberately Constrained Thought

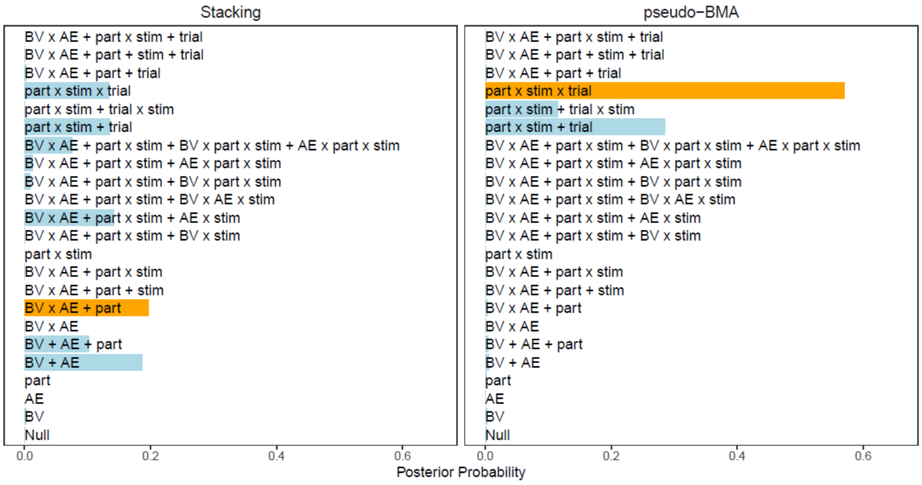


### H. IPL Deliberately Constrained Thought

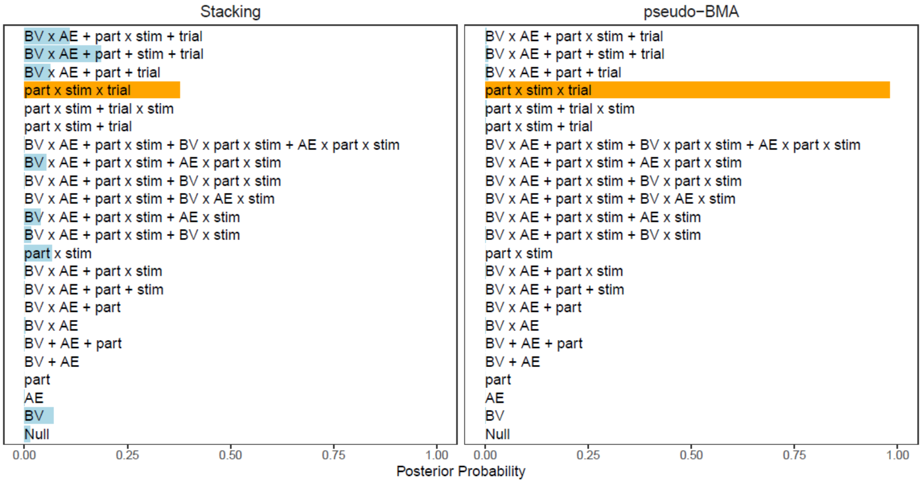


### I. VC Deliberately Constrained Thought

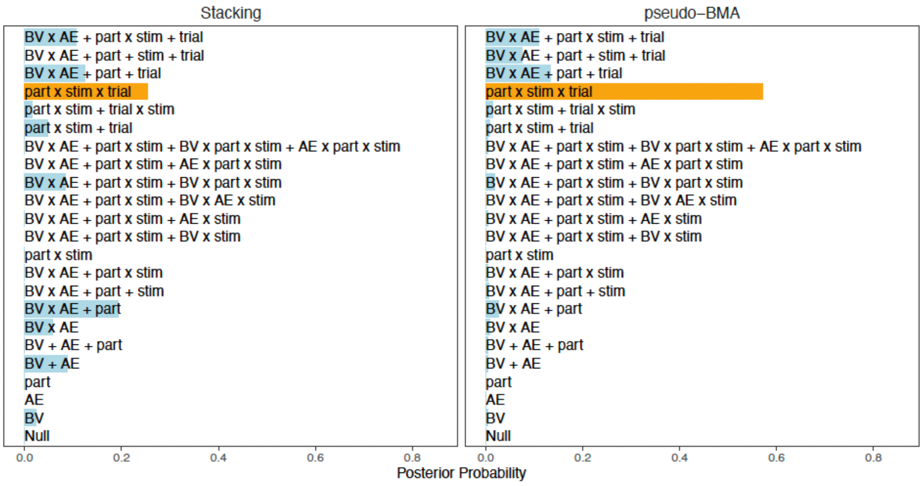

#### J. PFC Automatically Constrained Thought

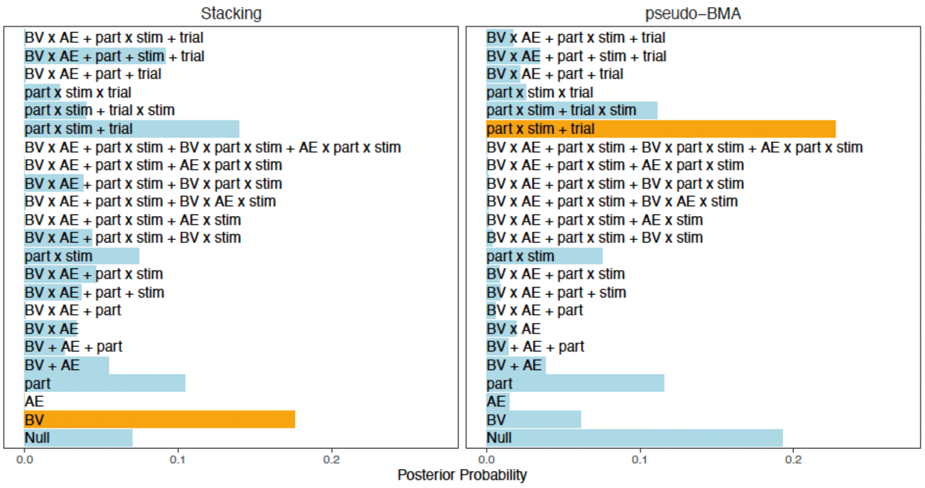


#### K. IPL Automatically Constrained Thought

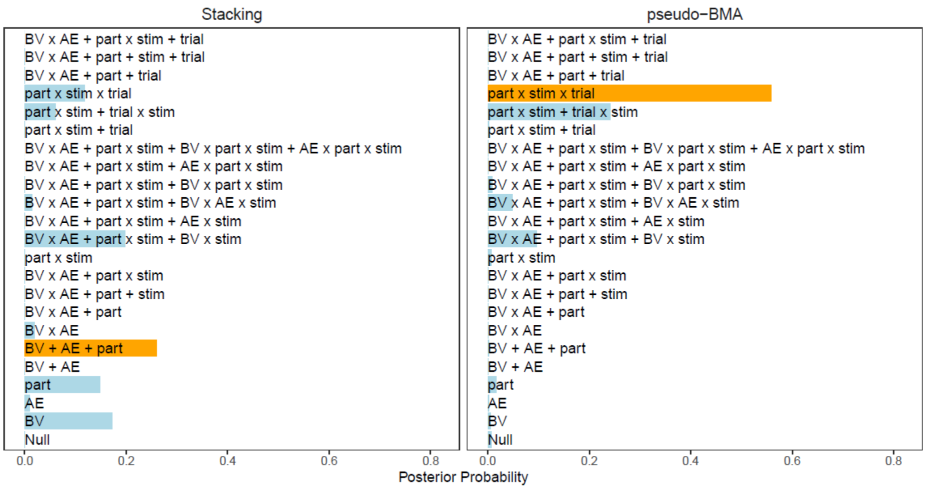


#### L. VC Automatically Constrained Thought

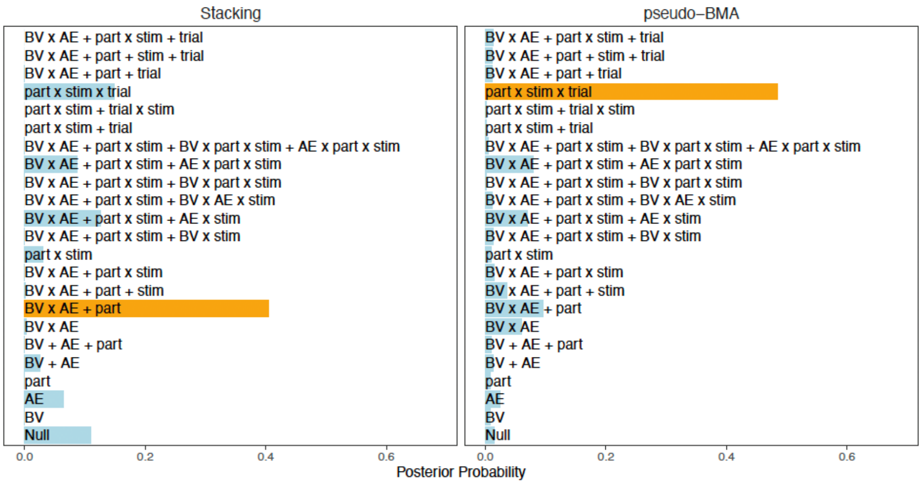

**Figure S2. Winning Model Predictors**

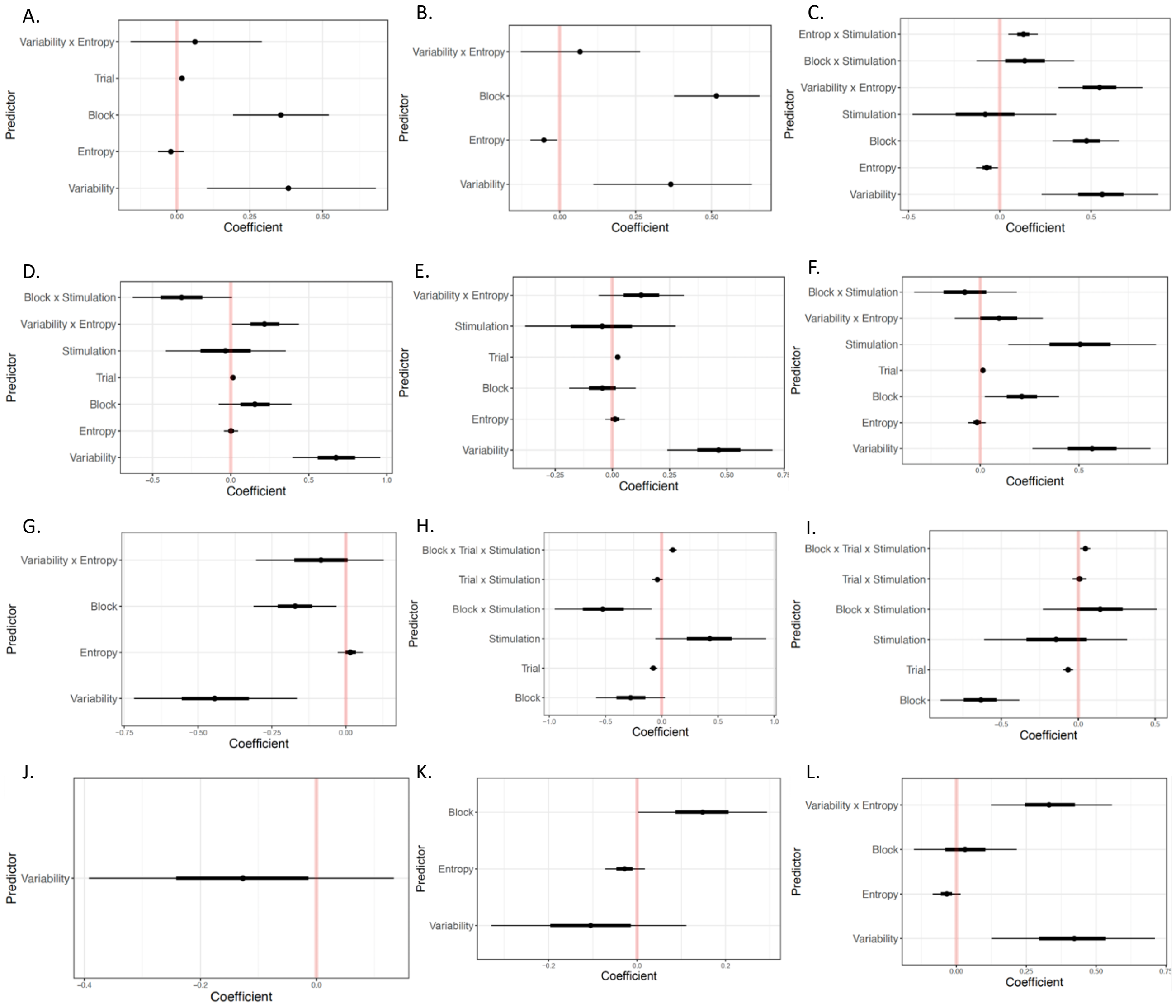


*Note*. This figure represents the winning probit model for each region and thought type. A: PFC task unrelated thought. B: IPL task unrelated thought. C: VC task unrelated thought. D: PFC freely moving thought. E: IPL freely moving thought. F: VC freely moving thought. G: PFC deliberately constrained thought. H: IPL deliberately constrained thought. I: VC deliberately constrained thought. J: PFC automatically constrained thought. K: IPL automatically constrained thought. L: VC automatically constrained thought.

#### Table S1. Selected model-coefficients

| Thought type | PFC | | IPL | | VC | |
|---|---|---|---|---|---|---|
| | LOOIC | Pseudo-BMA | LOOIC | Pseudo-BMA | LOOIC | Pseudo-BMA |
| **Task unrelated thought** | Mod20 - 0.24<br>Mod19 - 0.20 | Mod19 - 0.92<br>Mod18 - 0.03 | Mod07 - 0.24<br>Mod19 - 0.23 | Mod19 - 1.00<br>Mod21 - 0.00 | Mod12 - 0.25<br>Mod19 - 0.18 | Mod19 - 0.93<br>Mod20 - 0.03 |
| **Freely moving thought** | Mod22 - 0.49<br>Mod19 - 0.16 | Mod19 - 0.72<br>Mod21 - 0.10 | Mod21 - 0.34<br>Mod19 - 0.19 | Mod19 - 0.98<br>Mod21 - 0.01 | Mod22 - 0.32<br>Mod20 - 0.28 | Mod19 - 0.57<br>Mod20 - 0.21 |
| **Deliberately constrained thought** | Mod07 - 0.20<br>Mod04 - 0.19 | Mod19 - 0.57<br>Mod17 - 0.29 | Mod19 - 0.38<br>Mod21 - 0.18 | Mod19 - 0.98<br>Mod21 - 0.01 | Mod19 - 0.25<br>Mod07 - 0.19 | Mod19 - 0.57<br>Mod20 - 0.13 |
| **Automatically constrained thought** | Mod01 - 0.18<br>Mod17 - 0.14 | Mod17 - 0.23<br>Mod00 - 0.19 | Mod05 - 0.26<br>Mod11 - 0.20 | Mod19 - 0.56<br>Mod18 - 0.24 | Mod07 - 0.40<br>Mod19 - 0.15 | Mod19 - 0.49<br>Mod07 - 0.10 |

*Note.* The top two models selected for both the LOOIC and Pseudo-BMA procedures for each thought type and region.

#### Table S2. Blinding analyses

A. Task unrelated thought objective and subjective stimulation effects

| Region | | Active | | Sham | | $BF_{10}$ | $BF_{incl}$ | $F$ | $p$ | $\eta_p^2$ |
|---|---|---|---|---|---|---|---|---|---|---|
| | | *M* (SD) | *n* | *M* (SD) | *n* | | | | | |
| **PFC** | Objective | 4.11 (1.51) | 38 | 4.51 (1.51) | 38 | 0.43 | 0.43 | 0.65 | .423 | 0.009 |
| | Subjective | 4.36 (1.54) | 65 | 4.04 (1.37) | 11 | 0.37 | 0.37 | 0.466 | .4977 | 0.006 |
| | Objective * subjective | | | | | | 0.48 | <0.01 | .971 | 1.789e-5 |
| **IPL** | Objective | 4.55 (1.25) | 39 | 4.31 (1.59) | 37 | 0.30 | 0.30 | 0.03 | .862 | 4.218e-4 |
| | Subjective | 4.42 (1.34) | 67 | 4.54 (2.00) | 9 | 0.35 | 0.34 | 0.07 | .797 | 9.242e-4 |
| | Objective * subjective | | | | | | 0.45 | 0.15 | .704 | 0.002 |
| **VC** | Objective | 4.54 (1.52) | 37 | 4.40 (1.34) | 39 | 0.26 | 0.26 | 1.86 | .177 | 0.025 |
| | Subjective | 4.38 (1.41) | 65 | 5.00 (1.48) | 11 | 0.64 | 0.64 | 2.16 | .146 | 0.029 |
| | Objective * subjective | | | | | | 0.68 | 2.10 | .151 | 0.028 |

B. Freely moving thought objective and subjective stimulation effects

| Region | | Active | | Sham | | BF$_{10}$ | BF$_{incl}$ | *F* | *p* | $\eta_p^2$ |
|---|---|---|---|---|---|---|---|---|---|---|
| | | *M* (SD) | *n* | *M* (SD) | *n* | | | | | |
| PFC | Objective | 3.51 (1.40) | 38 | 4.08 (1.33) | 38 | 1.01 | 1.00 | 0.75 | .390 | 0.010 |
| | Subjective | 3.73 (1.35) | 65 | 4.17 (1.63) | 11 | 0.46 | 0.45 | 0.90 | .347 | 0.012 |
| | Objective * subjective | | | | | | 0.48 | 0.31 | .582 | 0.004 |
| IPL | Objective | 3.99 (1.09) | 39 | 4.01 (1.33) | 37 | 1.88 | 0.26 | 1.13 | .291 | 0.015 |
| | Subjective | 3.93 (1.16) | 67 | 4.82 (1.31) | 9 | 0.25 | 1.89 | 4.93 | .030 | 0.064 |
| | Objective * subjective | | | | | | 0.50 | 1.02 | .317 | 0.014 |
| VC | Objective | 4.11 (1.25) | 37 | 3.46 (1.32) | 39 | 1.89 | 1.88 | 5.25 | .025 | 0.068 |
| | Subjective | 3.80 (1.31) | 65 | 3.66 (1.42) | 11 | 0.61 | 0.33 | 0.04 | .842 | 5.565e-4 |
| | Objective * subjective | | | | | | 0.47 | 1.12 | .293 | 0.015 |

C. Deliberately constrained thought objective and subjective stimulation effects

| Region | | Active | | Sham | | BF$_{10}$ | BF$_{incl}$ | *F* | *p* | $\eta_p^2$ |
|---|---|---|---|---|---|---|---|---|---|---|
| | | *M* (SD) | *n* | *M* (SD) | *n* | | | | | |
| PFC | Objective | 3.89 (1.42) | 38 | 3.61 (1.36) | 38 | 0.34 | 0.33 | 0.17 | .680 | 0.002 |
| | Subjective | 3.84 (1.37) | 65 | 3.21 (1.46) | 11 | 0.68 | 0.67 | 1.88 | .175 | 0.025 |
| | Objective * subjective | | | | | | 0.42 | 0.06 | .805 | 8.519e-4 |
| IPL | Objective | 3.72 (1.38) | 39 | 3.81 (1.50) | 37 | 0.25 | 0.24 | 0.97 | .327 | 0.013 |
| | Subjective | 3.82 (1.38) | 67 | 3.35 (1.78) | 9 | 0.47 | 0.47 | 0.64 | .425 | 0.009 |
| | Objective * subjective | | | | | | 0.59 | 1.20 | .277 | 0.016 |
| VC | Objective | 3.37 (1.24) | 37 | 3.63 (1.46) | 39 | 0.33 | 0.33 | 1.45 | .233 | 0.020 |
| | Subjective | 3.42 (1.33) | 65 | 3.99 (1.46) | 11 | 0.61 | 0.61 | 1.42 | .237 | 0.019 |
| | Objective * subjective | | | | | | 0.44 | 0.78 | .380 | 0.011 |

D. Automatically constrained thought objective and subjective stimulation effects

| Region | | Active | | Sham | | BF$_{10}$ | BF$_{incl}$ | *F* | *p* | $\eta_p^2$ |
|---|---|---|---|---|---|---|---|---|---|---|
| | | *M* (SD) | *n* | *M* (SD) | *n* | | | | | |
| PFC | Objective | 1.98 (0.91) | 38 | 2.16 (1.06) | 38 | 0.31 | 0.30 | 0.22 | .637 | 0.003 |
| | Subjective | 2.04 (0.96) | 65 | 2.24 (1.17) | 11 | 0.36 | 0.36 | 0.47 | .498 | 0.006 |
| | Objective * subjective | | | | | | 0.83 | 1.97 | .165 | 0.027 |
| IPL | Objective | 2.31 (0.91) | 39 | 2.13 (0.91) | 37 | 0.33 | 0.34 | 0.12 | .727 | 0.002 |
| | Subjective | 2.26 (0.93) | 67 | 1.94 (0.73) | 9 | 0.49 | 0.49 | 0.94 | .335 | 0.013 |
| | Objective * subjective | | | | | | 0.43 | 0.10 | .758 | 0.001 |

| | | | | | | | | | | |
|---|---|---|---|---|---|---|---|---|---|---|
| | Objective | 2.54 (1.15) | 37 | 2.16 (1.08) | 39 | 0.64 | 0.661 | 2.33 | .131 | 0.031 |
| **VC** | Subjective | 2.27 (0.95) | 65 | 2.78 (1.84) | 11 | 0.68 | 0.699 | 2.23 | .140 | 0.030 |
| | Objective * subjective | | | | | | 0.407 | 0.37 | .548 | 0.005 |

### Figure S3. Stimulation Block Model Comparisons

A. PFC Task Unrelated Thought

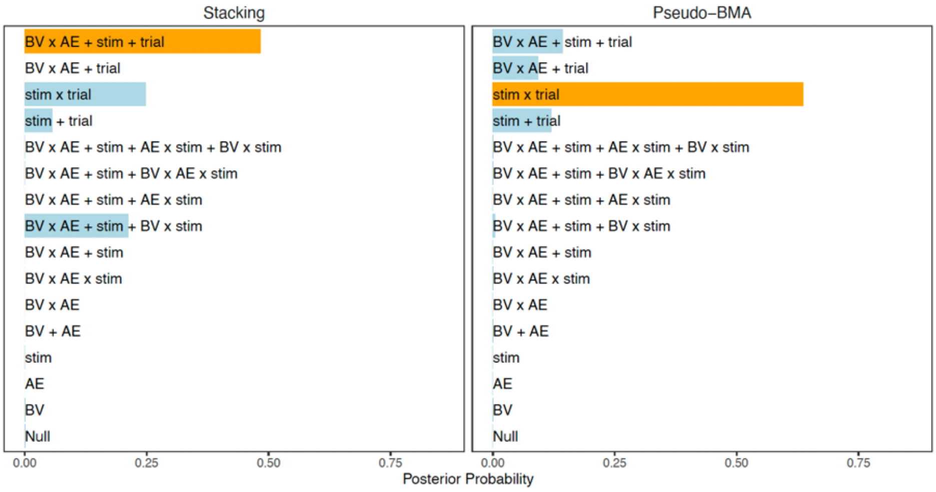


B. IPL Task Unrelated Thought

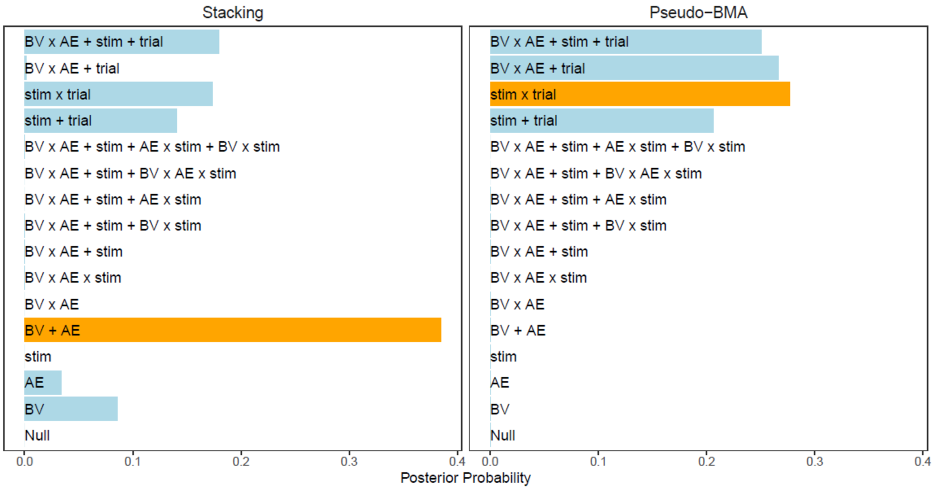

C. VC Task Unrelated Thought

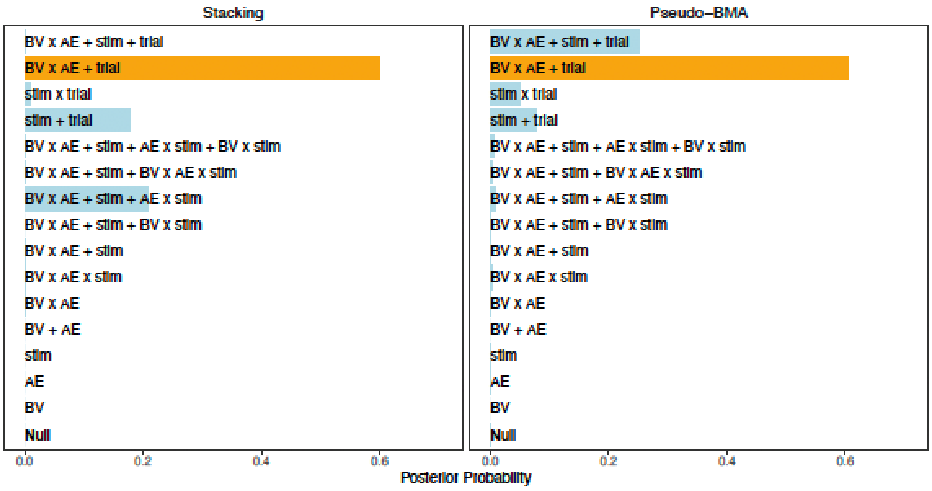


D. PFC Freely Moving Thought

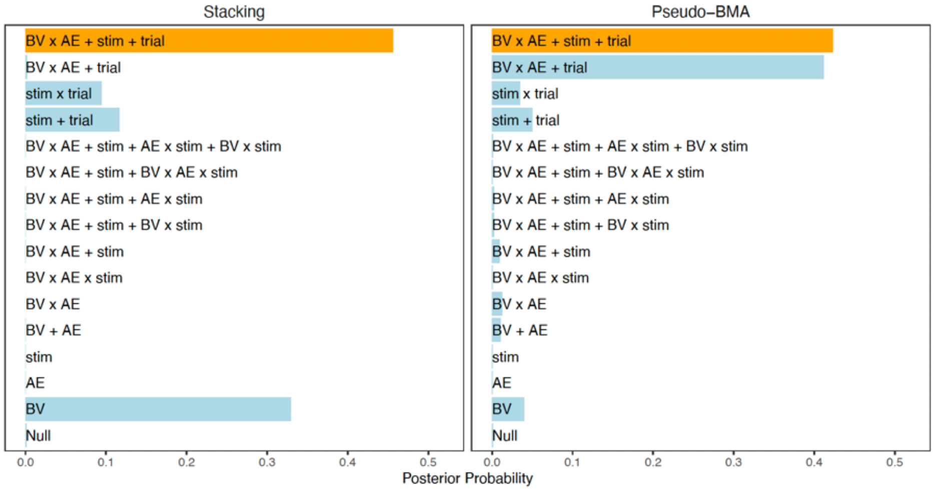


E. IPL Freely Moving Thought

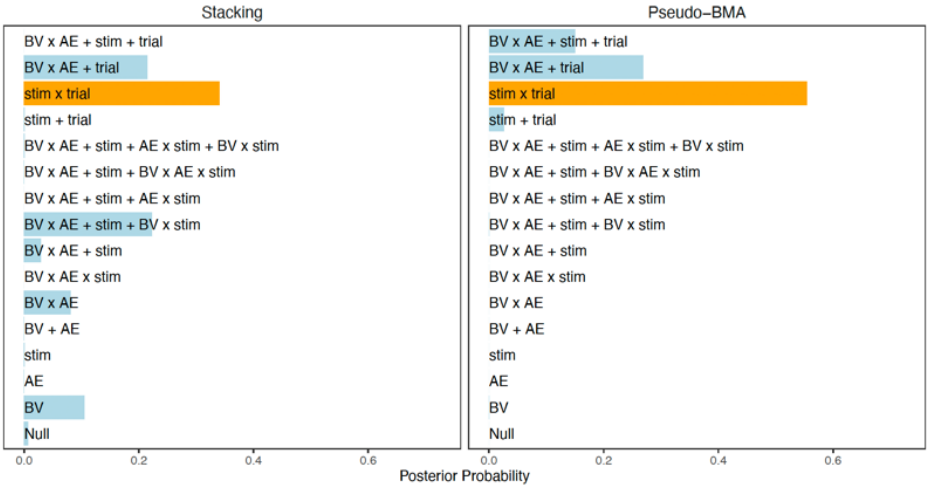

#### F. VC Freely Moving Thought

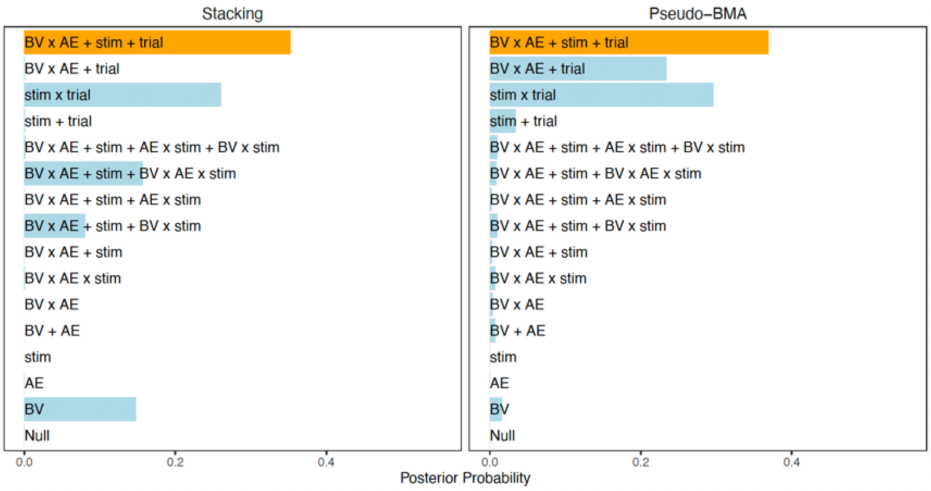


#### G. PFC Deliberately Constrained Thought

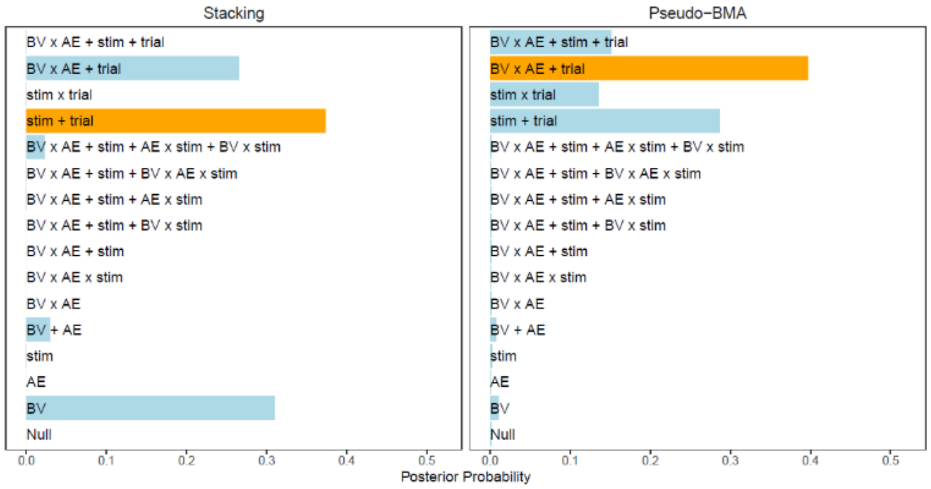


#### H. PFC Automatically Constrained Thought

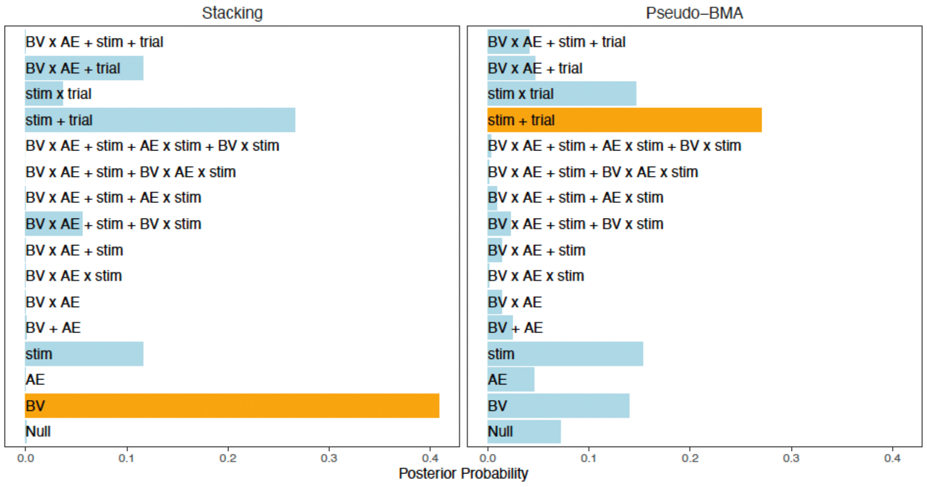

I. IPL Automatically Constrained Thought

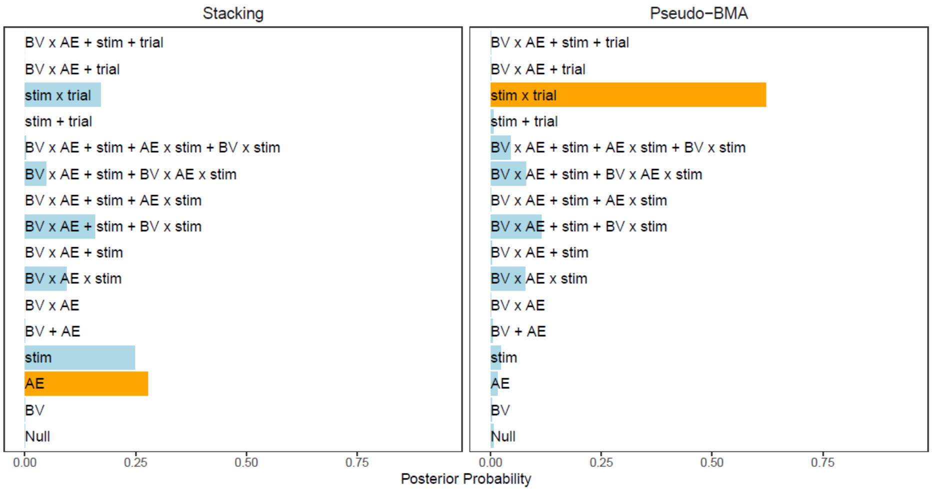


J. VC Automatically Constrained Thought

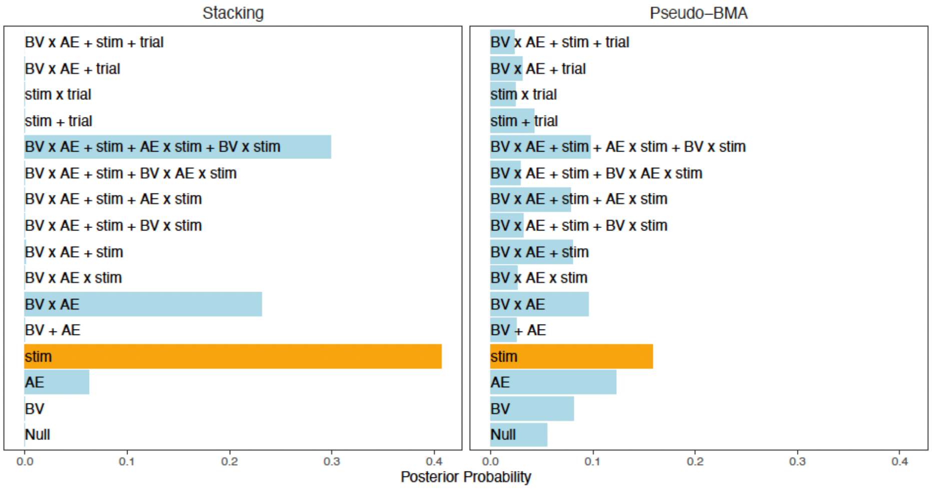

#### Figure S4. Stimulation Block Winning Model Predictors

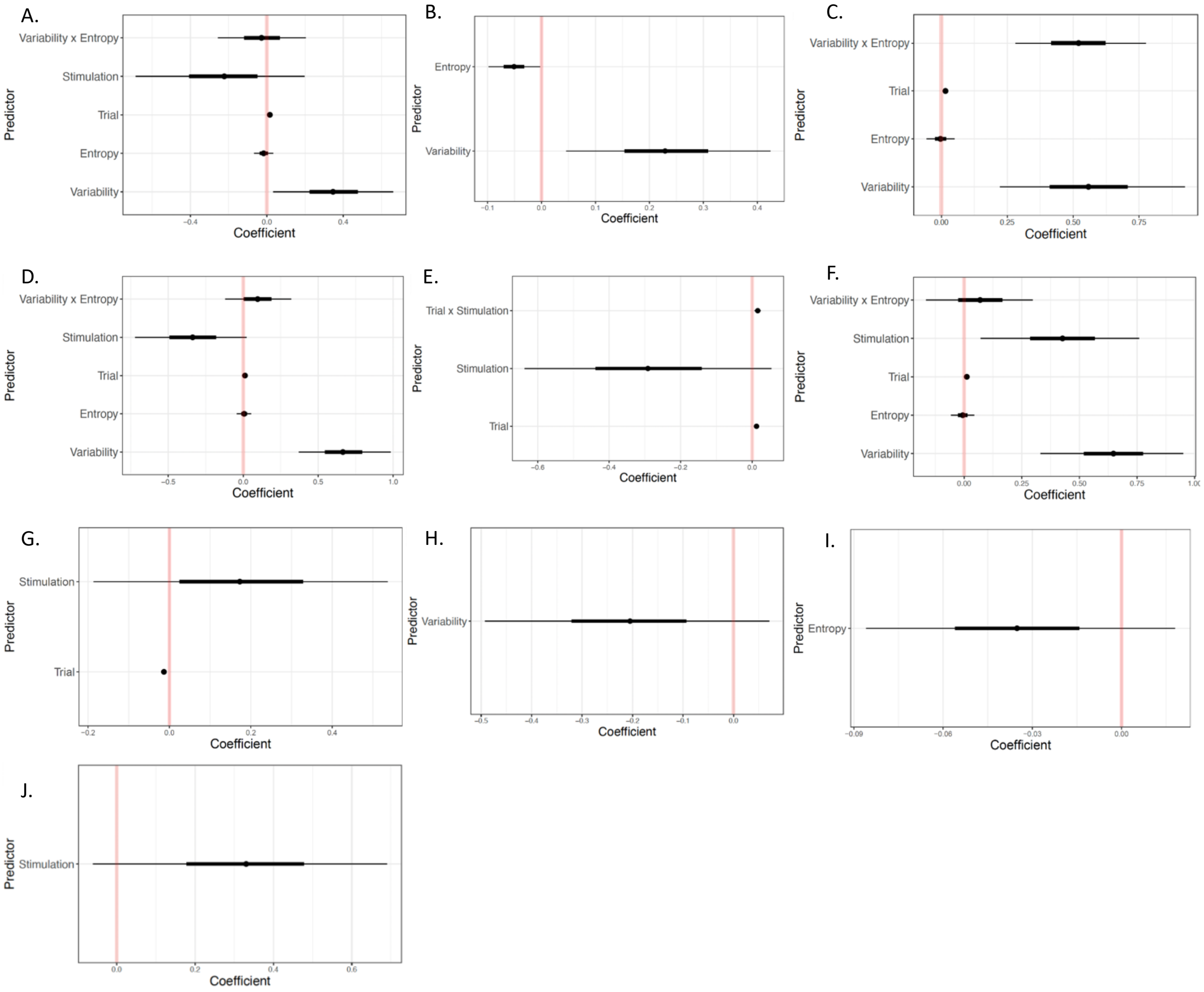


*Note.* This figure represents the winning probit model for each region and thought type that had conflicting winning models in the full model comparisons. A: PFC task unrelated thought. B: IPL task unrelated thought. C: VC task unrelated thought. D: PFC freely moving thought. E: IPL freely moving thought. F: VC freely moving thought. G: PFC deliberately constrained thought. H: PFC automatically constrained thought. I: IPL automatically constrained thought. J: VC automatically constrained thought.

#### Table S3. Selected model-coefficients

| Thought type | PFC | | IPL | | VC | |
|---|---|---|---|---|---|---|
| | LOOIC | Pseudo-BMA | LOOIC | Pseudo-BMA | LOOIC | Pseudo-BMA |
| **Task unrelated thought** | Mod15 – 0.48<br>Mod13 – 0.25 | Mod13 – 0.64<br>Mod15 – 0.14 | Mod04 – 0.39<br>Mod15 – 0.18 | Mod13 – 0.28<br>Mod14 – 0.27 | Mod14 – 0.60<br>Mod09 – 0.21 | Mod14 – 0.60<br>Mod15 – 0.25 |
| **Freely moving thought** | Mod15 – 0.46<br>Mod01 – 0.33 | Mod15 – 0.42<br>Mod14 – 0.41 | Mod13 – 0.34<br>Mod08 – 0.22 | Mod13 – 0.55<br>Mod14 – 0.27 | Mod15 – 0.35<br>Mod13 – 0.26 | Mod15 – 0.37<br>Mod13 – 0.30 |
| **Deliberately constrained thought** | Mod12 – 0.37<br>Mod01 – 0.31 | Mod14 – 0.40<br>Mod12 – 0.29 | | | | |
| **Automatically constrained thought** | Mod01 – 0.41<br>Mod12 – 0.27 | Mod12 – 0.27<br>Mod03 – 0.15 | Mod02 – 0.28<br>Mod03 – 0.25 | Mod13 – 0.62<br>Mod08 – 0.11 | Mod03 – 0.41<br>Mod11 – 0.30 | Mod03 – 0.16<br>Mod02 – 0.12 |

*Note.* The top two models selected for both the LOOIC and Pseudo-BMA procedures for each thought type and region in the stimulation block.
